# Elucidation of the essentiality of lumazine synthase (RibH) for *Mycobacterium tuberculosis* survival and discovery of potent inhibitors for enhanced antimycobacterial therapy

**DOI:** 10.1101/2023.07.18.549608

**Authors:** Monica Singh, Anannya Dhanwal, Arpita Verma, Linus Augustin, Niti Kumari, Soumyananda Chakraborti, Nisheeth Agarwal, Dharmarajan Sriram, Ruchi Jain Dey

## Abstract

Tuberculosis (TB) continues to be a global health crisis, necessitating urgent interventions to address drug resistance and improve treatment efficacy. In this study, we validate the indispensable role of lumazine synthase (RibH), a vital enzyme in the riboflavin biosynthetic pathway, in the survival of *Mycobacterium tuberculosis (M. tb)* using a CRISPRi-based conditional gene knockdown strategy. We show that genetic and functional ablation of RibH in *M. tb* cannot be compensated by exogenous supply of Riboflavin, or co-factors Flavin Adenine Dinucleotide (FAD) or Flavin Mononucleotide (FMN). Capitalizing on the essentiality of RibH, we employ a high-throughput molecular docking approach to screen ∼600,000 compounds and identify inhibitors of RibH. Through *in vitro* screening of 55 shortlisted compounds, we discover 3 inhibitors that exhibit potent antimycobacterial activity. These compounds effectively also eradicate intracellular *M. tb* during macrophage infection and prevent the resuscitation of the nutrient-starved persister bacteria. Moreover, these 3 compounds synergistically enhance the bactericidal effect of first-line anti-TB drugs, Isoniazid and Rifampicin. Corroborating with the *in silico* predicted high docking scores along with favorable ADME and toxicity profiles, all 3 compounds demonstrate exceptional binding affinity towards purified lumazine synthase enzyme *in vitro*, and display an acceptable safety profile in mammalian cells, with a high selective index. By providing mechanistic evidence for the essentiality of RibH in *M. tb* survival, and discovering potent RibH inhibitors with outstanding antimycobacterial activity, our study contributes to the development of superior TB treatment strategies and advances the global fight against this devastating disease.

## Introduction

Tuberculosis (TB) remains a leading cause of death globally^1, 2^. The current standard TB treatment regimen involves a combination of multiple drugs taken over a long duration of 6 to 9 months^1, 2^. The lengthy treatment duration escalates the risk of patient non-compliance, leading to treatment failure and the development of drug resistance^1^. The mounting challenges with current anti-TB therapies have hindered the efforts to eliminate TB by 2030 and have created an urgent demand for the development of novel and impactful antibiotics^2^. *Mycobacterium tuberculosis* (*M. tb*) is highly vulnerable to developing resistance mutations against existing drugs ^2, 3^, as the majority of these drugs have one or few target(s) which allow pathogens to evolve mechanisms for evading or offsetting the inhibitory effects ^3, 4^. Hence, the development of drugs targeting new pathways is considered one of the important strategies to circumvent the current drug resistance mechanisms^5, 6^. It unlocks the prospect of combining them with existing first-line anti-TB drugs, resulting in synergistic effects with enhanced treatment outcomes, thus allowing for shorter, more efficacious regimens with lesser side effects and is more convenient to administer^7^.

Discovering new antibiotics for TB utilizes two strategic approaches: target-based (gene-to-drug model) and drug-based (drug-to-gene model) approaches^5, 7^. Availability of *M. tb* genome sequence^8^, and the advent of various genome engineering tools including recent-most CRISPR-interference (CRISPRi)^9–13^ approach have truly revolutionized the field bridging the divide between genomics and TB drug discovery aiding a better understanding of the target vulnerabilities. In this study, we focused on the riboflavin biosynthetic pathway of *M. tb* that comprises of seven genes^14^, out of which, four have been reported essential for *in vitro* and *in vivo* survival of the pathogen, namely, *ribA2* (Rv1415), *ribG* (Rv1409), *ribH* (Rv1416) and *ribF* (Rv2786c) by transposon site hybridization (TraSH) screen ^12–14^. We have earlier shown that over-producing these essential genes reduce mycobacterial virulence by triggering a greater mucosal-associated invariant T cell response^14^. This pathway produces essential redox cofactors, flavin mononucleotide (FMN) and Flavin adenine dinucleotide (FAD), which are required for > 3% of the *M. tb* proteins’ acivity^15^. Recently, using a systems biology approach, Beste et al, predicted that FAD is a crucial cofactor with a significant role in respiratory metabolism, nucleotide biosynthesis, and amino acid metabolism^16, 17^. FAD is also essential for the beta-oxidation of fatty acids, which is a crucial cofactor for mycolic acid and cell wall lipid biosynthesis and for metabolizing host-derived lipids^16^. Out of the 250 genes involved in fatty acid metabolism ^18^, *in silico* studies predict that ∼70 mycobacterial gene products rely on FAD as a cofactor^16^. To explore the therapeutic potential of the riboflavin biosynthesis pathway, multiple attempts were made in the early 2000s ^19–25^, towards the structural analysis of RibH (lumazine synthase) and other enzymes of this pathway^26^ and identification of their inhibitors. However, the antimycobacterial potential of these compounds is yet to be explored. A recent study reported designing riboflavin analogs targeting FMN riboswitch, showing inhibitory activity against *M. tb*^19^.

We hypothesize that the riboflavin pathway could be a potential Achilles’ heel and proteins involved in this pathway could be potential drug targets. Among the various enzymes, RibH shows a high level of expression in the non-replicating persistence stage of *M. tb*^27^ and is essential for *in vitro* and *in vivo* growth. These observations suggest a potential role of this enzyme in actively replicating as well as non-replicating dormant mycobacteria. In this study, we establish the indispensability of RibH for *M. tb*. viability and identify potential compounds capable of binding to the enzyme from a library of ∼600,000 molecules. We report three highly potent compounds that exhibit promising inhibitory effects against *M. tb H37Rv* when tested as standalone drugs or in combination with first-line anti-TB drugs leading to a substantial *en bloc* enhancement in the anti-mycobacterial activity of rifampicin and isoniazid. Further, we report that all three inhibitors are non-toxic against human cells and possess a high selective index. Importantly, these molecules are highly effective against intracellular as well as non-replicating dormant bacteria.

## Results

### Conditional silencing by CRISPRi reveals the role of lumazine synthase in mycobacterial growth

To determine the requirement of lumazine synthase encoded by *ribH* (*Rv1416*) for mycobacterial growth, we utilized the CRISPRi method to create knockdown strains utilizing *M. tb H37Rv* mc^2^ 7902^28^. The *M. tb* knockdown strain is conditionally depleted for *ribH* (annotated as *ribH*-KD) in an anhydrotetracycline (ATc)– dependent manner, as described earlier and in methods^29, 30^. The ATc-dependent induction of dCas9 results in the silencing of *ribH* in a highly specific manner guided by gene-specific guide RNAs encoded on a separate plasmid **(Supplementary Figure 1A, B)**. Treatment of *ribH-*KD strain with 50 and 100 ng/ml ATc for 9 days in broth results in visible growth defects as evidenced by reduced cell pellet size (**Figure 1A**) and reduced growth on 7H11 pantothenate, leucine, and arginine (PLA)-supplemented agar plates (**Figure 1B**). The impact of ATc induction on the growth of *ribH*-KD strain of *M. tb* is also evident from the growth curves assessed in the presence of ATc for 9 days (**Figure 1C**). Importantly, growth defects start appearing from day 4 onwards, causing ∼27% and ∼40% reduction in OD_600_ following incubation with 50 and 100 ng/ml of ATc, respectively. The growth defects are further accentuated by day 7, wherein a marked reduction of ∼54% and ∼64% was observed with 50 and 100 ng/ml of ATc, respectively, in comparison to the parent strain (**Figure 1C**). These differences are found to be statistically significant at 50 ng/ml (*p* < 0.05) and 100 ng/ml (*p* < 0.0001). However, *ribH*-KD strain of *M. tb* without ATc treatment shows a similar growth profile as that of the parent strain.

**Figure 1.**
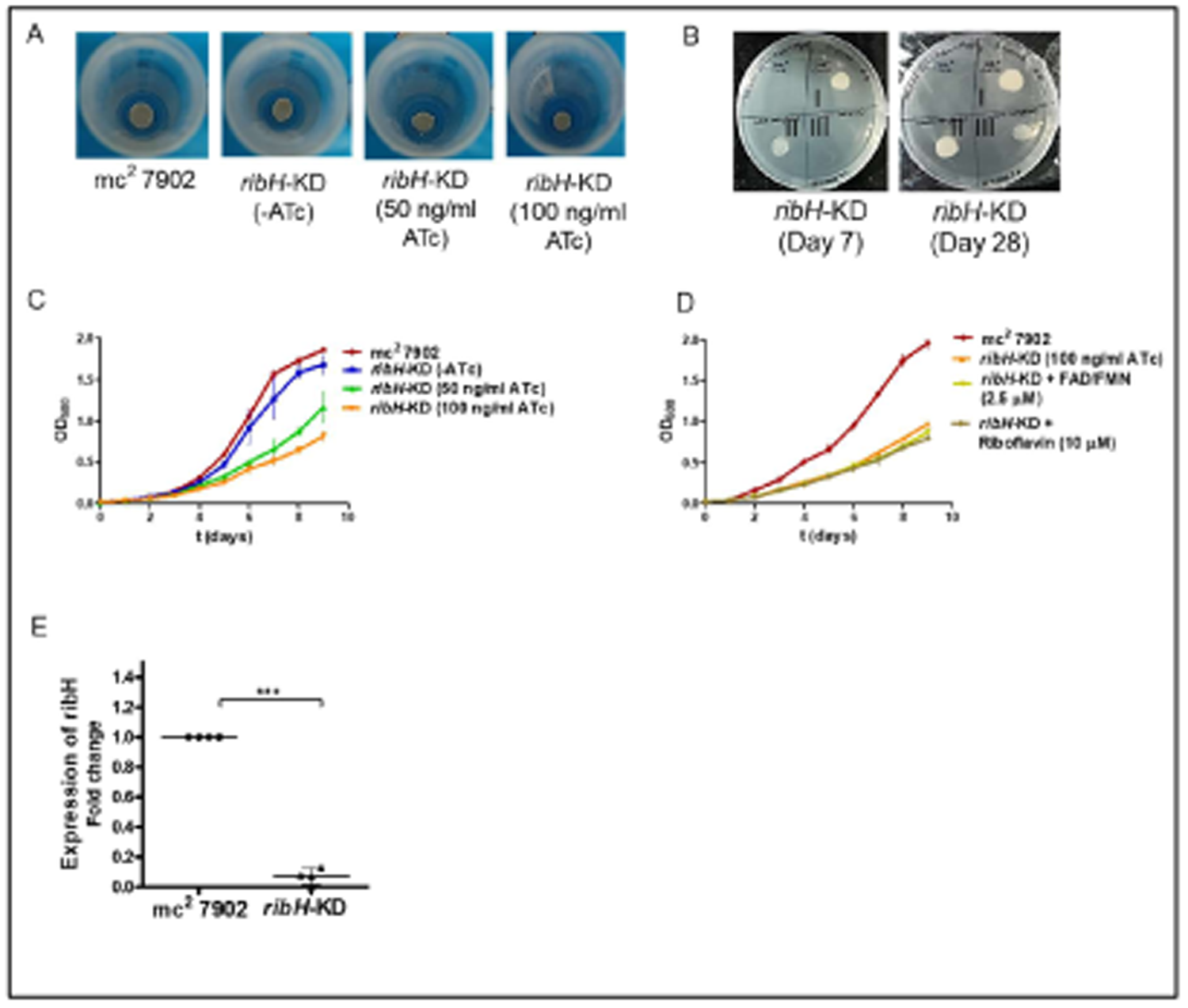
Growth characteristics of conditionally silenced *ribH*-KD *M. tb*. A. Image showing growth defects in the *ribH*-KD strain of *M. tb mc^2^ 7902* as evident from reduced pellet size after 9 days of treatment with 50 and 100 ng/ml of ATc. B. Image showing delayed *in vitro* growth of *ribH*-KD strain spotted on 7H11 agar plates after 9 days of treatment with 0 ATc (I), 50 ng/ml ATc (II) and 100 ng/ml ATc (III). *ribH*-KD shows delayed and reduced growth when monitored at 7 and 28 days of inoculation. C. Time-stamped growth curves of parent mc^2^ 7902 and *ribH-*KD at different concentrations of ATc (0, 50 and 100 ng/ml) at 37_°_C, 200 rpm. OD_600_ was recorded at regular intervals up to 9 days. Graph is plotted using means ± s.d of two independent experiments. D. Time-stamped growth curves of parent mc^2^ 7902 and *ribH-*KD at 100 ng/ml ATc in chemically defined synthetic media with riboflavin (10μM) or cocktail of FMN and FAD (2.5μM each) at 37_°_C, 200 rpm. OD_600_ was recorded at regular intervals up to 9 days. Graph is plotted using means ± s.d of two independent experiments. E. Quantitative RT-PCR to validate CRISPRi-mediated silencing of *ribH*. RNA samples were isolated after 9 days of treatment with 100 ng/ml ATc and RT-PCR analysis was carried out showing reduced levels of *ribH* transcripts in *ribH*-KD in comparison to the parent *M. tb mc^2^ 7902*. Fold change data (2^-ddCT^; n = 4) is plotted with respect to the parent strain. Fold change value is represented as mean ± s.d and statistical significance was measured using Student’s t-test, ***, *p < 0.0001*.

We next assessed if supplementation with riboflavin (10µM) or a cocktail of FMN and FAD (2.5 µM each) in the media compensates for the growth defects observed in the case of *ribH*-KD on ATc induction. Strikingly, exogenous supplementation fails to resurrect the growth defects of this knockdown strain (**Figure 1D**), suggesting a possible inability of *M. tb* to procure riboflavin or readymade FAD and FMN from the external environment, further strengthening the essentiality of riboflavin biosynthesis for mycobacterial survival.

The growth defect is accompanied by a significant reduction (92%; *p*< 0.0001) in the expression of *ribH* gene in *ribH-KD* following 9 days of treatment with 100 ng/ml ATc (**Figure 1E**). At 50 ng/ml ATc, we observe 21-44% reduced *ribH* gene expression **(Supplementary Figure 1C)**. However, there is no impact of the dCas9-based knockdown of *ribH* on the general transcription capability of *M. tb* as signified by the unaltered expression of the internal control gene, sigma factor A (*sigA; Rv 2703)*. As a more consistent and pronounced reduction in gene expression of *ribH* and growth defects is observed at 100 ng/ml ATc, the same concentration was employed for all the follow-up experiments.

Interestingly, when the 10-fold serial dilutions of *ribH*-KD cultures from the above experiments were spotted on the 7H11 agar plates following 9 days of ATc treatment and allowed to grow for 3-4 weeks, we observed a striking difference in the colony morphology of the *ribH*-KD strain compared to the parent strain (**Figure 2A**). The colonies of *ribH*-KD are found to be relatively thinner, shrunken, dry, and raised in appearance compared to the parent strain. The bacilli also appear to be arranged more compactly in these colonies compared to the arrangement in the parent colonies. Supplementation of *ribH*-KD with FAD, FMN or riboflavin cannot recover the original colony morphology. Hence, we next assessed if the reduction in *ribH* gene expression impacts cell length and thickness (or width) by scanning electron microscopy (SEM). Strikingly, *ribH*-KD bacilli (∼ 2.377 ± 0.06378 µm; n=61) is found to be significantly longer (∼1.5 - fold, *p* < 0.0001) than the parent *M. tb* strain (1.593 ± 0.04382 µm; n=61) (**Figure 2B, C)**. However, no evident changes are observed in the thickness (or width) of *ribH*-KD strain (**Figure 2C)**. Altogether, *ribH* is found to play a key role in regulating mycobacterial replication, colony, and bacterial morphology.

**Figure 2.**
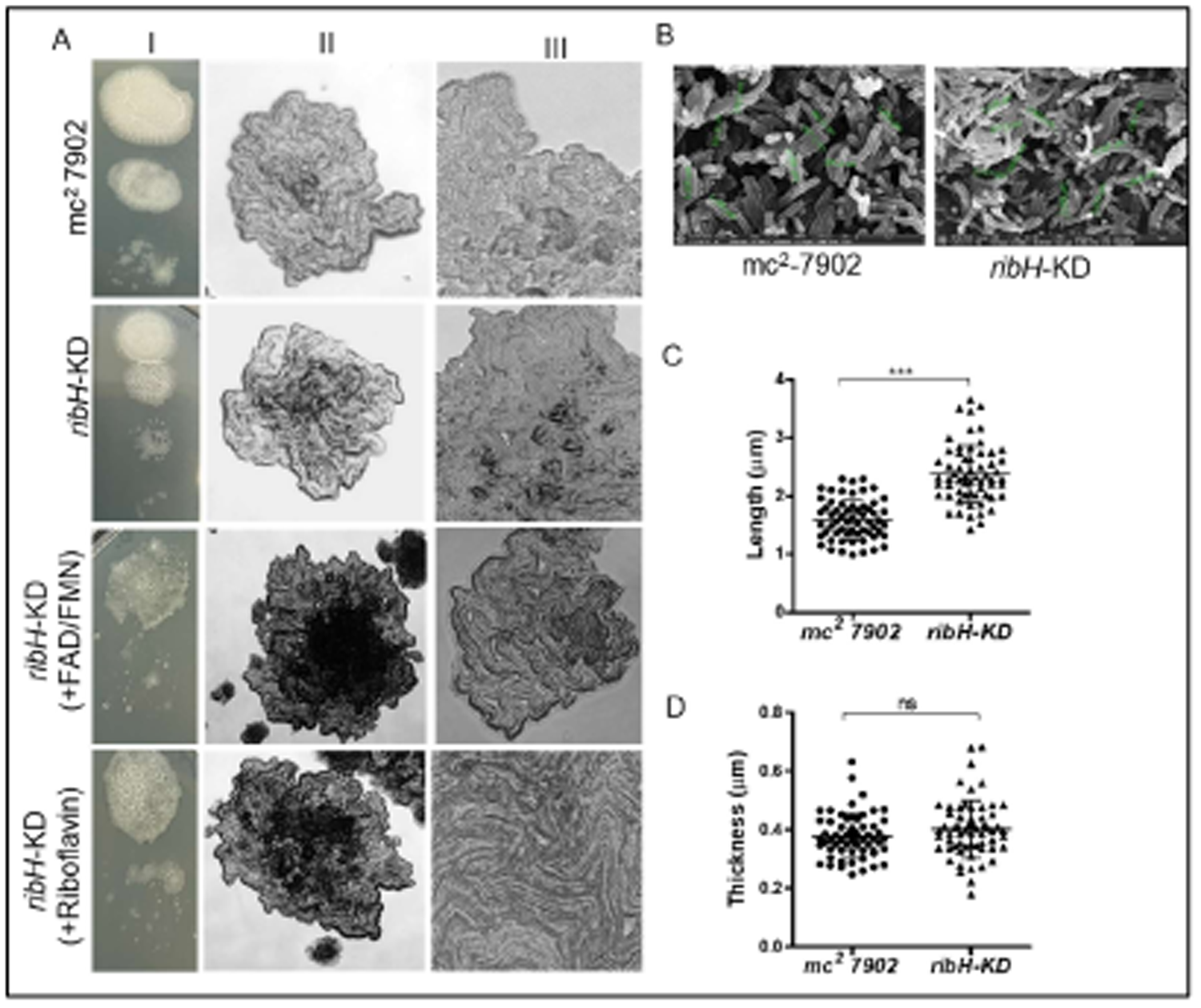
Altered colony morphology and bacillary length in *ribH-KD*. A.(I) Visual illustration of the reduced growth after 9 days of conditional knockdown with 100 ng/ml ATc in broth followed by 2 weeks of inoculation on agar plates. A (II, III). Micrographs showing altered colony morphology of *ribH-* KD strain in the absence or presence of riboflavin and cocktail of FAD and FMN with respect to the parent *mc^2^ 7902* strain when measured at 4x (column II) and 20x (column III), respectively. B. Reduced bacillary length of *ribH*-KD strain with respect to the parent strain. At day-5 post treatment with 100 ng/ml ATc, the *ribH*-KD strain and its parental *mc^2^ 7902* strain were assessed for their bacillary length and thickness (width) using scanning electron microscopy (SEM), Bars-3μm. C. Bacillary length (μm) and thickness (μm) were measured using ImageJ software and data (n = 61) was plotted as mean ± s.d in form of dot plots. Each data point corresponds to a single measurement of a bacillus. Data was analyzed using Student’s t-test, the *p* values were determined, \*\*\**p* < 0.0001; ns: non-significant.

### *In silico* molecular docking to identify compounds that could potentially inhibit RibH

Taking cues from the growth defects observed on conditional knockdown of the *ribH* gene and the inability of *M. tb* to secure this nutrient from outside, we next assessed the prospects of designing drugs targeting RibH, lumazine synthase. For this, we utilized the crystal structure of *M. tb* lumazine synthase complexed with TP6 (PDB ID 2C92 of 1.6 Å resolution)^21^. Structurally, *M. tb* lumazine synthase exists as a homopentamer belonging to the α/β family of proteins. Each subunit of *M. tb* RibH consists of 160 amino acid residues forming a three-layer α/β/α architecture (**Figure 3A**) containing 6,7-dimethyl-8-ribityllumazine synthase (DMRL) domain (**Figure 3B**). As reported earlier, the crystal structure of *M. tb* lumazine synthase has active sites present at the interface of the two adjacent subunits, making a total of 5 substrate binding sites^31^ (**Figure 3C**). Visualization of the binding pocket using CB-Dock2 reveals a deep cavity of volume 889 Å^3^ consisting of twenty-nine amino acid residues viz., Ser 25, Trp 27, His 28, Leu 57, Gly 58, Ala 59, Ile 60, Glu 61, Val 81, Val 82, Ile 83, Arg 84, Gly 85, Gln 86, Thr 87, Pro 88, His 89, Phe 90, Val 93 of chain A and Ile 112, Ala 113, Asn 114, Gly 115, Arg 128, Glu 136, Lys 138, Gln 141, Ala 142 and Ala 145 of chain B. The dimension of the cavity along the x, y, and z axes is found to be 10, 14, and 16 Å, respectively (**Figure 3D**). According to the literature, lumazine synthase can bind riboflavin in its active sites^32, 33^. We re-docked the co-crystal ligands TP6 and riboflavin into the binding site and observed that these molecules are well fitted at the active site. Riboflavin penetrates deep inside into the *M. tb* RibH binding groove (**Figure 3D**), where it forms several hydrogen bonds with amino acid residues His 28, Ala 59, Glu 61, Val 81, Ile 83 of chain A, and Asn 114 and Lys 138 of chain B (**Figure 4A**).

**Figure 3.**
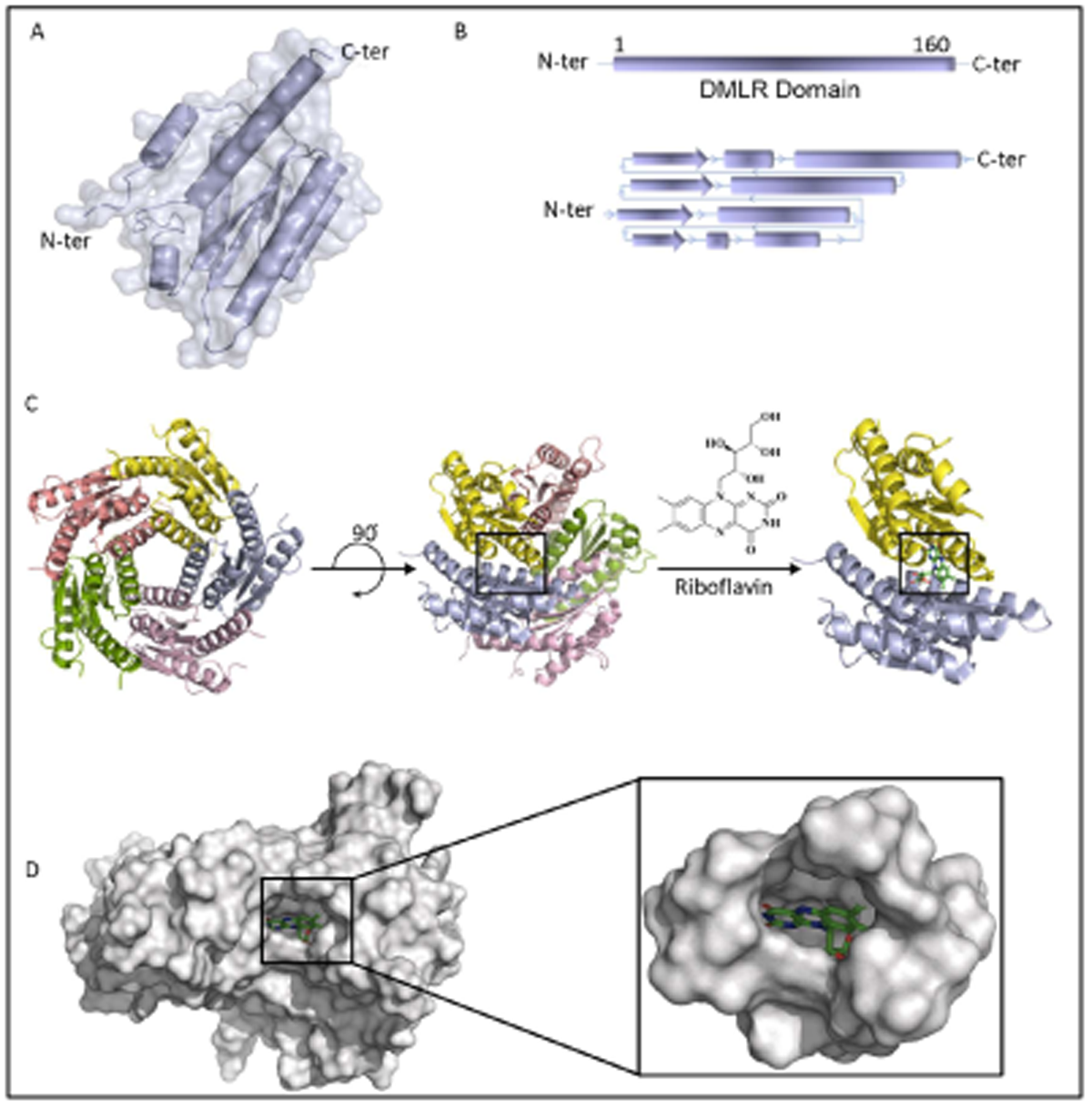
Structural characterization of *M. tuberculosis* RibH. A. Surface and cartoon representation of one subunit of *M. tb* RibH depicting the secondary structure elements from the N-terminus to the C-terminus, B. Schematic representation of the DMRL (6,7-dimethyl-8-ribityllumazine synthase) domain (upper panel) and secondary structure organization of *M. tb* RibH that belongs to α/β/α sandwich topology, where alpha helices are represented as pipes, beta strands are represented as arrows (bottom panel) and C. The homopentameric *M. tb* RibH (PDB ID 2C92) is shown as a cartoon with its active site present at the interface of two chains. Overall view of the *M. tb* RibH-riboflavin complex where the active site is occupied by riboflavin is shown, D. Surface representation of the *M. tb* RibH bound to riboflavin. Figure illustrates the substrate binding groove (cavity volume 889 Å^3^). The zoomed view shows the surface of the binding pocket with riboflavin buried inside.

**Figure 4.**
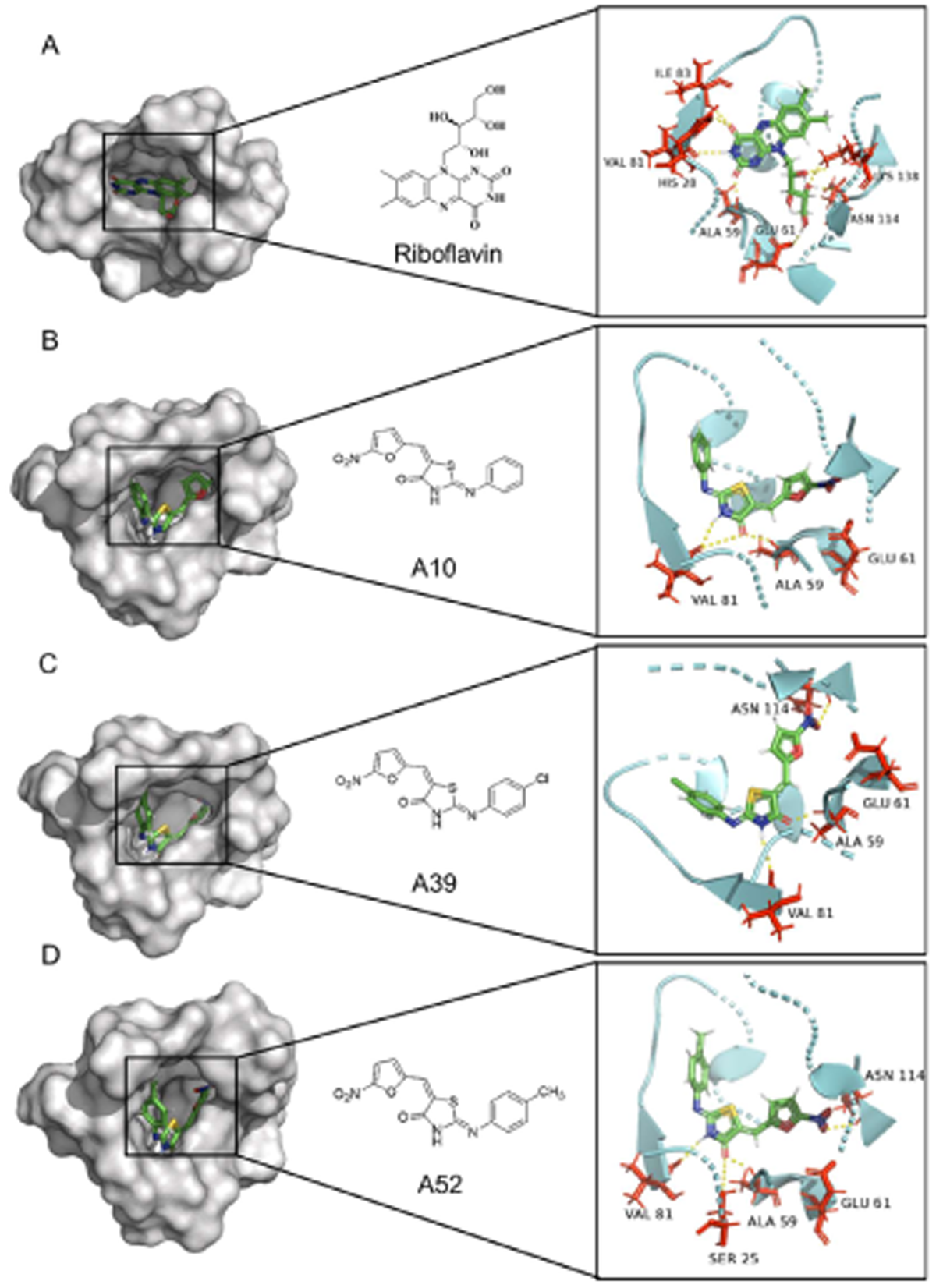
Active site surface representation and amino acid interactions of *M. tb* RibH with riboflavin and shortlisted drug molecules. Interaction of amino acids in the RibH active site are shown for A. Riboflavin, B. A10, C. A39 and D. A52. The zoomed view shows the bound fragment in the active pocket, wherein, A. Riboflavin interacts with His 28, Ala 59, Glu 61, Val 81, Ile 83, Asn 114 and Lys 138; B. A10 interacts with Ala 59, Glu 61 and Val 81; C. A39 interacts with Ala 59, Glu 61, Val 81 and Asn 114; D. A52 interacts with Ser 25, Ala 59, Glu 61, Val 81 and Asn 114. Hydrogen bonds are shown as yellow dashed lines.

We next docked our in-house library consisting of ∼ 3000 compounds and ∼0.57 million compounds from the ASINEX screening library (Asinex Corp.), to identify the leads with the highest docking scores and desirable absorption, distribution, metabolism, and excretion (ADME) properties as described in methods. On molecular docking, we identify fifty-five compounds (40 from our in-house library and 15 from the Asinex library) (as listed in **Supplementary Table 1**) with the best binding orientations, docking scores, and ADME values. These are also different from those reported earlier and hence, were selected for further assessment of antimycobacterial activity. As shown in **Supplementary Table 1**, we identified three compounds, namely A10 (5-((5-Nitrofuran-2-yl)methylene)-2-(phenylimino)thiazolidin-4-one), A39 (2-((4-Chlorophenyl)imino)-5-((5-nitrofuran-2-yl)methylene)thiazolidin-4-one) and A52 (5-((5-Nitrofuran-2-yl)methylene)-2-(*p*-tolylimino)thiazolidin-4-one) (**Figure 4B-D)** which exhibit the best minimum inhibitory concentration (MIC) among all the 55 compounds tested against *M. tb* strain *H37Rv* in the range of 0.78-1.56 µg/ml **(Supplementary Table 1)**. Hence, these three molecules were considered for further evaluation of their anti-TB activity by various biological assays. Strikingly, these compounds appear to be similar in their basic structure (see supplementary methods), with minor modifications in the R group resulting in different binding attributes (for synthesis scheme, see supplementary methods). The compounds namely, A10, A39, and A52, show a docking score of −7.276, −6.825, and −8.431 kcal/mol, respectively, signifying that these molecules can tightly bind to the RibH active site **(Supplementary Table 1)**.

As shown in **Figure 4B and Supplementary Table 1**, binding pocket analysis reveals the interaction of A10 with the RibH binding groove resulting in the formation of two hydrogen (H) bonds. The amide-carbonyl of A10 interacts with Ala 59 and the amide-NH interacts with Val 81 forming H-bonds of bond length 2.28 Å and 2.33 Å, respectively. In addition, the furan nitro (-NO2) group of A10 forms a salt bridge of bond length 3.79 Å with Glu 61 (**Figure 4B**). The furan nitro (-NO2) group of A39 interacts with Asn 114 of chain B forming an H-bond of length 2.46 Å, and the same group also interacts with Glu 61 of chain A forming a salt bridge of bond length 3.75 Å. Similar to A10, the amide-carbonyl and the amide-NH form H-bonds with Ala 59 and Val 81 of lengths 2.07 Å and 2.55 Å, respectively. Additionally, the chloride (-Cl) group attached to the benzene ring interacts with Thr 87 to form a halogen bond of length 2.64 Å (**Figure 4C**). A52 interacts with the binding pocket to form four H-bonds. The furan nitro (-NO_2_) group forms an H-bond with Asn 114 of length 2.72 Å and a salt bridge of 3.56 Å with Glu 61. The amide-carbonyl forms H-bonds with Ser 25 and Ala 59 of lengths 3.5 Å and 2.05 Å, respectively. The amide-NH group interacts with Val 81 forming a H-bond of length 2.13 Å (**Figure 4D**).

### Microscale Thermophoresis revealed a high binding affinity of the shortlisted compounds with purified lumazine synthase

We next assessed the binding affinity of the three shortlisted compounds with the purified lumazine synthase by determining the dissociation constant (K_d_) with the help of MST. Details of cloning, purification, and characterization of these molecules are provided in the Supplementary Methods and **Supplementary Figures 2, 3, and 4**.

Our results presented in Figure 5 reveal that riboflavin, A10, A39, and A52 interact with lumazine synthase with the K_d_ values of 195 nM, 69.7 nM, 2.0 nM, and 58.6 nM, respectively (**Figure 5A-H)**, which suggests a direct effect of these compounds on the target protein RibH. Furthermore, A39 appears to have the highest binding affinity with RibH among the compounds tested.

**Figure 5.**
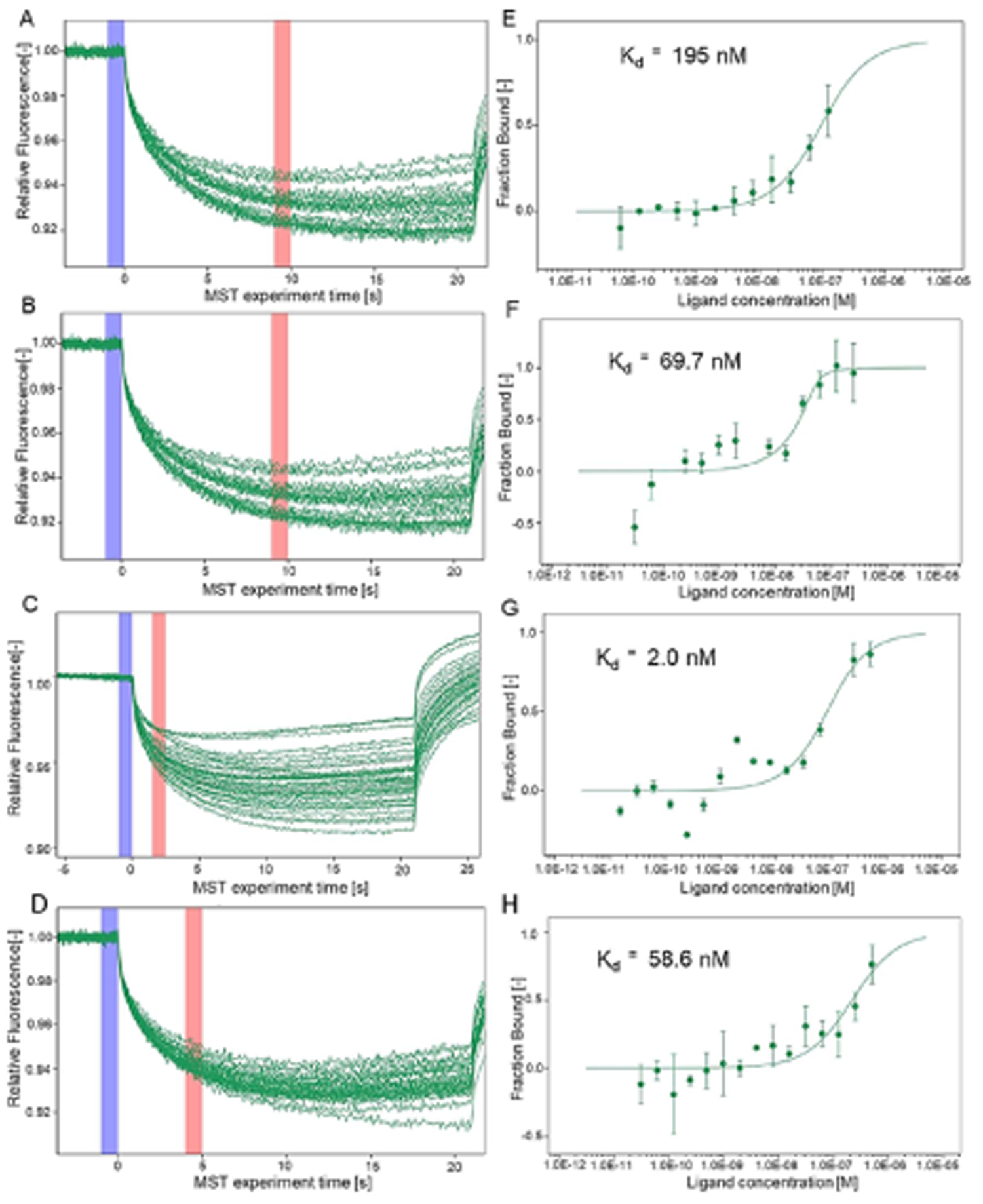
Microscale thermophoresis (MST) analysis of RibH protein with riboflavin and shortlisted drug molecules. Binding affinity of riboflavin and shortlisted drug molecules to RibH purified protein was determined by studying protein-ligand interaction using 16 concentrations (500 nM - 0.015 nM) of riboflavin or shortlisted drug molecules namely, A10, A39 and A52 with a fixed concentration of fluorescently labelled *M. tb* RibH protein (50 nM). MST traces corresponding to various concentrations are illustrated for, A. Riboflavin, B. A10, C. A39 and D. A52. Regression curves were generated to determine the bound fraction of the ligands, thereby calculating the binding affinity as K_d_ values. Binding affinity were determined to be E. 195 nM for riboflavin, F. 69.7 nM for A10, G. 2 nM for A39 and H. 58.6 nM for A52. The data for three independent experiments (n= 3) is presented as mean ± s.d.

### Antimycobacterial activity of the short-listed compounds

After the preliminary high-throughput virtual screening, fifty-five short-listed compounds were screened against *M. tb H37Rv* strain to evaluate their minimum inhibitory concentration (MIC) using microplate Alamar Blue assay (MABA) (**Figure 6A-C)**. Among 9 compounds showing MIC in the range of 0.78 - 12.5 µg/ml, A52 is found to be the most potent with a MIC of 0.78 µg/ml (**Figure 6C, Supplementary Table 1)**, whereas the MIC of both A10 and A39 is 1.56 µg/ml. Twelve compounds show moderate anti-tubercular activity in the range of 25 - 50 µg/ml and thirty-four show poor activity with MIC > 50 µg/ml, despite the high docking score **(Supplementary Table 1)**. We also assessed the MIC of these compounds in the presence of various concentrations of riboflavin (3.1 nM, 10.4 Nm, and 300 nM), however, we do not find any considerable change in the MICs of the RibH targeting molecules **(Supplementary Figure 5)**. These results suggest that *M. tb* is unable to procure riboflavin from exogenous sources to compensate for the loss of RibH activity, and hence can be considered a potential drug target. Based on these results, we further assessed A10, A39, and A52 as described below.

**Figure 6.**
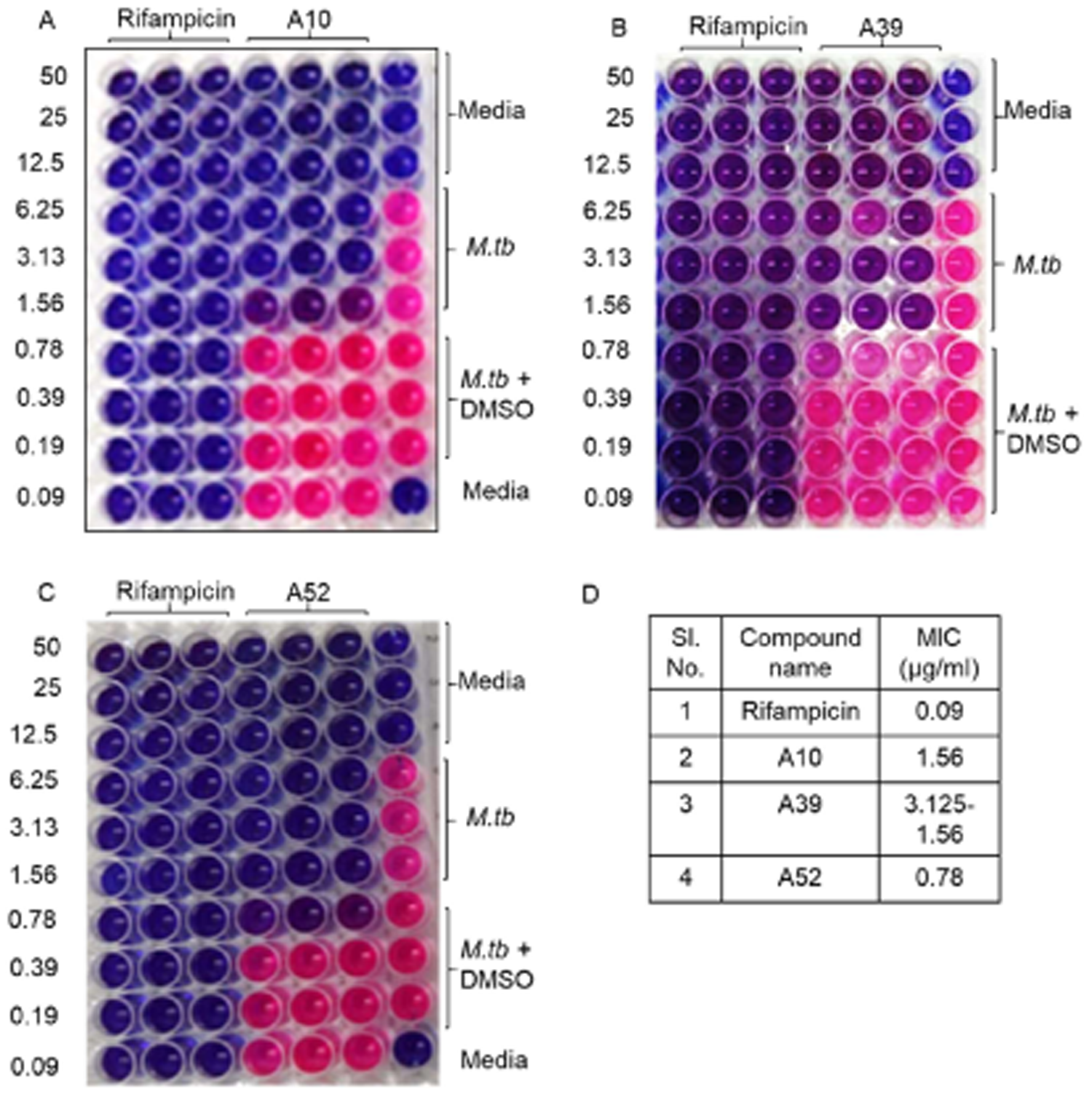
Anti-mycobacterial activity of shortlisted RibH inhibitors. The figure depicts the representative pictograms obtained on screening of anti-mycobacterial activity of three of the shortlisted compounds namely, A10, A39 and A52 against *M. tuberculosis H37Rv* by microplate Alamar blue assay (MABA). The assay was performed at least three times with three technical replicates in each case. Rifampicin and shortlisted compounds were tested at various concentrations between 50 µg/ml to 0.09 µg/ml (two-fold serial dilution). In our assay rifampicin exhibited an MIC of 0.09 - 0.048 µg/ml. A. Compound A10 exhibited an MIC of 1.56 µg/ml; B. Compound A39 exhibited an MIC of 3.125-1.56 µg/ml; C. Compound A52 exhibited an MIC of 0.78 µg/ml. D. Tabulation of MIC values obtained for various compounds with respect to rifampicin.

### Drugging lumazine synthase results in synergistic action with the first-line anti-TB drugs

We sought to determine if: (i) the antimycobacterial action of A10, A39, and A52 can be enhanced by combining with the first-line anti-TB drugs, and (ii) the addition of these compounds at the sub-MIC level increases *M. tb* susceptibility to the first-line anti-TB drugs. For this, a checkerboard MABA^34^ was employed, wherein A10, A39, and A52 were tested at various concentrations above and below MIC in combination with different concentrations at or below MIC of rifampicin (0.19-0.006 µg/ml) (**Figure 7A-C)** and isoniazid (0.39 - 0.012 µg/ml) **(Supplementary Figure 6 A-C)** against *M. tb H37Rv*. Strikingly, the addition of all three RibH-targeting drugs show synergistic action with rifampicin dramatically improving its MIC to ≤ 0.006 µg/ml, with 15-, 8-, and 7.5-fold reduction by A10, A39, and A52, respectively. The addition of rifampicin also enhances the antimycobacterial activity of A10, A39, and A52 resulting in a reduction in MIC by 4-, 4-, and 2-fold, respectively. Similarly, A10, A39, and A52 improve the efficacy of isoniazid with 15.8-, 32.5- and 15.8-fold reduction in MIC, respectively. **Table 1** depicts the fractional inhibitory concentration index (FICI), wherein combining these compounds with rifampicin and isoniazid results in synergistic action with FICI ≤ 0.5.

**Figure 7.**
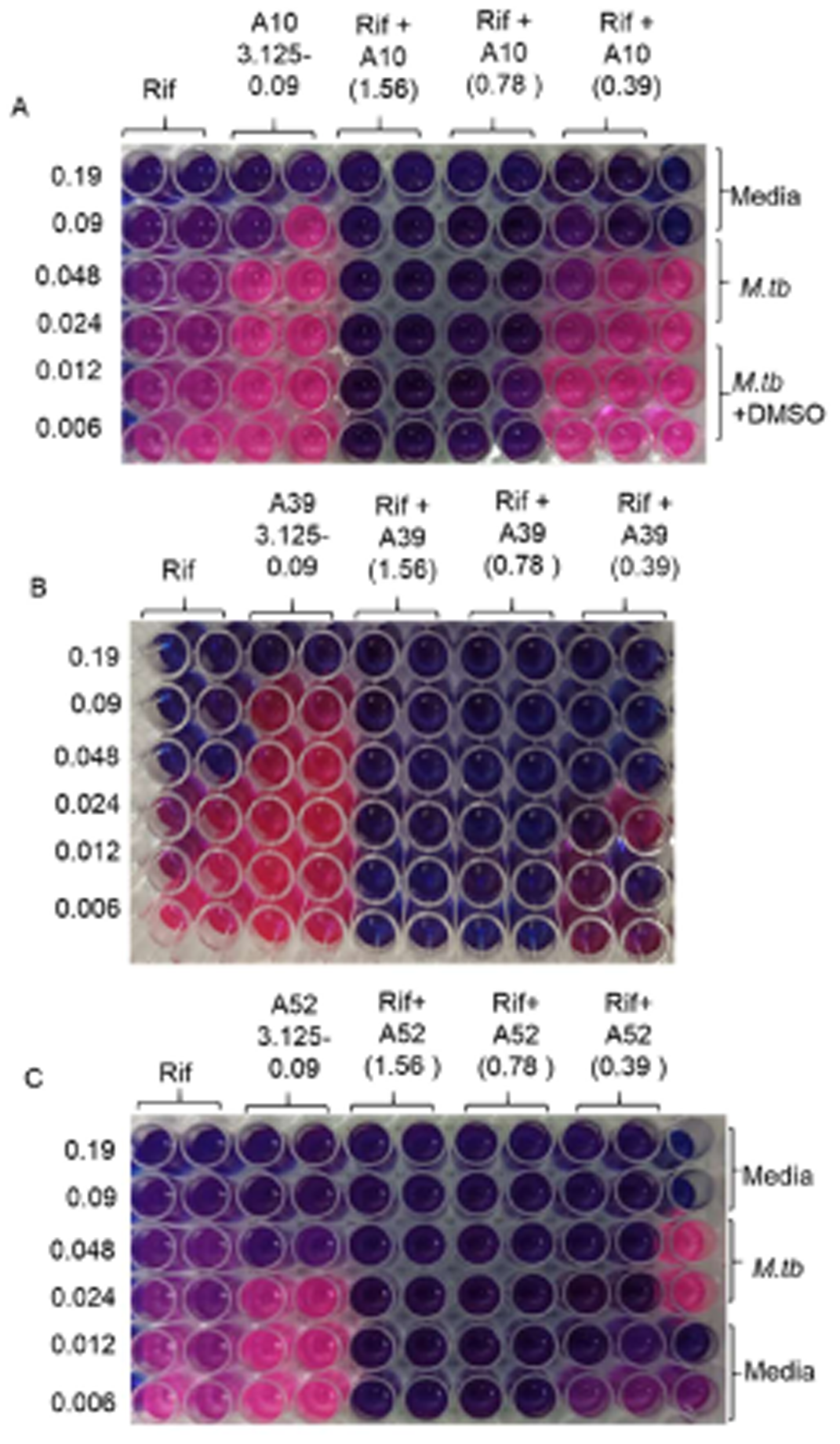
Anti-mycobacterial activity of short-listed compounds in combination with first-line anti-TB drug rifampicin. The figure depicts the representative pictograms obtained on screening of anti-mycobacterial activity of three of the shortlisted compounds against *M. tuberculosis H37Rv* in combination with rifampicin by microplate Alamar blue assay (MABA). The assay was performed at least three times with three technical replicates in each case. In each case (A, B and C) rifampicin was tested at various concentrations between 0.19 µg/ml to 0.006 µg/ml (two-fold serial dilution) either alone or in combination with various concentrations of A. A10, B. A39 and C. A52. These shortlisted compounds were either tested alone at various concentrations between 3.125-0.09 µg/ml (two-fold serial dilution) or at their respective MICs and sub-MIC concentrations in combination with various concentrations of rifampicin (MIC to sub-MIC range). A synergistic enhancement in antimycobacterial activity of rifampicin and shortlisted compounds was observed as described in Table 1. (For combined anti-TB activity with Isoniazid see supplementary Figure 5)

**Table 1.** Synergistic interactions between RibH targeted drugs and first line anti-TB drugs rifampicin and isoniazid. Table describes degree of interaction of pairwise combinations of rifampicin and isoniazid in combination with A10, A39 and A52 against *M. tb H37Rv.* Fractional inhibitory concentration index (FICI) was calculated using formula = FIC_A_ + FIC_B_ FIC_A_ = MIC of compound A in combination with B/MIC of compound A alone FIC_B_ = MIC of compound B in combination with A/MIC of compound B alone

| Compound | FICI for Rifampicin | Fold reduction in MIC of Rifampicin | FICI for Isoniazid | Fold reduction in MIC of Isoniazid |
| --- | --- | --- | --- | --- |
| A10 | 0.31 | 15 | 0.18 | 15.8 |
| A39 | 0.37 | 8 | 0.154 | 32.5 |
| A52 | 0.6 | 7.5 | 0.3 | 15.8 |
| Synergy is defined as FICI of $\leq 0.5$, antagonism as $\geq 4$. | | | | |

The observed synergism between RibH inhibitors causing functional ablation of the lumazine synthase is mirrored by genetic ablation of *ribH* using CRISPRi, such that *ribH*-KD strain (+ATc) shows enhanced susceptibility to killing by rifampicin in comparison to the *ribH*-KD strain (- ATc). Altogether, these observations establish an important link between lumazine synthase and mycobacterial susceptibility to various anti-TB drugs **(Supplementary Figure 7)**.

### Cytotoxicity and selectivity index of A10, A39, and A52

Based on the anti-mycobacterial activity, the three compounds were further selected for assessment of cytotoxicity and selectivity index using 3-(4,5-dimethylthiazol-2-yl)-2,5-diphenyltetrazolium bromide (MTT) assay, as described in methods. The cytotoxicity was assessed in human embryonic kidney 293 (HEK 293T) cells. Among the three potent compounds, A39 shows the least cytotoxicity at MIC, followed by A10 and A52. However, A52 displays the highest IC50 of 100 µg/ml corresponding to a selectivity index (SI) of 128 (**Table 2**). In contrast, A10 and A39 exhibit IC_50_ of 50 and 56 µg/ml, and a corresponding SI of ∼32 and ∼36, respectively. Most, importantly, at the MIC level all three compounds show minimal toxicity, (13.5%, 0%, and 19.82% by A10, A39, and A52, respectively) (**Table 2**).

**Table 2.**
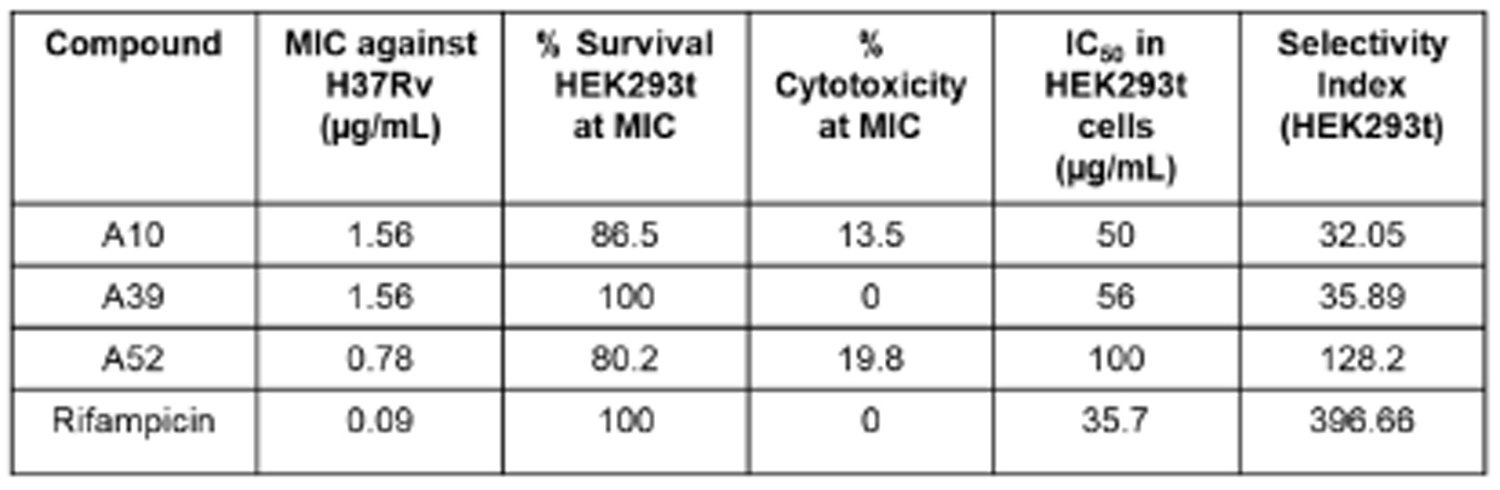
*In vitro* cytotoxicity and selectivity index of compounds A10, A39 and A52. Cytotoxicity of the RibH targeting drugs was measured by MTT assay against human embryonic kidney (HEK293t) cells and data (n=3) was analysed and expressed as IC_50_ (half-maximal inhibitory concentration). Selectivity index (SI) = IC_50_ (mammalian cells)/MIC (*M. tb H37Rv*).

### Efficient intracellular killing of *M. tb* by RibH inhibitors

To understand if these potent molecules can traverse the barrier of the host cell membrane to effectively kill the intracellular mycobacteria inside the macrophages, the intracellular antimycobacterial activity of A10, A39, and A52 is examined at various concentrations above and below the MICs (3.125, 1.56, 0.78 and 0.39 µg/ml), as described in methods. Infection of human macrophage line THP-1 by *M. tb H37Rv* and drug treatment is performed for 5 days, and on the 6^th^ day infected macrophages are lysed and colony forming units (CFU) are determined. Our results show that all the compounds are highly effective in the intracellular killing of *M. tb,* resulting in a significant reduction in colony forming units (CFU) in the case of A10, A39, and A52, respectively, compared to the untreated infected cells (*p* < 0.0001) (**Figure 8**, **Table 3**). Importantly, A52 is found to be the most potent among all the compounds that we tested across various concentrations causing a 99-94% reduction in CFU as compared to A39 (95-92% reduction in CFU) and A10 (95-81% reduction in CFU).

**Figure 8.**
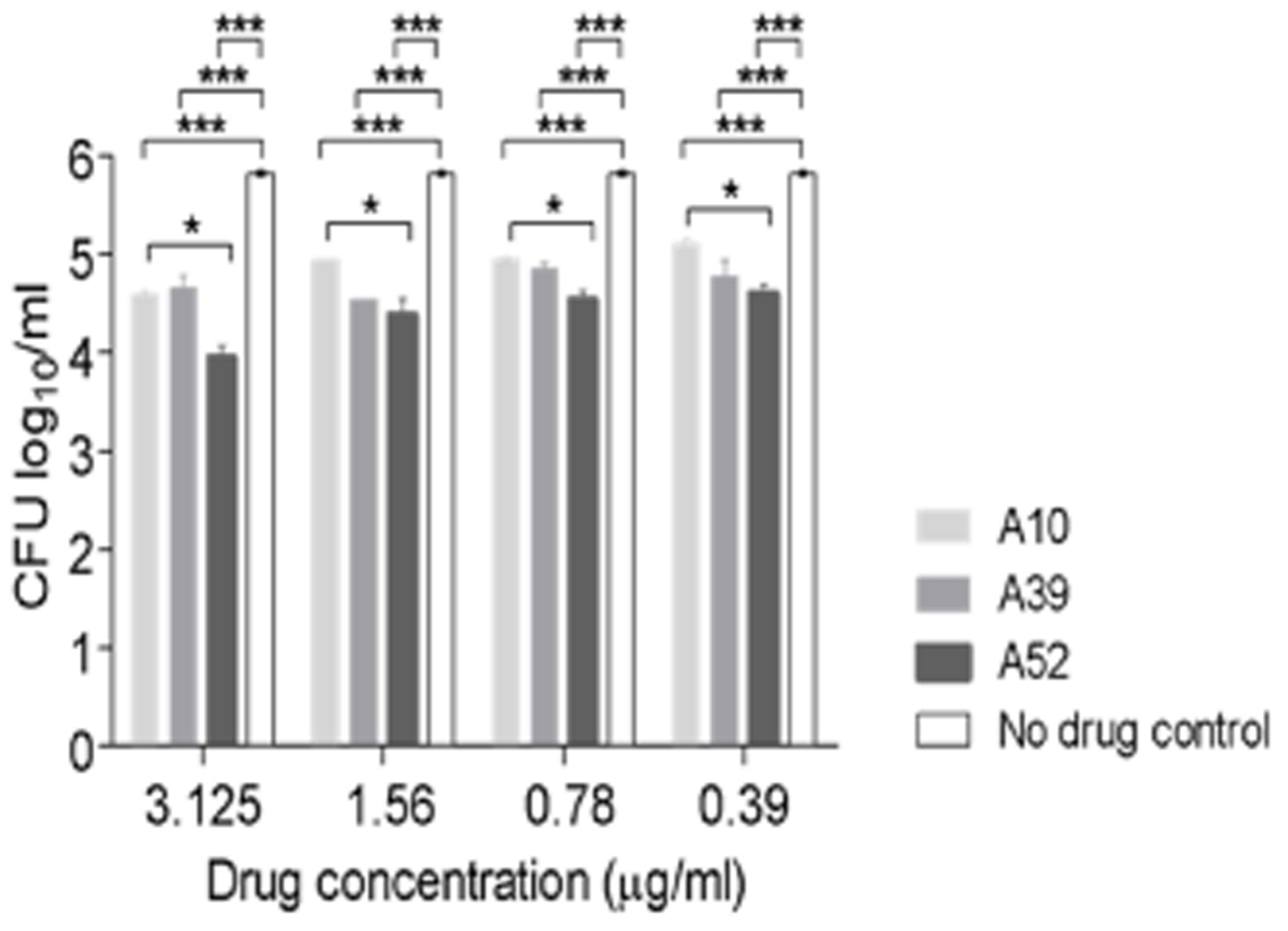
RibH targeting drugs reduce the intracellular growth of *M. tb.* in macrophages. Intracellular survival of *M. tb H37Rv* in THP-1 derived macrophages post five-days treatment with three different concentrations of A10, A39 and A52. THP-1 cells were infected with *M. tb. H37Rv* at an MOI of 1:5. Infected macrophages were treated with A10 or A39 at a concentration above MIC (3.125 µg/ml), MIC (1.56 µg/ml) and sub-MIC (0.78 µg/ml); or A52 at two concentrations above MIC (3.125 µg/ml-1.56 µg/ml) and MIC (0.78 µg/ml). CFU data (n=3) is represented as mean ± SEM log_10_CFU/ml. All the three compounds gave significant growth inhibition as compared to untreated control (***, *p* < 0.0001, *, *p* < 0.05; two-tailed Student’s t-test using GraphPad Prism Software). (For detailed CFU see table 3)

**Table 3:**
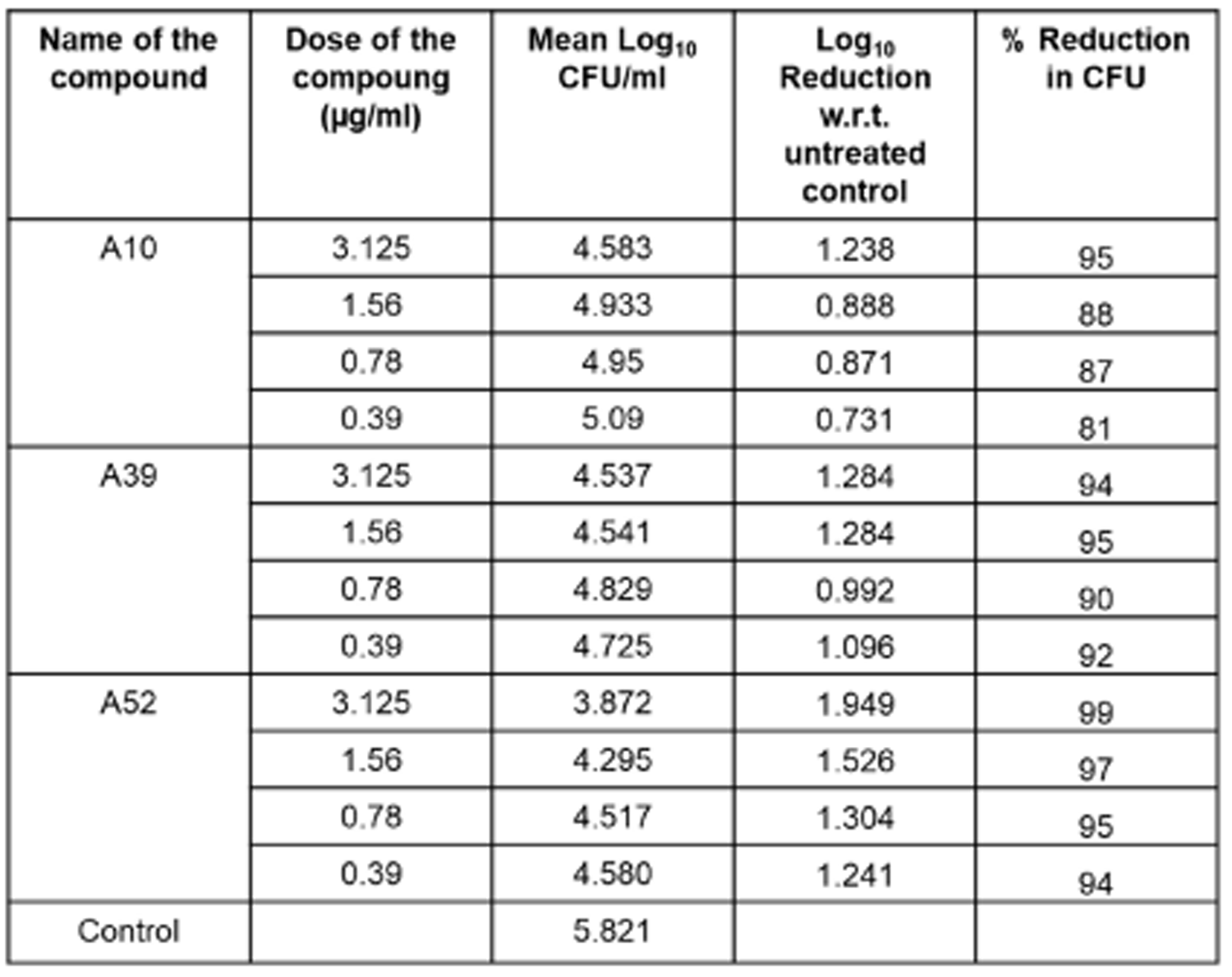
Intracellular antimycobacterial activity of short-listed compounds in THP-1 derived human macrophages. Table depicts CFU obtained on plating of infected THP-1 derived macrophages post five-days treatment with three different concentrations of A10, A39 and A52. THP-1 cells were infected with *M. tb*. *H37Rv* at an MOI of 1:5. Infected macrophages were treated with A10 or A39 at a concentration above MIC (3.125 µg/ml), MIC (1.56 µg/ml) and sub-MIC (0.78 µg/ml); or A52 at two concentrations above MIC (3.125 µg/ml-1.56 µg/ml) and MIC (0.78 µg/ml). CFU data (n=3) is represented as mean ± SEM log_10_CFU/ml. Reduction in CFU was calculated with respect (w.r.t) to untreated control, followed by calculation of % reduction in CFU as described in methods. Among the three compounds, A52 showed most potent activity in eradicating intracellular *M. tb*. *H37Rv*.

| Name of the compound | Dose of the compound (µg/ml) | Mean Log<sub>10</sub> CFU/ml | Log<sub>10</sub> Reduction w.r.t. untreated control | % Reduction in CFU |
| --- | --- | --- | --- | --- |
| A10 | 3.125 | 4.583 | 1.238 | 95 |
|  | 1.56 | 4.933 | 0.888 | 88 |
|  | 0.78 | 4.95 | 0.871 | 87 |
|  | 0.39 | 5.09 | 0.731 | 81 |
| A39 | 3.125 | 4.537 | 1.284 | 94 |
|  | 1.56 | 4.541 | 1.284 | 95 |
|  | 0.78 | 4.829 | 0.992 | 90 |
|  | 0.39 | 4.725 | 1.096 | 92 |
| A52 | 3.125 | 3.872 | 1.949 | 99 |
|  | 1.56 | 4.295 | 1.526 | 97 |
|  | 0.78 | 4.517 | 1.304 | 95 |
|  | 0.39 | 4.580 | 1.241 | 94 |
| Control |  | 5.821 |  |  |

### Lumazine synthase targeting drugs interfere with resuscitation in an *in vitro* dormancy model

We next tested, if RibH targeting drugs function against nutritionally starved *M. tb* in an *in vitro* dormancy/non-replicating persistence model **(Supplementary Figure 8)**, and whether these drugs interfere with the ability of the bacteria to resuscitate when exposed to the nutrient-rich conditions. We mimicked dormancy by starving *M. tb H37Rv* in phosphate-buffered saline for a period of six week, and then exposed the bacteria to A10, A39, and A52 with rifampicin and isoniazid as control drugs. We followed *M. tb* growth by most probable number (MPN) assay, as described in the methods. Remarkably, our findings reveal that these drugs not only act upon the nutritionally starved bacteria but also interfere with their ability to resuscitate despite the presence of nutrient-rich conditions, which otherwise favors bacterial growth in the absence of drugs. All the compounds show potent activity against nutritionally starved *M. tb H37Rv* (**Figure 9**), however, A10 causes maximum growth inhibition resulting in ∼2.9 log_10_ reduction in CFU compared to that of untreated control (*p* < 0.0001). Noteworthy to mention, A10 performs much better than the first-line anti-TB drugs, resulting in 0.6 log_10_ and 1.081 log_10_ fewer bacillary counts compared to rifampicin and isoniazid (*p* < 0.01), respectively (**Figure 9**). A10 also shows better activity compared to A39 (*p* < 0.01) and A52 (*p* < 0.05). Treatment with A52 and A39 results in 2.02 log_10_ and ∼1.8 log_10_ reduction, respectively, compared to the untreated control (*p* < 0.0001). However, the potency of A52 and A39 is comparable to that of rifampicin (2.29 log_10_ reduction) and isoniazid (1.8 log_10_ reduction).

**Figure 9.**
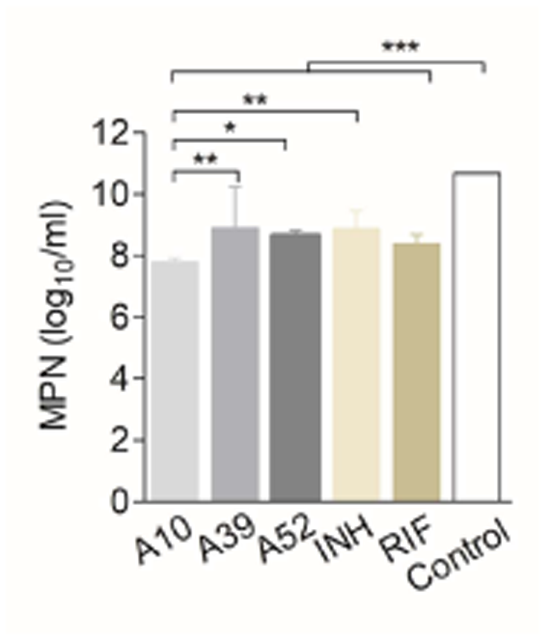
RibH targeting drugs interfere with growth (resuscitation) of nutritionally starved *M. tuberculosis* in an *in vitro* dormancy model. *M. tb*. *H37Rv* cultures were starved for 6 weeks in phosphate buffered saline, followed by treatment with the compounds namely, A10, A39 and A52 or first-line anti-TB drugs isoniazid or rifampicin at an equimolar concentration of 10 µM. Bacterial count estimation was carried out using the MPN (most probable number) assay. CFU data (n=3) is represented as geometric mean ± SEM log_10_CFU/ml. Data represents geometric mean with 95% CI for (n=3). All the three compounds gave significant growth inhibition as compared to untreated control (*p* < 0.0001, two-tailed t-test using GraphPad Prism Software). Compound A10 shows significant reduction in *M. tb* growth as compared to the first line anti-TB drug Isoniazid (p < 0.05, One-way ANOVA using GraphPad Prism Software).

### RibH-a highly conserved drug target across various mycobacterial species

Having established the essentiality, druggability, synergism with existing drugs and safety profile of drugs targeting RibH of *M. tb* we next analyzed if RibH is conserved during evolution among various mycobacterial species belonging to Mycobacterium tuberculosis complex (MTC) and Non-tuberculous mycobacteria (NTM).

We first conducted multiple sequence alignment of RibH protein for various mycobacterial species and determined the phylogenetic relationship between them **(Supplementary Figures 9 and 10**). RibH protein was found to be highly conserved in nature across the MTC, NTM, and *M. leprae* with minimum divergence. We then specifically compared the crucial amino acids present in the drug binding and co-crystal ligand binding pocket across various mycobacterial species. Remarkably, the binding site for A10, A39, A52, and TP6 was found to be highly conserved (95-100 % identity) across the mycobacterial species **(Supplementary Tables 2 and 3**). Especially, major mycobacterial pathogens were found to have 100% identity in the drug-binding pocket, thereby increasing the likelihood of broad-spectrum activity of these drugs for treating various other mycobacterial infections. Interestingly, drug binding pocket was also found to be conserved in *M. leprae* (causative agent of leprosy in humans and belonging to *Mycobacterium leprae complex*, MLC). The conserved nature of the RibH protein and drug binding pocket increases the translational potential of the drugs developed by us, making it suitable for future testing against MTB, NTMs, and MLC with broad spectrum applications for treating human and animal diseases caused by mycobacteria.

## Discussion

The major challenge of the current TB treatment is prolonged therapy (6-9 months), which often leads to non-compliance and the development of drug resistance to the existing drugs. The emergence of MDR and XDR TB demands the development of novel antibiotics. Existing drugs primarily target well-established pathways, making *M. tb* highly susceptible to evolving and developing drug resistance mutations^3^. To overcome this challenge, targeting new pathways that are previously unexplored for drug development is expected to reduce the likelihood of acquiring or having pre-existing drug resistance^5^. Most importantly, if the essentiality could be proven across diverse physiological states experienced by *M. tb* inside the host such as macrophages, dormancy etc, targeting such essential pathway(s) further increase the chances of developing a successful drug with a widespread impact on *M. tb* physiology.

Target-based or gene-to-drug model for drug development is considered to have the highest chance of success provided the target gene is essential for bacterial survival, unique to bacteria (absent from humans), and functionally irreplaceable. Strategic drug designing may involve targeting multiple essential pathways by a single drug^35^ or drugs are designed against such pathways which produce metabolites, for example, cofactors required for a plethora of indispensable processes in mycobacteria^5, 35^. A target becomes more promising if it is conserved in evolution and shows a minimal divergence in the active site or protein sequence allowing similar antibiotic liability and druggability across various mycobacterial species/strains^36^. It also signifies that targeting such a protein or pathway would have substantial one health significance due to the broad-spectrum activity against a variety of mycobacterial species responsible for TB, leprosy, bovine TB, or opportunistic NTM infections which have been on the rise globally recently. Importantly, recent studies suggest a link between cystic fibrosis and NTM infections in patients especially those with immunosuppressive conditions ^37, 38^. The conserved nature of the drug binding site is also suggestive of the fact that these drug targets are less prone to mutations or genetic variations and hence are less likely to develop drug resistance.

With a premise to cause a widespread effect on mycobacterial physiology, crippling several processes at once, we identified drugs targeting the riboflavin biosynthesis pathway, a crucial source of important cofactors-FAD and FMN in *M. tb*. Among the various enzymes involved in the pathway, we identified RibH to be the most suitable candidate for several reasons. First, it was found to be highly conserved across the evolutionary scale of mycobacteria **(Supplementary Figures 9 and 10 and Supplementary Table 3)**. With only a single copy of this gene in the *M. tb* genome and no evidence for redundancy in its enzymatic function, interference at the penultimate step of riboflavin synthesis catalyzed by lumazine synthase is anticipated to substantially curtail the pathway. The *ribH* gene has also been listed by various studies as an essential gene required for mycobacterial survival^12, 13^. As riboflavin biosynthesis is unique to microorganisms and is absent from humans, targeting this pathway is all the more promising. Moreover, to the best of our knowledge, there are no reported riboflavin transporters in *M. tb*, hence we hypothesized that functional interference of this enzyme employing specific drug molecules would possibly leave *M. tb* metabolically unfit failing to replicate. We provide evidence for the same in this study, wherein exogenous supplementation with riboflavin or a cocktail of FMN/FAD failed to compensate for the loss of enzyme function in knockdown studies and hence the MICs of drug molecules remain unaltered.

Our high-throughput molecular docking approach allowed us to screen ∼0.6 million compounds capable of binding to RibH. Through this screening process, we discovered three highly potent compounds that exhibited promising antimycobacterial activity against the *M. tb* H37Rv as standalone drug. The most noteworthy finding of our research was the significant synergism of these compounds with the first-line anti-TB drugs, rifampicin, and isoniazid. These observations are reminiscent of several earlier studies wherein, mutations in genes or drugs targeting cell wall biosynthesis or FtsZ resulted in defective growth along with altered cellular morphology, bacillary length, and cell wall permeability to first-line TB drugs^39–42^. Drugs like ethambutol was also found to alter membrane fluidity which renders *M. tb* susceptible to drug classes it is normally resistant against^43^. Future studies to understand the changes in mycobacterial cell wall will provide mechanistic insights into the increased vulnerability of *M. tb* to first-line anti-TB drugs on exposure to RibH targeting drugs.

The first-line anti-TB drugs are known for their high toxicity and side effects in 3-40% of the patients, resulting in non-compliance to TB therapy^44, 45^. *In silico* analysis of lead compounds targeting RibH predicts these compounds to possess good ADME properties **(Supplementary Table 8)**, bioavailability, and gastrointestinal absorption and are also predicted to have better cellular retention as they do not induce P-glycoprotein efflux pumps, unlike rifampicin^46–48^. Hence, RibH-targeting drugs are predicted to have good pharmacokinetic properties with minimum drug-drug interactions and toxicity **(Supplementary Table 8, Table 2**). Possibly a better cellular retention due to non-induction of P-glycoprotein^49^ would have contributed to an effective intracellular mycobacterial clearance as observed in this study (**Figure 8 and Table 3**).

Most of the anti-TB drugs such as rifampicin and isoniazid, used to treat TB caused by *M. tb*, are primarily effective against actively replicating bacteria. However, their activity declines against non-replicating persister cells when tested in an *in vitro* dormancy model^50^. Among the various reasons for the reduced activity of rifampicin against non-replicating bacteria, poor penetration of the drug into dormant bacterial cells, and the downregulation of target RNA polymerase during dormancy are the top reasons^51–54^. We observed that exposure to RibH targeting drugs to nutrient-starved non-replicating *M. tb* interferes with their resuscitation highlighting an enhanced requirement for Flavin-based cofactors during reactivation. As we observed a synergism with rifampicin and isoniazid in actively replicating *M. tb*, we anticipate that due to the multi-faceted role of riboflavin and cofactors derived thereof, newly discovered RibH targeting drug molecules may improve the sensitivity of dormant bacteria towards first-line anti-TB drugs, which needs to be tested in our future studies. Increased expression of RibH in oxidative stress and non-replicating persistent *M. tb*^55^ further signifies the importance of targeting this pathway during dormancy.

Application of transcriptomics, proteomics, and metabolomics in our future studies would underpin the widespread impact of genetic or functional ablation of RibH on *M. tb* and may also shed light on the increased sensitivity of *M. tb* to rifampicin and isoniazid on exposure to newly designed drugs in this study. Our future research would involve identifying the *M. tb* resistant mutants towards the newly discovered drugs and studying the impact of those mutations on protein structure and function employing protein crystallography and other biochemical enzymatic assays. Another limitation of our study is our inability to study enzyme activity in the presence of various inhibitors due to commercial non-availability of the substrates of lumazine synthase, however, our MST data signifies direct binding of the inhibitors to the RibH enzyme with high affinity. Although we do show that one of the compounds is far more effective in preventing the growth/resuscitation of dormant bacteria in the *in vitro* dormancy model, it is worth noting the *in vitro* models may not be able to fully recapitulate the intricacies of dormancy inside the human host. As the response of non-replicating persister cells can vary in different conditions encountered inside the host such as hypoxic granulomas or host immunity, our future research focus is to understand the *in vivo* activity of these newly discovered drugs in small animal models to recapitulate active disease and dormancy stages. Good ADME properties as shown in this study, increase the chance of efficacy in *in vivo* models.

Having established, *M. tb* RibH being a potential drug target, our study is anticipated to promote future research toward designing compounds with better fit, activity, and ADME properties. Given the limitations of rifampicin and isoniazid against non-replicating persister cells, and improved MIC of rifampicin by combination therapy observed in our checkerboard MABAs, combining multiple drugs with different mechanisms of action, could achieve effective clearance of both actively replicating bacteria as well as non-replicating persister cells, thereby improving the treatment outcomes. Future studies comparing and combining these newly discovered drugs with other anti-TB drugs, such as pyrazinamide, fluoroquinolones^56^ (e.g., moxifloxacin), or bedaquiline^57^ which have demonstrated better activity against NRP would reveal their synergy and interactions with drugs of the anti-TB regimen, other than rifampicin and isoniazid.

In summary, our study not only validates the crucial role of RibH in the survival and growth of *M. tb* but also identifies three potent compounds with promising antimycobacterial activity. These findings open up new avenues for the development of improved treatments against TB. As RibH protein and the drug binding sites are found to be highly conserved, the leads identified in this study have the potential to tackle a variety of geographically widespread *M. tb* lineages causing active, persistent, and reactivation TB^3^. The combination of these compounds with existing first-line anti-TB drugs also presents a potential strategy for overcoming drug resistance and enhancing treatment outcomes. Further preclinical studies are warranted to evaluate the full potential of these compounds and study their impact on TB control.

## MATERIAL AND METHODS

### Ethics Statement

All the *in vitro* studies involving risk groups 3 and 2 *M. tuberculosis* are done as per the guidelines and approval of the institutional biosafety committee.

### Bacterial strains and growth conditions

*M. tb* strain H37Rv was obtained from ATCC (ATCC27294 strain). Frozen stocks were revived and cultured in Middlebrooks 7H9 broth (Himedia) supplemented with 10% oleic acid-albumin-dextrose-catalase (OADC) (Himedia), 0.4% glycerol (Himedia) and 0.05% Tween-80 (Sigma) in a BSL3 facility. For estimation of colony forming units, Middlebrooks 7H11 agar (Himedia) supplemented with 10% oleic acid-albumin-dextrose-catalase (OADC) (Himedia) and 0.5% glycerol (Himedia) was used.

The knockdown strain of *ribH* gene of *M.tb* used in this study was derived from mc^2^ 7902 (*H37Rv* Δ*panCD* Δ*leuCD* Δ*argB*)^28^ and was cultured in Middlebrook 7H9 broth or 7H11 agar supplemented with 10% oleic acid-albumin-dextrose-catalase (OADC) (Himedia) along with 0.5% glycerol, 0.05% tyloxapol and pantothenate, leucine and arginine (PLA) supplement containing l-pantothenate (24 mg/liter); l-leucine (50 mg/liter); and l-arginine (200 mg/liter) to circumvent synthetic lethality^28^. The knockdown strains were selected on kanamycin (Kan) and hygromycin (Hyg) at concentrations of 25 μg/ml and 50 μg/ml, respectively as described^30^.

DH5-α and BL21(DE3) strains of *E. coli* (NEB) were used for cloning and protein expression studies. *E. coli* strains were cultured in Luria Bertani (LB) broth (Himedia) or LB agar containing Kanamycin (Himedia) **(Supplementary Table 4) Construction of Knockdown Strains** To achieve the repression of *ribH*, a pair of complementary oligonucleotides (Cr_UP: GATCTTTCCGTGCCAGCTG and Cr_DN: CGCAGCTGGCACGGAAAGATCCATG) specific to *Rv1416* ORF between 73-93bp downstream to the 5’ end, were synthesized, annealed and cloned in a hygromycin-resistant (listed in **Supplementary Table 5)**, *E. coli-mycobacteria* shuttle plasmid pGrna2 at *Sph*I-*Acl*I sites, as previously described^29, 30^, and illustrated in **Supplementary Figure 1**. The recombinant pGrna2 plasmid containing *Rv1416*-specific guide sequence was transformed into a kanamycin-resistant *M. tb* mc^2^ 7902::pTetint-dcas9^29^ to generate knockdown strain, namely, *ribH*-KD. Suppression was achieved by treatment of bacterial cultures with various concentrations of Anhydrotetracycline (ATc) in the range of 50-100 ng/ml for 4-9 days (unless indicated otherwise).

### *In vitro* growth analysis

Growth of *ribH*-KD strain of *M. tb* was monitored in 7H9 broth in the absence or the presence of 50 ng/ml and 100 ng/ml Anhydrotetracycline (ATc) at 37^◦^C, 200 rpm. Further, to investigate if the *ribH* knockdown strain takes up FAD/FMN or riboflavin when added exogenously, growth of the knockdown strain was assessed in nutrient-rich and chemically defined synthetic media containing FAD/FMN (2.5μM each) and riboflavin (10 μM)^58^. The growth was monitored by measuring optical density spectrophotometrically at a wavelength of 600nm (OD_600_) for 9 days. The optical density was used to create time-stamped growth curves.

### RNA extraction and Quantitative RT-PCR

The total RNA was extracted from the *M. tb* using Trizol reagent (Sigma) and purified using RNeasy Plus mini kit (Qiagen) as per the instructions given by the manufacturer. cDNA synthesis was done using the iScript cDNA synthesis kit (Bio-Rad). Then the qRT-PCR was performed with iTAq Universal SYBR Green Supermix (Bio-Rad), employing *ribH* gene-specific primers, and 15 ng cDNA. Gene expression was quantified using the qRT-PCR system (Roche LightCycler 480). The expression of *ribH* in the knockdown strain was compared to the parent strain (*mc^2^ 7902*) and *sigA* was used as an internal control for normalization. The primers used are listed in **Supplementary Table 5**.

### Scanning Electron Microscopy (SEM)

Scanning electron microscopy was performed to examine the alterations in the morphology of bacteria. Briefly, 5-ml bacterial cultures were pelleted down and resuspended in 5% glutaraldehyde (Himedia) prepared in 0.1M phosphate buffer, pH 7.2 and incubated for 30 minutes, followed by centrifugation at 4500rpm. The resulting pellets were washed in 0.1M phosphate buffer for 10 minutes followed by treatment with 1% osmium tetroxide (Himedia) (in 0.1M phosphate buffer, pH 7.2) for 1 hour at room temperature. The pellet was washed with distilled water before dehydrating the sample with sequential ethanol series (35%, 50%, 70%, 95%, and absolute ethanol) each for 10 minutes. The samples were air-dried and mounted onto the stub with carbon tape. Mounted samples were sputtered with gold-palladium alloy (Leica UltraMicrotome EM UC7, Sputter Coater) and visualized in Field emission-scanning electron microscope (ApreoVac, FEI). The bacillary length was measured using ImageJ Software.

### Computational Details

*In silico* analyses were performed in Dell Precision T7610 workstation (8 processors; 8 GB RAM; ZOTAC 3GB graphics; Maestro 9.8, Schrodinger, New York, U.S.A) workstation running on Redhat 6.1 Linux environment. Molecular docking calculations were conducted using two different strategies i.e., pharmacophore-based virtual screening (PBVS) and Structure-based flexible docking using Glide application in the Maestro 9.8 software package (Schrodinger, LLC, New York, 2015). The BITS in-house and Asinex compound libraries were first docked against *M. tb* RibH using PBVS approach by e-pharmacophore generation and validation leading to the identification of 15 molecules from Asinex library and 14 from BITS in-house library. We also performed blind docking on the BITS in-house library and short-listed an additional 26 molecules.

### Protein Preparation

Crystal structure of *M. tuberculosis* lumazine synthase (encoded by *ribH*, *Rv1416*) with Protein Databank (PDB) ID 2C92 bound to *3-(1,3,7-trihydro-9-d-ribityl-2,6,8-purinetrione-7-yl) pentane 1 phosphate* (TP6) having a resolution of 1.60 Å was considered for this study. There are other crystal structures published for this protein (with PDB IDs - 1W19, 1W29, 2C94, 2C97, 2C9B, 2C9D, and 2VI5), however we have used 2C92 due to the best crystal structure resolution among all the published structures^21^. The protein exists in homopentameric form with substrate binding sites at the interface of two protein chains/subunits, hence for protein preparation and all the calculations, the complete dimer was used. Protein was prepared using the protein preparation wizard of Maestro 9.8, Schrodinger, New York, U.S.A. The co-crystal structure was pre-processed and water molecules within 5 Å distance from the ligand were removed, missing hydrogens and loops were added using Schrödinger protein preparation wizard. In addition, protein preparation also involved the addition of the bond orders and formal charges along with hydrogens to the hetero groups of the protein. Followed by an energy minimization to a convergence of RMSD 0.30 Å using OPLS_2003 as a force field.

### Ligand Database Screening

#### Compound Libraries

For identifying lead compounds that could potentially inhibit the enzymatic activity of lumazine synthase and hence serve as an anti-mycobacterial agent, we utilized two libraries, the first library consists of ∼ 3000 compounds from our BITS (Birla Institute of Technology and Science) *in-house* database containing compounds that have ADME properties commonly observed in drug and drug-like compounds^49^ and ASINEX compound library comprising of ∼0.57 million compounds (https://www.asinex.com/screening-libraries-(all-libraries)).

### Hypothesis generation for energy-optimized structure-based E-pharmacophore

Glide energy grids were generated for the prepared protein-ligand complex. The binding site of RibH was defined by a rectangular box surrounding the ligand in the X-ray structure. The ligand was refined using Glide and default settings were used for the refinement and scoring. Starting with the refined ligand, pharmacophore sites were automatically generated with Phase with the default set of six chemical features: hydrogen bond acceptor (A), hydrogen bond donor (D), hydrophobic (H), negative ionizable (N), positive ionizable (P), and aromatic ring (R). Hydrogen bond acceptor sites were represented as vectors along the hydrogen bond axis in accordance with the hybridization of the acceptor atom. Hydrogen bond donors were represented as projected points, located at the corresponding hydrogen bond acceptor positions in the binding site. Projected points allow the possibility for structurally dissimilar active compounds to form hydrogen bonds in the same location, regardless of their point of origin and directionality. Each pharmacophore feature site is first assigned an energetic value equal to the sum of the Glide XP contributions of the atoms comprising the site. This allows sites to be quantified and ranked based on these energetic terms.

### Database Preparation

The commercial chemical ASINEX virtual library with over 600,000 compounds & in-house 3000 compounds library were processed to avoid redundancy and Lipinski filters were applied to select compounds that have drug-like properties. The purpose of the redundancy check is to avoid structures with the same SMILES notation (Singleline entry system for structures) and also to remove conformers of very similar conformation using an RMSD cutoff of 1.0 Å. Database molecules were prepared using LigPrep and Epik to expand protonation and tautomeric states at pH 7.0 (± 2.0 pH units). For each ligand, LigPrep generates a maximum of eight tautomeric forms. If for a compound, chirality is already specified, it is retained. However, if the compound chirality is undefined, at most 32 stereoisomers were generated for each ligand used for screening. Conformational sampling was performed for all database molecules using the ConfGen search algorithm. We employed ConfGen with the OPLS_2005 force field and a duplicate pose elimination criterion of 1.0 Å RMSD to remove redundant conformers. A distance-dependent dielectric solvation treatment was used to screen electrostatic interactions. A maximum relative energy difference of 10.0 kcal/mol was chosen to exclude high-energy structures. Using Phase, the database was indexed with the automatic creation of pharmacophore sites for each conformer to allow rapid database alignments and screening.

### E-Pharmacophore Database Screening

To dock the compound libraries into the generated grid, the libraries were first pre-processed to create an e-pharmacophore. e-pharmacophore is defined as a hypothetical ideal 3D orientation or spatial and electronic characteristics of the functional groups of a molecule necessary to ensure optimal interactions with a specific target protein to achieve a biological response. For the E-pharmacophore approach, explicit matching was required for the most energetically favorable site, with a scoring cutoff better than −1.0 kcal/mol. Screening molecules were required to match a minimum of 4 sites. Distance matching tolerance was set to 2.0 Å as a balance between stringent and loose-fitting matching alignment. Database hits were ranked in the order of their Fitness score as implemented in the default database screening in Phase. We validated the pharmacophore by docking the co-crystal ligand TP6. E-pharmacophore generation involves two steps, in the first step based on the co-crystal structure of the protein with a ligand (TP6), e-pharmacophore features such as hydrogen bond acceptor (A), hydrogen bond donor (D), hydrophobic (H), negative ionizable (N), positive ionizable (P) and aromatic ring (R) were assigned to the bound ligand using the Glide tool.

### Molecular Docking

Database ligands were docked into the binding sites of RibH with Glide utilizing the high-throughput virtual screening (HTVS) scoring function to estimate protein-ligand binding affinities. The center of the Glide grid was defined by the position of the co-crystallized ligand. Default settings were used for both grid generation and docking. Compounds with best docking and GLIDE scores were then subjected to GLIDE XP screening. Docked protein and ligand datasets were visualized and molecular surface complex pictures were generated using Maestro. Using this tool, we identified 17 compounds from ASINEX

### Structure-based design by virtual screening

#### Grid generation

The receptor site or the active site at the interface of two subunits, where the inhibitors are to be docked was generated using the ‘generate grid’ sub-application of the Glide tool in Maestro 9.8. For the generation of the receptor grid, active site residues were selected to locate the coordinates of the receptor center. The grid has default parameters of van der Waals scaling factor of 1.0 and charge cut off of 0.25 subject to an OPLS_2003 force field. The generated grid was then utilized as an active site for docking [in-house database of 3000 compounds] using the ‘extra precision’ (XP) flexible docking method located in the Glide tool. Using this tool, we further identified 26 compounds.

### Determination of minimal inhibitory concentration (MIC) of shortlisted compounds against *M. tuberculosis* by microplate Alamar Blue assay (MABA)

MIC for the new compounds identified through *in silico* analysis was carried out using methods as described^59^. Briefly, *M. tb* strain H37Rv (ATCC27294 strain) was cultured in Middlebrooks 7H9 broth (Himedia) supplemented with 10% OADC (Himedia) and 0.05% Tween-80 (Sigma) for 4-5 days. The culture was grown to the mid-log phase until an OD_600nm_ of 0.4-0.8. The culture was then diluted to obtain an OD_600nm_ of 0.01 corresponding to ∼10^6^ CFU/ml. Based on the docking score and chemical structure 40 compounds were selected from BITS *in-house* library and 15 compounds from the ASINEX compound library (**Supplementary Table 1**). These compounds along with standard anti-TB drugs - Rifampicin (Himedia) and Isoniazid (Sigma) were dissolved in absolute DMSO at a concentration of 2mg/ml to prepare the drug stock. The drug stock was then serially diluted in a 96-well plate to achieve a final concentration in the range of 50 µg/ml to 0.006 µg/ml. To each well containing the desired concentration of compounds or standard anti-TB drugs, 0.1 ml of the bacterial culture (∼10^5^ CFU/well) was added in a final volume of 200 µl. The plate was then sealed and incubated at 37°C for 6 days. For the Alamar Blue assay, a mixture of resazurin sodium salt (Sigma) and Tween-80 was freshly prepared in autoclaved single distilled water to obtain a final concentration of 1.27 mM and 8%, respectively and the solution was then filter sterilized using 0.2 µ syringe filter. On day 6, 30 µl of this freshly prepared sterile solution is added to each well of the plate and incubated for 16-18 hours at 37°C in the dark. Color change from purple to pink is noted and photographed, wherein, purple color indicates a lack of growth whereas, pink indicates live bacteria. MIC is determined from the minimal concentration of the antibiotics or compounds exhibiting purple color.

The three most potent compounds identified from the above MABA screen were then tested in combination with the first-line anti-TB drugs. For this experiment, MABA was performed with minor modifications. Briefly, for each of the three potent compounds, 3-4 concentrations were typically tested, (a) at MIC (b) at two concentrations below, and (c) at one concentration above MIC in the range of 3.125 µg/ml to 0.19 µg/ml (two-fold serial dilution). The standard anti-TB drugs were tested in the range of 0.19 µg/ml to 0.006 µg/ml (two-fold serial dilution) for rifampicin and 0.39 µg/ml to 0.012 µg/ml (two-fold serial dilution) for isoniazid. To achieve this, 100 µl of first-line anti-TB drug at 2X of desired concentrations were added to the wells, followed by the addition of 50 µl of 4X concentration of the short-listed compounds.

Following the addition of compounds, bacterial culture was diluted to obtain an OD_600nm_ of 0.02, and 50 µl was added to each well to obtain a final volume of 200 µl/well containing ∼10^5^ CFU/well. The plate was then sealed and incubated at 37°C for 6 days, followed by Alamar Blue addition and analysis of results as described above. The fractional inhibitory concentration index (FICI) was calculated to determine the synergistic effect of these compounds with rifampicin and isoniazid as described^60^. The fractional inhibitory concentration for each compound was calculated as: Fractional inhibitory concentration of compound A: FIC_A_ = (MIC of compound A in the presence of compound B)/ (MIC of compound A alone); Fractional inhibitory concentration of compound B: FIC_B_ = (MIC of compound B in the presence of compound A)/ (MIC of compound B alone). FICI was calculated as FIC_A_ + FIC_B._ Synergism is defined as FICI ≤ 0.5, antagonism as FICI ≥ 4 and no interaction if the FICI values range from 0.5-4.

### Determination of Cytotoxicity of shortlisted compounds by MTT assay

To determine the cytotoxicity of the shortlisted compounds, 3-(4,5-Dimethylthiazol-2-yl)-2,5-Diphenyltetrazolium Bromide (MTT) cytotoxicity assay was performed as per manufacturer’s instructions. This assay allows the measurement of cellular metabolic activity as an indicator of cell viability, proliferation, and cytotoxicity. Briefly, HEK293T cell line (human embryonic kidney cell line) was maintained in high glucose Dulbecco’s Modified Eagle’s Medium (DMEM) with 4.5 gm/liter glucose, 2 mM L-glutamine, 25 mM HEPES buffer, and 3.7 gm/liter sodium bicarbonate (Himedia) supplemented with 10% fetal bovine serum and 1X Pen Strep (Penicillin and streptomycin solution) at 37^°^ C and 5% CO_2._ In a 96-well plate, 2 ×10^4^ cells were seeded and allowed to get adhered to the dish for 12 hours. Following, adherence, all the media was replaced with 100 µl of fresh media (without antibiotic). In a separate 96-well plate, the three shortlisted compounds were 2-fold serially diluted in complete DMEM (without antibiotics). Diluted compounds were then added to the cells at a final concentration of 50 µg/ml to 0.78 µg/ml. The plate was then incubated for 48 hours at 37 ^°^C and 5% CO_2_. On Day 4, a stock solution of MTT (Sigma) was freshly prepared in sterile 1X PBS (Himedia) at a concentration of 20 mg/ml. MTT was further diluted to 5 mg/ml in complete DMEM (without antibiotics), followed by the addition of 20 µl of this diluted MTT into each well and incubated for 4 hours. After 4 hours, the media was completely removed and 100 µl of absolute DMSO was added to dissolve the insoluble formazan crystals formed by the action of mitochondrial dehydrogenase of live cells on soluble MTT (yellow). After two hours of the dissolution of formazan crystals, purple colored solution was analyzed by taking absorbance using a microplate (ELISA) reader at 570 nm and 620 nm using M4 SpectraMax® (Molecular Devices LLC). Deeper the purple color, the greater is the cell viability. Percentage cytotoxicity was calculated as described previously, using the following formula:

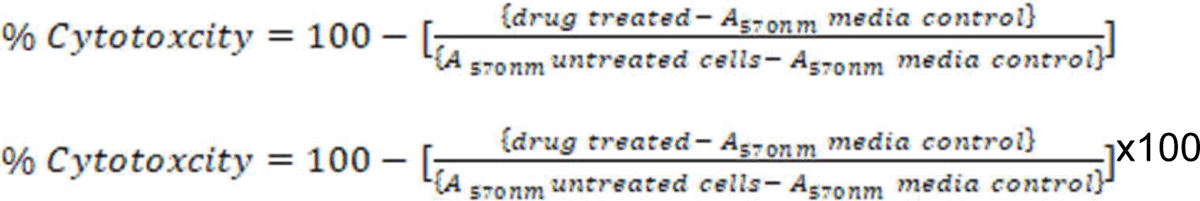

### Determination of Intracellular anti-mycobacterial activity of the shortlisted compounds in the THP-1-derived macrophages

Shortlisted compounds were tested for their intracellular anti-mycobacterial activity by methods as described previously ^61^ with minor modification. Briefly, the THP-1 cell line (human monocytic leukaemia cell line) was maintained in RPMI 1640 with 2 mM L-glutamine, 25 mM HEPES buffer, and 2 gm/liter sodium bicarbonate (Himedia) supplemented with 10% fetal bovine serum and 1X Pen Strep (Penicillin and streptomycin solution) at 37^°^ C and 5% CO_2._ In a 48-well plate (Thermo Scientific™ Nunc), 3 ×10^4^ THP-1 cells were seeded and differentiated by the addition of 5 ng/ml PMA (Sigma) in complete RPMI media (Himedia) (with 1X Pen Strep) as described above. Differentiated macrophages were then infected with *M. tb.* (H37Rv) at a multiplicity of infection (MOI) of 1:5 (1 macrophage: 5 bacteria) for 4 hours. Extracellular non-phagocytosed bacteria were removed by washing the infected cells three times with warm DPBS. The shortlisted compounds or first-line anti-TB drugs were prepared as stock containing 5X of the desired concentration in incomplete RPMI media (without FBS and without Pen Strep). For treatment, DPBS was removed and 400 µl of incomplete media (without FBS and without Pen strep) was added to each well, followed by the addition of 100 µl of first-line anti-TB drug or short-listed compounds at 5X of the desired concentrations, to obtain a final volume of 500 µl/well. Plates were then incubated at 37^°^ C and 5% CO_2_ for 5 days.

On day 5, infected macrophages were lysed using 0.025% SDS (sterile) and CFU plating was done on 7H11 agar plates. The plates were incubated at 37 ^°^C for 3-4 weeks and colonies were counted for CFU determination.

### Determination of anti-mycobacterial activity of drugs using nutrient starvation model of dormancy by most probable number (MPN) method

Shortlisted compounds were tested for their anti-mycobacterial activity employing nutrient starvation dormancy model using MPN assay as described previously^62^ and in **Supplementary Figure 8**. To mimic the dormant conditions, *M. tuberculosis* H37Rv was first grown in nutrient rich (7H9 supplemented with OADC and 0.05% tween 80) media for 7 days until an OD_600nm_ of 0.4-0.8. On day 8, the bacterial culture was centrifuged, washed twice with sterile PBS (Himedia) and then re-suspended in 10 mL PBS in sealed tubes. The sealed tubes were incubated at 37^°^C with 5% CO_2_ in humid and stationary condition (without shaking) for six weeks. Following 6 weeks of starvation, 200 µl of nutrient-starved cultures were taken in a microfuge tubes and treated with 10 µM of the first-line TB drugs, rifampicin and isoniazid and three of our shortlisted compounds for 7 days at 37 ^°^C with 5% CO_2_ in humid and stationary condition (without shaking). Untreated starved cells were similarly incubated as a control group. Post 7 weeks of starvation (including 1 week of drug treatment) cells were ten-fold serially diluted (10 to10^6^) in complete 7H9 media in microfuge tubes. From each serial dilution, 50 µl of diluted cultures were plated into a 48-well plate in triplicates in a 450 µl of 7H9 (complete) making a total volume of 500 µl/well. The plates were then incubated at 37 ^°^C with 5% CO_2_ in humid and stationary condition for 2-3 weeks (until bacterial growth is observed in most diluted untreated culture). Following this, MPN of bacterial cells was calculated. Briefly, as described in **Supplementary Figure 8**, bacterial growth in three consecutive dilutions (10^-5^, 10^-6^ and 10^-7^) was noted. *M. tuberculosis* viability was calculated as mean MPN/ml. The data is represented as the mean of n = 3 biological replicates with 95% confidence interval as per the three-tube MPN table^62^. The MPN methods have recently been applied for preclinical evaluation of anti-TB drugs using mouse tissues as well^63^.

## Statistical analysis

Data was analyzed using Microsoft excel, GraphPad Prism Software Version 5.0 using Student’s t-test and one-way ANOVA, wherever applicable.

**List of Figures, Tables and Supplementary Materials Figures:** Figures 1-9

**Tables:** Table 1-3

**Supplementary Materials:** Supplementary Figures 1-10; Supplementary Tables 1-8 and Supplementary Methods

## Acknowledgements

RJD is thankful to Birla Institute of Science and Technology (BITS) Pilani, Hyderabad campus for their funding support through intramural funding under Research Initiation Grant. RJD and DS are thankful for generous funding support from BITS Pilani Hyderabad campus, Centre for Human Disease Research (CHDR) for providing research funding and fellowship to MS. RJD is also thankful to the Department of Biotechnology (DBT) for supporting research through Ramalingaswami Re-entry fellowship (BT/RLF/Re-entry/18/2016). RJD is thankful to the overall infrastructure support by Department of Biological Sciences, BITS Pilani Hyderabad. MS is thankful to Lady Tata Memorial Trust for junior research fellowship. AD, NK and LA are thankful to the Council for Scientific and Industrial Research (CSIR) for the research fellowship. NA is thankful to the Department of Biotechnology (DBT), Ministry of Science and Technology, Govt. of India for funding support (BT/PR25690/GET/119/142/2017). SC thankfully acknowledges the Ramalingaswami Fellowship (BT/RLF/Reentry/09/2019).

Manish Bansal and Shiwani Goswami from THSTI are acknowledged for their technical assistance during construction of CRISPRi tools. Prof. William R. Jacobs, Jr at Albert Einstein College of Medicine, New York, USA is deeply acknowledged for providing mc^2^7902 strain utilised by RJD and NA for construction of knockdown strains. Dr. Bappaditya Dey, National Institute of Animal Biotechnology, Hyderabad is acknowledged for his help with MST analysis. Dr. DV Sai Prasad and Dr. Vagolu Siva Krishna, Dept of Pharmacy, BITS Pilani Hyderabad Campus are highly acknowledged for their help during compound characterization and *in silico* analysis, respectively. Jyothi Kumari, Department of Pharmacy, BITS Pilani Hyderabad is acknowledged for her technical support. Central Analytical Laboratory (CAL), Department of Science and Technology Fund for Improvement of S&T Infrastructure (DST-FIST) facilities at Department of Biological Sciences and Department of Pharmacy at BITS Pilani Hyderabad campus are highly acknowledged for provision of various central facilities and instruments used during the study. High Performance Computing Laboratory at BITS Pilani Hyderabad campus is highly appreciated for provision of central computational resources.

## Author Contributions

MS and RJD conceived and designed all the experiments, analyzed the data and designed the figures related to drug-based inhibition, protein assays, analysis of the protein structures and drug interactions at the binding site. AD and RJD conceived and designed experiments related to knockdown strains, analyzed the data and designed the figures. MS, AD and RJD wrote the manuscript. RJD and DS obtained the funding for the study. MS and AV performed the bioinformatic analysis for evolutionary conservation of the protein target. SC supervised analysis of the protein structures and drug interactions at the binding site and edited the manuscript. NK contributed to the MST assay. LA and NA designed the CRISPRi strategy for gene knockdown. NA also contributed to editing of the manuscript. DS contributed to design and synthesis scheme of the compound library and supervised *in silico* analysis and contributed to the writing of the manuscript. DS and RJD provided overall supervision during the study.

## Data availability statement

All data that supports the findings of this study are available from corresponding author upon reasonable request.

