## Supplementary Material for "Elucidation of the essentiality of lumazine synthase (RibH) for *Mycobacterium tuberculosis* survival and discovery of potent inhibitors for enhanced antimycobacterial therapy"

**Supplementary Figure 1-10**

**Supplementary Tables 1-9**

**Supplementary Methods**

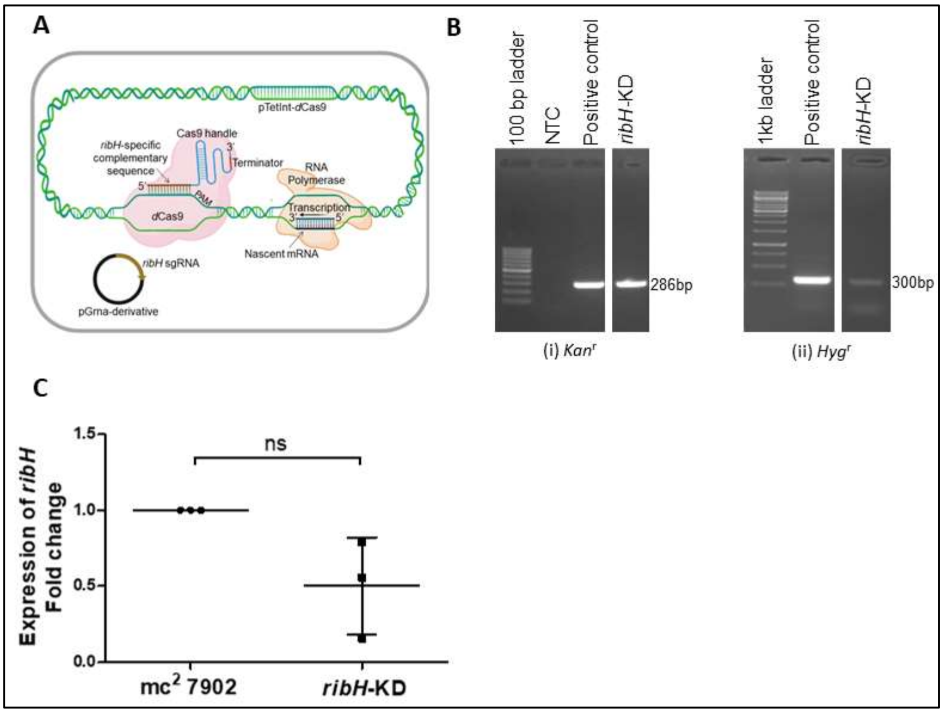


**Supplementary Figure 1. Conditional silencing or *ribH* by CRISPRi**. A. CRISPRi design strategy. *ribH* gene (*Rv1416*) specific complementary oligonucleotides were designed and cloned in *Hyg*r pGrna plasmid at *Sph*I-*Acl*I sites, downstream to the tetracycline-inducible promoter. The recombinant pGrna plasmid was then electroporated in to *M. tb* mc$^2$7902 strain having integrative plasmid pTetInt-dCas9 (*Kan*r). Following electroporation, recombinants were selected on kanamycin and hygromycin and the mutants were screened for *Kan*r and *Hyg*r specific gene primers. To achieve the desired level of suppression, the resulting strain was treated with Anhydrotetracycline (Atc). B. Verification of knockdown strains. Image shows agarose gel electrophoresis image showing the colony PCR for verification of *ribH*-KD strain of *M. tb* for (i) *Kan*r -286bp (ii) *Hyg*r -300bp C. Gene expression studies to determine the effect of induction with 50ng/ml of ATc on *ribH*-KD. Quantitative RT-PCR was used to validate CRISPRi-mediated silencing of the gene *ribH* with 50ng/ml ATc induction. Data (n=3) shows levels of ribH transcripts in *ribH*-KD strain. Data was analysed using Student's t-test and *p* values were determined.

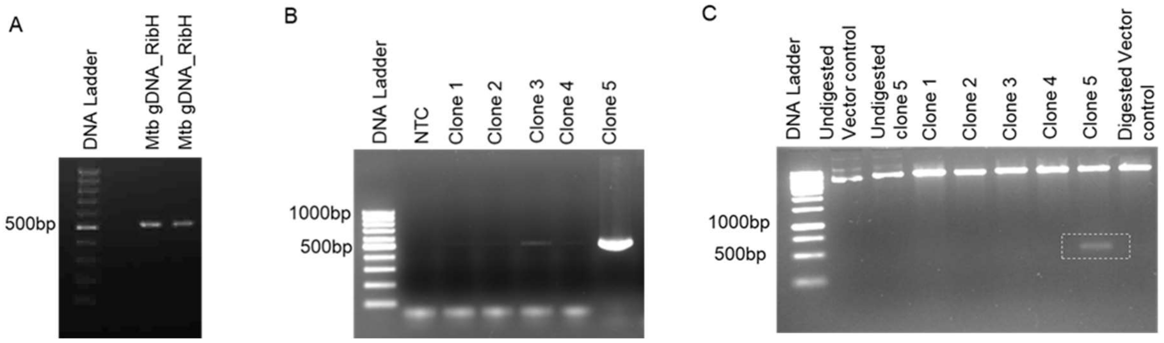


**Supplementary Figure 2. Cloning of *M. tb ribH* (*Rv1416*) in pet28a**. A. Gene encoding lumazine synthase (*ribH*, *Rv1416*) was PCR amplified using *H37Rv* chromosomal DNA as template and oligonucleotides (5' GGGGCATATGAAGGGTGGCGCCGGGGT 3') as forward primer and (5'GGGGCTCGAGTAGTCACGAGTGAGCGCGCAGCT 3') as reverse primer. The amplified PCR product was digested using appropriate restriction enzymes (*Nde* **I**, *Xho* **I**) and cloned into expression vector *pET28*a. B. The clone was confirmed using PCR amplification and C. restriction digestion and sequencing.

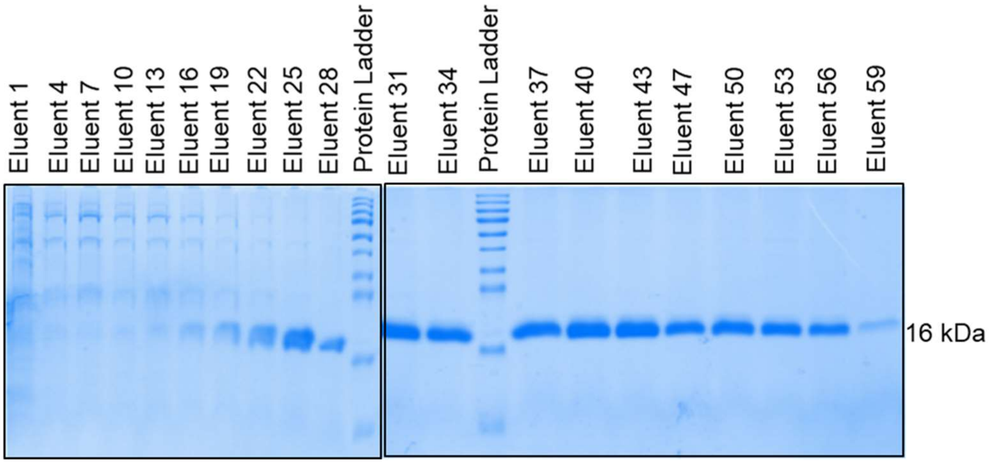


**Supplementary Figure 3. RibH protein expression and purification.** *E. coli* strain BL21(DE3) was transformed with pet28a.*ribH*. Clones were selected on kanamycin. Protein expression was induced with the help of IPTG as described in supplementary methods. Protein expression was analysed using SDS-PAGE for cultures induced with 0.2 mM IPTG concentration at 16° C overnight incubation. Cell pellets were lysed and protein was purified in native condition. 16 kDa RibH protein was purified by Ni-NTA affinity purification and the cleanest fractions were pooled for further dialysis and binding assays. The pictogram depicts the Coomassie brilliant blue stained SDS-PAGE

A

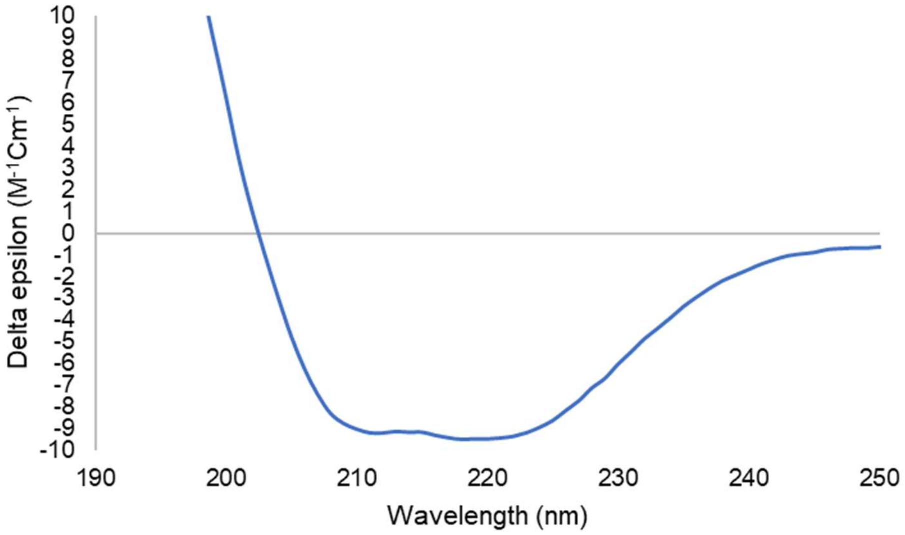


B

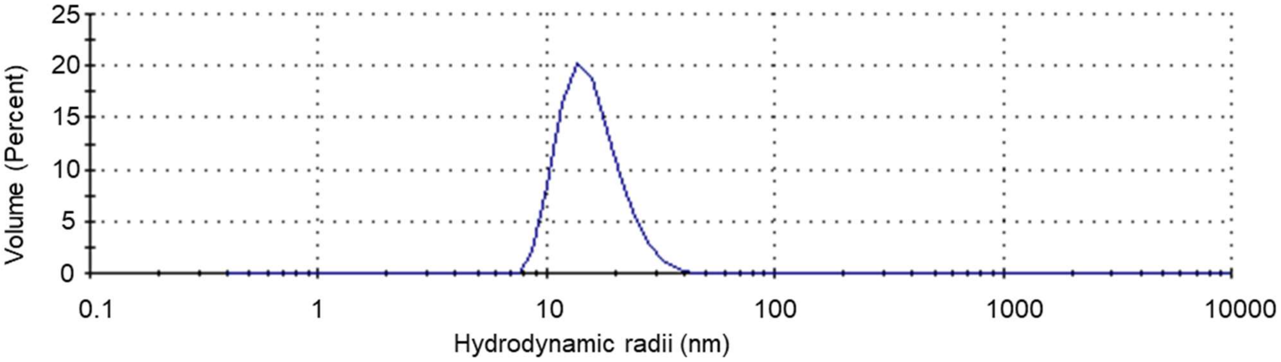


**Supplementary Figure 4. RibH protein characterization post-purification.** A. A far-UV CD spectrum was recorded for the purified *M. tb* RibH (in 50mM potassium phosphate buffer, pH 7.0) to characterize the secondary structure content of the protein. RibH protein consists of 61.9 % helices, 8.2% beta strands, 2.8% turns and 27.1% others. B. Dynamic light scattering (DLS) measurement of the purified RibH protein was assessed to determine the hydrodynamic radii (10nm).

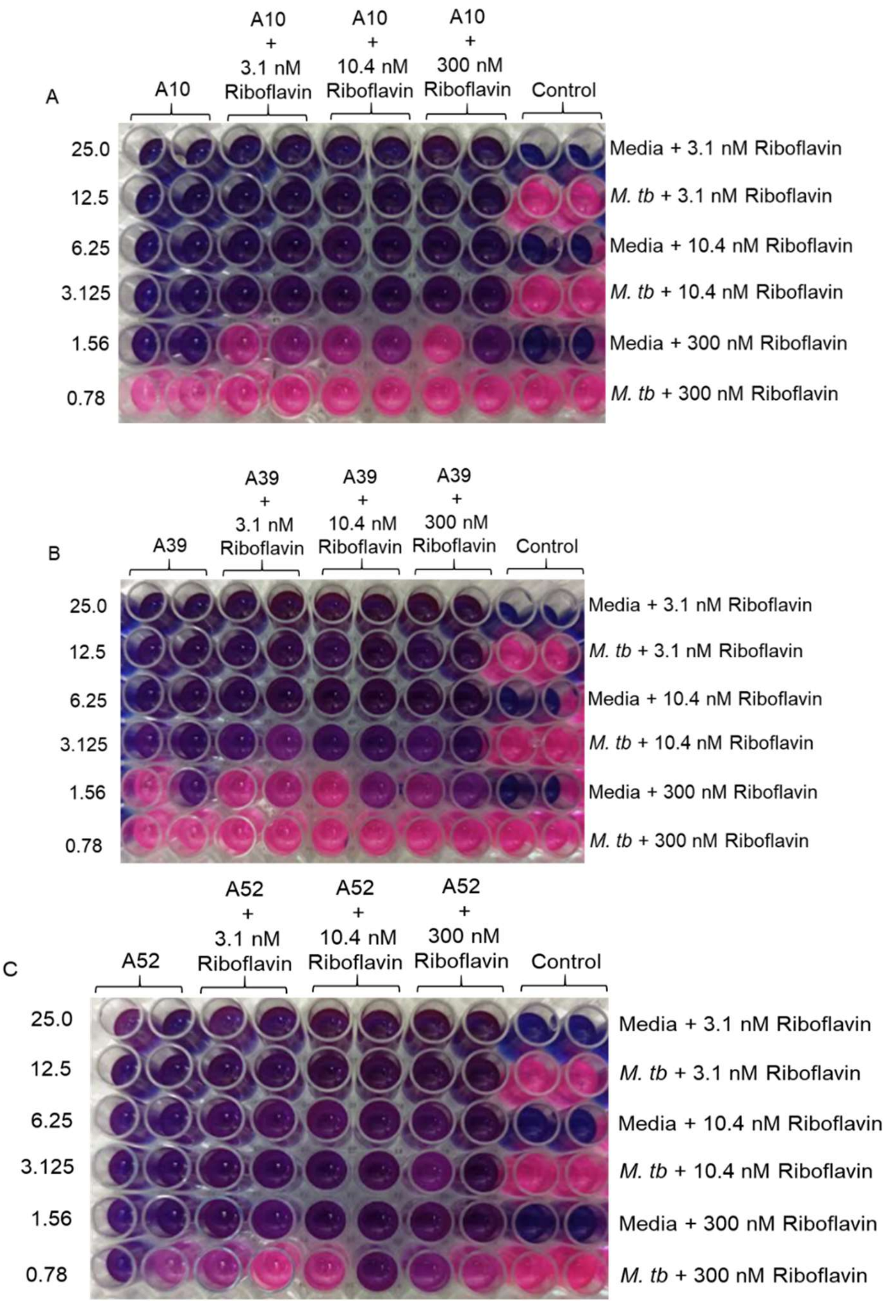


**Supplementary Figure 5. Anti-mycobacterial activity of short-listed compounds in combination with various concentrations of riboflavin.** The figure depicts the representative pictograms obtained on screening of anti-mycobacterial activity of three of the shortlisted compounds against *M. tuberculosis H37Rv* in combination with riboflavin. Riboflavin was tested at three different concentrations (3.1 nM, 10.4 nM and 300 nM) based on the range of plasma riboflavin concentrations in human population. In our assay conditions, Riboflavin was tested in combination with various concentrations of A10 (A), A39 (B) and A52 (C). A. A10 was tested alone at various concentrations between 25-0.78 µg/ml (two-fold serial dilution) and in combination with various concentrations of riboflavin (3.1 nM, 10.4 nM and 300 nM). In combination with riboflavin A10 shows an MIC of 3.125-1.56 µg/ml at all riboflavin concentrations. B. A39 was tested alone at various concentrations between 25-0.78 µg/ml (two-fold serial dilution) and in combination with various concentrations of riboflavin (3.1 nM, 10.4 nM and 300 nM). In combination with riboflavin A39 shows an MIC of 3.125 µg/ml at all riboflavin concentrations C. A52 was tested alone at various concentrations between 25-0.78 µg/ml (two-fold serial dilution) and in combination with various concentrations of riboflavin (3.1 nM, 10.4 nM and 300 nM). In combination with riboflavin A52 shows an MIC of 1.56-0.78 µg/ml at

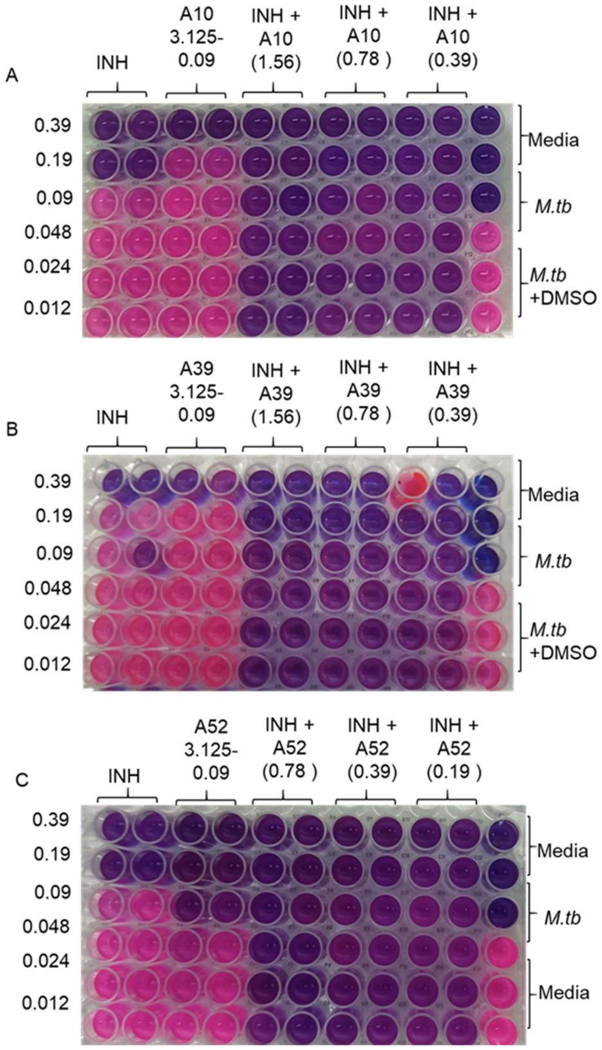


**Supplementary Figure 6. Anti-mycobacterial activity of short-listed compounds in combination with first-line anti-TB drug isoniazid.** The figure depicts the representative pictograms obtained on screening of anti-mycobacterial activity of three of the shortlisted compounds against *M. tuberculosis H37Rv* in combination with isoniazid by microplate Alamar blue assay (MABA). The assay was performed at least three times with three technical replicates in each case. In each case (A, B, and C) isoniazid was tested at various concentrations between 0.39 µg/ml to 0.012 µg/ml (two-fold serial dilution) either alone or in combination with various concentrations of A. A10, B. A39 and C. A52. These shortlisted compounds were either tested alone at various concentrations between 3.125-0.09 µg/ml (two-fold serial dilution) or at their respective MICs and sub-MIC concentrations in combination with various concentrations of isoniazid (MIC to the sub-MIC range). A synergistic enhancement in antimycobacterial activity of rifampicin and shortlisted compounds was observed as described in Table 1.

A

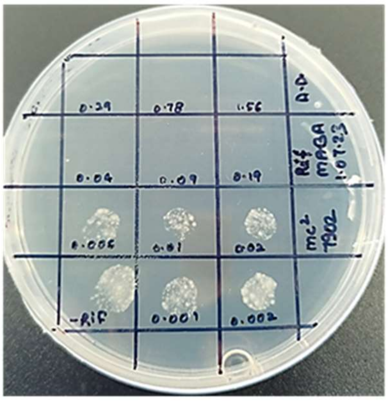


*mc*[2] *7902*
+ Rifampicin
(1.56 - 0.001 µg/ml)

B

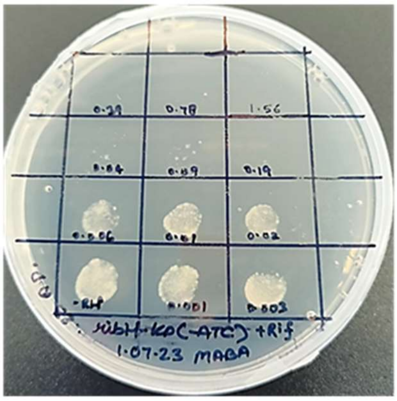


*ribH*-KD (-ATc)
+ Rifampicin
(1.56 - 0.001 µg/ml)

C

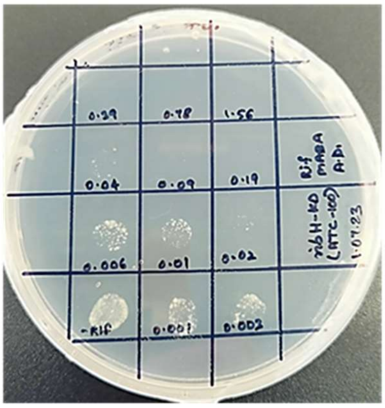


*ribH*-KD ( + 100 ng/ml ATc)
+ Rifampicin
(1.56 - 0.001 µg/ml)

**Supplementary Figure 7. Enhanced sensitivity of *ribH*- KD towards rifampicin** The figure depicts the representative pictograms obtained on screening of anti-mycobacterial activity of rifampicin against A. *M. tb mc*$^2$*7902* B. *ribH*-KD (-ATc) and C. *ribH*-KD (+100 ng/ml ATc). The respective strains were incubated in presence of various concentrations of rifampicin (1.56 - 0.001 µg/ml) in a 96-well plate for 7 days, following which, an equal volume was plated onto 7H11 Agar plates (with PLA supplements). While all the strains show similar growth in absence of rifampicin, *ribH*-KD strain (+ATc) shows much higher sensitivity to rifampicin as evident from significantly reduced growth on treatment with rifampicin at concentrations much below MIC (0.02 – 0.001 µg/ml). In comparison, parent strain and *ribH*-KD (-ATc) show comparable growth on exposure to rifampicin.

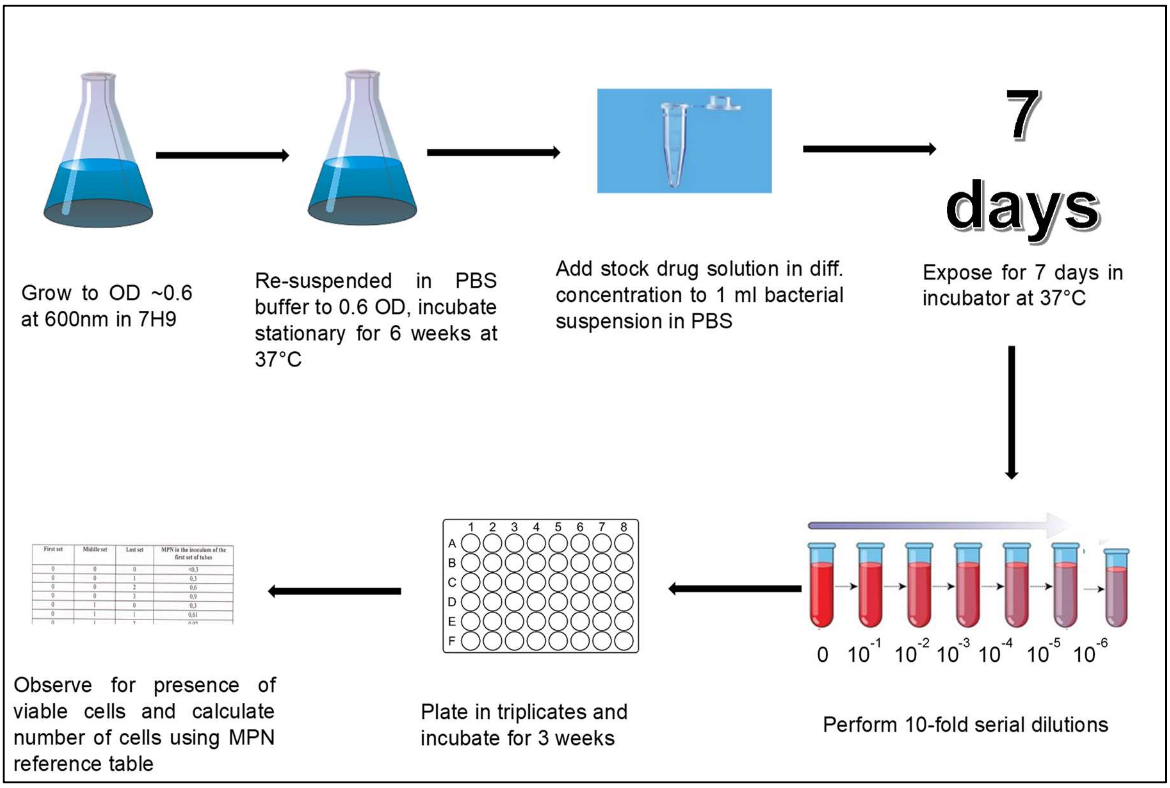


**Supplementary Figure 8. Nutrient starvation (an in vitro dormancy/non-replicating persistence) model and MPN assay design**

To mimic the dormant conditions, *M. tuberculosis H37Rv* is first grown in nutrient-rich (7H9 supplemented with OADC and 0.05% tween 80) media for 7 days until an $OD_{600}$ of 0.4-0.8. On day 8, the bacterial culture is centrifuged, washed with sterile PBS, and then re-suspended in PBS in sealed tubes. The sealed tubes are incubated at 37°C with 5% $CO_2$ in humid and stationary conditions for six weeks. Following 6 weeks of starvation, 200 µl of nutrient-starved cultures are taken in a microfuge tube and treated with drugs for 7 days. Untreated starved cells are similarly incubated as a control group. Post 7 weeks of starvation (including 1 week of drug treatment) cells are ten-fold serially diluted (10 to$10^{-6}$) in complete 7H9 media in microfuge tubes. From each serial dilution, 50µl of diluted cultures are plated into a 48-well plate in triplicates in a 450 µl of 7H9 (complete) making a total volume of 500 µl/well. The plates are then incubated 37°C with 5% $CO_2$ in humid and stationary condition for 2-3 weeks (until bacterial growth is observed in the most diluted untreated culture). Following this, MPN of bacterial cells is calculated. Bacterial growth in three consecutive dilutions ($10^{-5}$, $10^{-6}$, and $10^{-7}$) is noted. *M. tuberculosis* viability is calculated as mean MPN/ml. The CFU data (n = 3) is represented as the geometric mean (with 95%CI) as per the three-tube MPN table (See Figure 9).

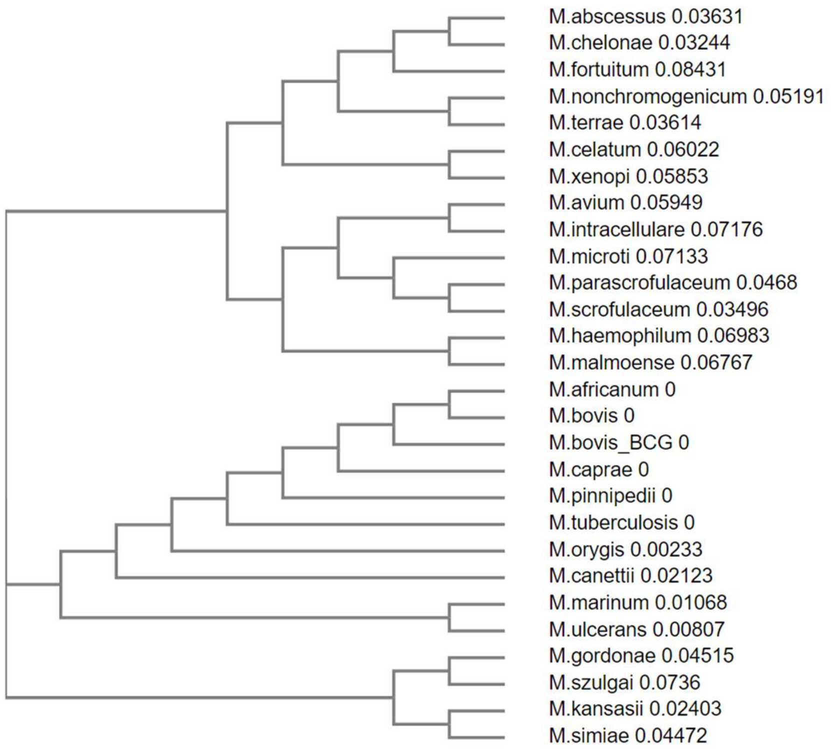


**Supplementary Figure 9. Phylogenetic analysis of RibH protein sequence across various mycobacterial species belonging to *M. tb* complex and non-tuberculous mycobacteria.** The phylogenetic tree was constructed for the RibH protein sequences of various mycobacterial species using T-Coffee. The scores mentioned in the tree are calculated by first calculating pairwise identity between two sequences. The identity between two sequences is then converted to a measure of evolutionary distance.

**Supplementary Figure 10. Sequence alignment of lumazine synthases from different mycobacterial species against the sequence of *M. tuberculosis* (strain ATCC 25618 / *H37Rv*).**

```
Mycobacterium tuberculosis (strain ATCC 25618 / H37Rv) RibH sequence

MKGGAGVPDLPSLDASGVRLAIVASSWHGKICDALLDGARKVAAGCGLDDPTVVRVLGAIEIPVVAQELARNHDAVVAL
GVVIRGQTPHFDYVCDAVTQGLTRVSLDSSTPIANGVLTTNTEEQALDRAGLPTSAEDKGAQATVAALATALTLRELRA
HS

CLUSTAL W (1.83) multiple sequence alignment
M.abscessus          MS-----------------------GDGIPSLSVGDASGISLAIVASTWH
M.africanum          MKG----------------------GAGVPDLPSLDASGVRLAIVASSWH
M.avium              MSP----------------------AAGVPEMPALDASGVRLGIVASTWH
M.bovis              MKG----------------------GAGVPDLPSLDASGVRLAIVASSWH
M.bovis_BCG          MKG----------------------GAGVPDLPSLDASGVRLAIVASSWH
M.canettii           MTG----------------------GAGVPDLPSLDASGVRLAIVASSWH
M.caprae             MKG----------------------GAGVPDLPSLDASGVRLAIVASSWH
M.celatum            MSP----------------------AAGVPDMPALDASSVKLAIVASTWH
M.chelonae           MS-----------------------GEGIPSLTVGDASSLSLAIVASTWH
M.fortuitum          MS-----------------------AHGVPDLPQVDASNVKLAIVASTWH
M.gordonae           MSG----------------------GAGVPEMPVLDASGLRLAIVASTWH
M.haemophilum        MSG----------------------GAGVPEVPAIDASGLRLGIVASTWH
M.intracellulare     MSP----------------------AEGVPEVPPLDASGLRLALVASTWH
M.kansasii           MSG----------------------GAGVPDIPALDASGVRLAIVASTWH
M.malmoense          MSG----------------------GAGVPEIPALDASGLRLGIVASTWH
M.marinum            MSG----------------------GAGIPDVPAFDASDVRLAIVASTWH
M.microti            MS-----------------------GAGIPDLGELDASGLRLGIVASTWH
M.nonchromogenicum   MSP----------------------AAGVPDMPALDASDVRLAIVASTWH
M.orygis             ----------------------------MPDLPSLDASGVRLAIVASSWH
M.parascrofulaceum   MEPCDRRRQDDAVRVSPAWGTAQVSGSGEPEIPALDATGLRLGIVASTWH
M.pinnipedii         MKG----------------------GAGVPDLPSLDASGVRLAIVASSWH
M.scrofulaceum       MS-----------------------GSGEPDIPALDASGLRVGIVASTWH
M.simiae             MSG----------------------GAGVPDIPALDASGVRLAIVASTWH
M.szulgai            MSG----------------------GAGVPEMPVLDAADVQLAIVASTWH
M.terrae             MS-----------------------GTGVPDMPALDASGVRLAIVASTWH
M.tuberculosis       MKG----------------------GAGVPDLPSLDASGVRLAIVASSWH
M.ulcerans           MSG----------------------GAGIPDVPAFDASGVRLAIVASTWH
M.xenopi             MSP----------------------AAGMPELPALDASGLKLAIVASTWH
                                                  *.:   **:.: :.:***:**


M.abscessus          DQICTALLEGAQRVATEAGIDRPTVVRVLGAIEIPVVAQALARTHDAVVA
M.africanum          GKICDALLDGARKVAAGCGLDDPTVVRVLGAIEIPVVAQELARNHDAVVA
M.avium              SRICDALLAGARKVAADSGVENPTVVRVLGAIEIPVVAQELARNHDAVVA
M.bovis              GKICDALLDGARKVAAGCGLDDPTVVRVLGAIEIPVVAQELARNHDAVVA
M.bovis_BCG          GKICDALLDGARKVAAGCGLDDPTVVRVLGAIEIPVVAQELARNHDAVVA
M.canettii           GKICDALLDGARNVAAGCGLDDPTVVRVLGAIEIPVVAQELARNHDAVVA
M.caprae             GKICDALLDGARKVAAGCGLDDPTVVRVLGAIEIPVVAQELARNHDAVVA
M.celatum            TTICDALLGGARKTAAEWGVDDPTVVRVLGAIEIPVVAQALAREHDAVVA
M.chelonae           DQICTALLEGAKRVAADAGIDRPTVVRVLGAIEIPVVAQALARTHDAVVA
M.fortuitum          TQICDALLDGARKVAADAGIAEPTVVRVLGAIEIPVVAQALAATHDAVVA
M.gordonae           DKICDALLAGARKVATDSGIDTPTVVRVLGAIEIPVVAQELARNHDAVVA
M.haemophilum        GRICDALLAGARKVAANSGVDNPTVVRVLGAIEIPVVAQELARTHDAVVA
M.intracellulare     SEICDALLAGASKVASESGVDDPTVVRVIGAIEIPVVAQELARNHDAVVA
M.kansasii           SQICDALLAGARKVAADSGIDNPTVVRVLGAIEIPVVAQELARQHDAVVA
M.malmoense          GQICDALLAGARKMAAESGVENPTVVRVLGAIEIPVVAQQLARDHDAVVA
M.marinum            TKICDALLAGARNTAADSGIDNPTVVRVLGAIEIPVVAQELTRNHDAVVA
M.microti            STICDALLAGAERVVARSGVDNPTVVRVHGAIEIPVVAQELARNHDAVIA
M.nonchromogenicum   TRICDALLAGARRVAAEAGIPEPTVVRVLGAIEIPVVTQEVARNHDAVVA
M.orygis             GKICDALLDGARKVAAGCGLDDPTVVRVLGAIEIPVVAQELARNHDAVVA
M.parascrofulaceum   GKICEALLQGARRVAAESGIDNPTVVRVLGAIEIPVVAQELTRNHDAVIA
M.pinnipedii         GKICDALLDGARKVAAGCGLDDPTVVRVLGAIEIPVVAQELARNHDAVVA
M.scrofulaceum       SKICEALLDGARRVAAESGIDNPTVVRVLGAIELPVVAQELARNHDAVIA
M.simiae             AEICDALLAGARKVATESGIDNPTVARVLGAIEIPVLAQELARSHDAVVA
M.szulgai            TEVCDALLAGALKVAADSGIPSPTVVRVLGAIEIPVVVQELARSHDAVVA
M.terrae             TRICDALLAGARKVAADAGVADPTVVRVLGAIEIPVVAQEVARNHDAVVA
M.tuberculosis       GKICDALLDGARKVAAGCGLDDPTVVRVLGAIEIPVVAQELARNHDAVVA
M.ulcerans           TKICDALLAGARNTAADSGIDNPTVVRVLGAIEIPVVAQELTRNHDAVVA
M.xenopi             TEICDALLTGARKTAAEWGIDDPTVVRVLGAIEIPVVAQELARSHDAVVA
                       :* *** ** . .:  *:  ***.** ****:**:.* ::  ****:*
```

```
M.abscessus           LGVVIQGETPHFGYVCDAVTTGLTRVSLDTSTPVANGVLTVNNEQQAIDR
M.africanum           LGVVIRGQTPHFDYVCDAVTQGLTRVSLDSSTPIANGVLTTNTEEQALDR
M.avium               LGVVIRGQTPHFEYVCDAVTQGITRVSLDASTPVANGVLTTDNEQQALDR
M.bovis               LGVVIRGQTPHFDYVCDAVTQGLTRVSLDSSTPIANGVLTTNTEEQALDR
M.bovis_BCG           LGVVIRGQTPHFDYVCDAVTQGLTRVSLDSSTPIANGVLTTNTEEQALDR
M.canettii            LGVVIRGQTPHFDYVCDAVTQGLTRVSLDSSTPIGNGVLTTNTEEQALDR
M.caprae              LGVVIRGQTPHFDYVCDAVTQGLTRVSLDSSTPIANGVLTTNTEEQALDR
M.celatum             LGVVIRGQTPHFDYVCDAVTQGLTRVSLDASTPVANGVLTTNTEEQALDR
M.chelonae            LGVVIQGETPHFNYVCDAVTTGLTRVSLDTSTPVANGVLTVNNEQQALDR
M.fortuitum           LGVVIRGQTPHFDYVCDAVTQGLTRVSLDASTPVANGVLTTDNEAQALDR
M.gordonae            LGVVIRGETPHFDYVCDAVTQGLTRVSLDASTPVANGVLTTNTEAQALDR
M.haemophilum         LGVVIRGATPHFDHVCNSVTQGLTRVALDTSTPVGNGVLTTNTEEQALDR
M.intracellulare      LGVVIRGQTPHFEYVCDAVTQGLTRVSLDTSTPVANGVLTTDTEQQALDR
M.kansasii            LGVVIRGETPHFDYVCDAVTQGLTRVSLDESTPVANGVLTTNTEEQALDR
M.malmoense           LGVVIRGETPHFDYVCDSVTQGLTRVSLDACTPIGNGVLTTNTEEQALAR
M.marinum             LGVVIRGETPHFDYVCDAVTQGLTRVSLDSSTPVANGVLTTNSEEQALNR
M.microti             LGVVIRGGTPHFEYVCDAVTQGLTRVSLDASTPVANGVLTTNTEEQALDR
M.nonchromogenicum    LGVVIRGATPHFDYVCDAVTQGLTRVSLDAATPVANGVLTVNDEEQALDR
M.orygis              LGVVIRGQTPHFDYVCDAVTQGLTRVSLDSSTPIANGVLTTNTEEQALDR
M.parascrofulaceum    LGVVIRGETPHFSYVCDAVTQGLTRVALDASTPVANGVLTTNTEQQALDR
M.pinnipedii          LGVVIRGQTPHFDYVCDAVTQGLTRVSLDSSTPIANGVLTTNTEEQALDR
M.scrofulaceum        LGVVIRGGTPHFEYVCDAVTQGLTRVALDASTPVANGVLTTNTEQQALDR
M.simiae              LGVVIRGETPHFDYVCDAVTQGLTRVSLDASTPVANGVLTTNTEEQARDR
M.szulgai             LGVVIRGETPHFDYVCDAVTQGLTRVSLDESTPVGNGVLTTNTEEQALDR
M.terrae              LGVVIRGGTPHFDYVCDAVTQGLTRVSLDAATPVANGVLTVNDEQQALDR
M.tuberculosis        LGVVIRGQTPHFDYVCDAVTQGLTRVSLDSSTPIANGVLTTNTEEQALDR
M.ulcerans            LGVVIRGETPHFDYVCDVVTQGLTRVSLDSSTPVANGVLTTNSEEQALNR
M.xenopi              LGVVIRGETPHFDYVCDAVTQGLTRVSLDASTPVANGVLTTNTEEQARDR
                      *****:* **** :**: ** *:***:** .**:.*****.: * **  *

M.abscessus           AGLPGSAEDKGAQAAAALDTALTLRRLRQPW--A
M.africanum           AGLPTSAEDKGAQATVAALATALTLRELRAH---S
M.avium               AGLPDSAEDKGAQAAGAALSAALTLRELRAR---S
M.bovis               AGLPTSAEDKGAQATVAALATALTLRELRAH---S
M.bovis_BCG           AGLPTSAEDKGAQATVAALATALTLRELRAH---S
M.canettii            AGLPTSAEDKGAQATVAALATALTLRELRAQ---P
M.caprae              AGLPTSAEDKGAQATVAALATALTLRELRAH---S
M.celatum             AGLPDSAEDKGAQATAAALSTALTLRDLRAK---P
M.chelonae            AGLPESSEDKGAQAAAALDTALTLRRLRQPWTEA
M.fortuitum           AGLPDSTEDKGAQAAAALSTALTLRELRSK---A
M.gordonae            AGLPSSAEDKGAQAAAALTTALTLRDLRAQ---S
M.haemophilum         AGLPTSAEDKGAQAAAALTTALTLLNLRAR---T
M.intracellulare      AGLPESAEDKGAQATLAALTTALTLRELRAR---S
M.kansasii            AGLPTSAEDKGAQATAAALTTALTLRELRVQ---S
M.malmoense           AGLPASAEDKGAQAAGAALTAALTLRDLRAR---S
M.marinum             AGLPTSDEDKGAQATAAALTTALTLRELRAE---A
M.microti             AGLPTSAEDKGAQAAGAALTAALALRDLRAR---S
M.nonchromogenicum    AGLPGSTEDKGAQAAAALSTALTLRELRAA---P
M.orygis              AGLPTSAEDKGAQATVAALATALTLRELRAH---S
M.parascrofulaceum    AGLPESVEDKGAQAAAALTAALTLRDLRAR---P
M.pinnipedii          AGLPTSAEDKGAQATVAALATALTLRELRAH---S
M.scrofulaceum        AGLPESVEDKGAQAAGAALTAALTLRELRAR---S
M.simiae              AGLPTSAEDKGAQATAAALTTALALRELRAQ---S
M.szulgai             AGLPTSAEDKGAQATAAALATALTLRDLRAQ---S
M.terrae              AGLPGSAEDKGAQAAAALSTALTLRDLRAA---L
M.tuberculosis        AGLPTSAEDKGAQATVAALATALTLRELRAH---S
M.ulcerans            AGLPTSDEDKGAQATAAALTTALTLRELRAE---S
M.xenopi              AGLPTSTEDKGAQATAALNTALTLRELRTR---S
                      **** * *******: *** :**:*  **
```

**Supplementary Figure 10.** Sequence alignment of lumazine synthases from different mycobacterial species against the sequence of *M. tuberculosis* (strain ATCC 25618 / *H37Rv*). The sequence highlighted in yellow is that of RibH from *Mycobacterium tuberculosis* (strain ATCC 25618 / *H37Rv*). The residues highlighted in red indicated the ones present in the drug binding pocket. The residues highlighted in blue indicate the additional residues (along with the residues highlighted in red) present in the co-crystal ligand binding pocket. Multiple sequence alignment was performed using T-Coffee.

#### Supplementary Tables

**Supplementary Table 1. *In silico* molecular docking of various compounds against RibH.** The table depicts the docking scores, structure of the compounds, minimum inhibitory concentration (MIC) against *M. tuberculosis H37Rv*, and important amino acids involved in interaction with the concerned drug molecules.

| Sl. No. | Compound Code | Structure | Docking Score | MIC (µg/mL) | Amino acid residues interacting with ligand |
|---|---|---|---|---|---|
| 1 | TP6 | 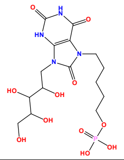 | - | Not applicable | SER 25, TRP 27, HIE 28, ILE 60, GLU 61, VAL 81, ILE 83, GLN 86, THR 87 |
| 2 | Riboflavin | 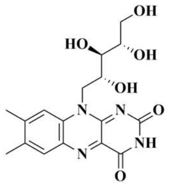 | - | Not applicable | HIS 28, ALA 59, GLU 61, VAL 81, ILE 83, ASN 114, LYS 138 |
| 3 | NR 353 (A52)[#] | 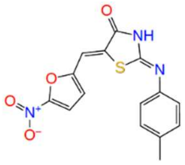 | -8.431 | 0.78 | SER 25, ALA 59, GLU 61, VAL 81, ASN 114 |
| 4 | NR-310 (A10)[#] | 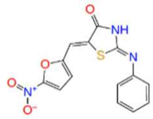 | -7.276 | 1.56 | ALA 59, GLU 61, VAL 81 |
| 5 | NR-340 (A39)[#] | 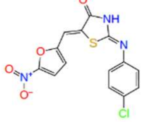 | -6.825 | 1.56 | ALA 59, GLU 61, VAL 81, THR 87, ASN 114 |
| 6 | 9F-Srihari*,[#] | 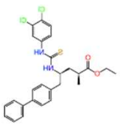 | -7.725* (-6.652[#]) | 3.125 | HIS 28, THR 87, HIE 89, ARG 128, LYS 138 |
| 7 | QNH-08*,[#] | 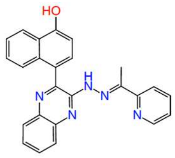 | -7.722* (-4.568[#]) | 3.125 | TRP 27, ILE 83, HIE 89, ARG 128, LYS 138 |
| 8 | T9*,[#] | 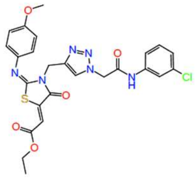 | -7.950* (-7.969[#]) | 6.25 | TRP 27, ALA 59, ILE 60, THR 87, GLU 136, LYS 138 |

| | | | | | |
|---|---|---|---|---|---|
| 9 | Brahm 2nd 15# | 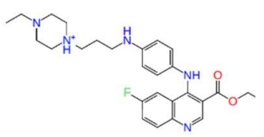 | -8.251 | 12.5 | TRP 27, ALA 59, HIE 89, GLU 136, LYS 138 |
| 10 | NR-341# | 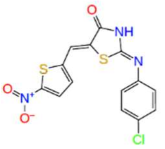 | -7.546 | 12.5 | TRP27, ALA 59, VAL 81, THR 87, LYS 138 |
| 11 | Boga-R-1000-299*,# | 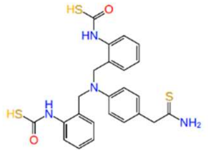 | -0.425* (-5.022#) | 12.5 | TRP 27, HIS 28, GLU 61, THR 87, LYS 138 |
| 12 | NR 302# | 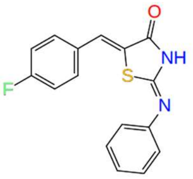 | -8.724 | 25 | TRP 27, ALA 59, VAL 81, |
| 13 | NR 317# | 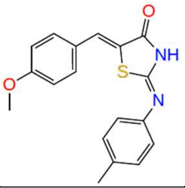 | -8.217 | 25 | ALA 59, VAL 81, ILE 83 |
| 14 | Brahm 2nd 22# | 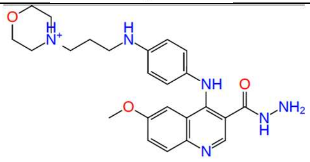 | -8.155 | 25 | TRP 27, ALA 59, ILE 83, HIE 89, ASN 114, GLU 136, LYS 138 |
| 15 | Srihari_28C*,# | 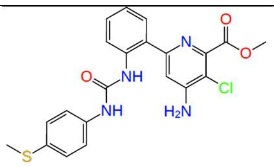 | -8.77* (-7.614#) | 25 | TRP 27, ILE 83, THR 87, ARG 128, GLU 136 |
| 16 | NIPER-TA-5*,# | 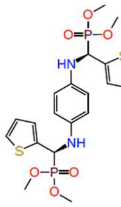 | -9.861* (-9.651#) | 50 | TRP 27, ILE60,THR 87, HIE 89, ARG 128, GLU 136 |
| 17 | Brahm_2nd 23# | 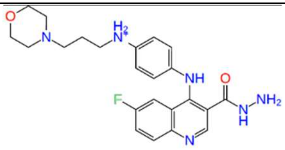 | -9.028 | 50 | TRP 27, ALA 59, HIE 89, ASN 114, GLU 136, LYS 138 |
| 18 | NR 336# | 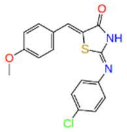 | -8.761 | 50 | ALA 59, VAL 81, ILE 83, THR 87 |

| | | | | | |
|---|---|---|---|---|---|
| 19 | GS 407# | 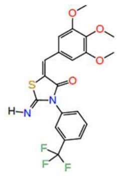 | -8.655 | 50 | ALA 59, THR 87, ARG 128 |
| 20 | Brahm 2nd 16# | 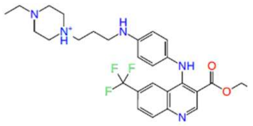 | -8.5 | 50 | TRP 27, ALA 59, HIE 89, GLU 136, LYS 138 |
| 21 | Brahm 2nd 7# | 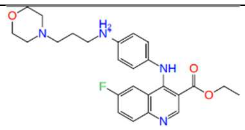 | -8.256 | 50 | TRP 27, ALA 59, HIE 89, GLU 136, LYS 138 |
| 22 | CLRI-ET3A*,# | 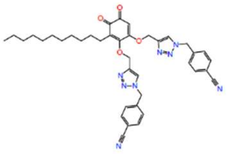 | -8.207* (-7.72#) | 50 | TRP 27, THR 87, GLN 86, ARG 128 |
| 23 | ASN 11105690* | 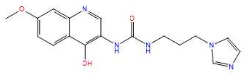 | -11.615 | >50 | TRP 27, ILE 60, VAL 81, ILE 83, GLU 122 |
| 24 | BAS 07023776* | 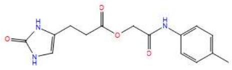 | -11.485 | >50 | ALA 59, ILE 83, GLN 86, THR 87 |
| 25 | BAS 14050851* | 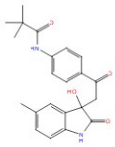 | -11.188 | >50 | TRP 27, HIS 28, ALA 59, ILE 60, ILE 83, LYS 138 |
| 26 | BAS 07805256* | 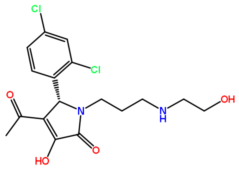 | -11.086 | >50 | TRP 27, ALA 59, GLU 61, VAL 81, ILE 83, HIE 89, ASN 114 |
| 27 | BAS 00728180* | 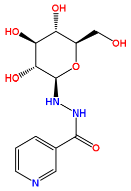 | -10.998 | >50 | TRP 27, ALA 59, ILE 83, THR 87, ARG 128, GLU 136 |
| 28 | BAS 00829582* | 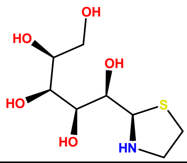 | -10.95 | >50 | ALA 59, ILE 60, GLU 61, ASN 114, LYS 138 |
| 29 | BAS 04239203* | 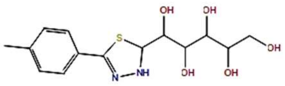 | -10.897 | >50 | TRP 27, ILE 83, GLY 85, THR 87 |

| | | | | | |
|---|---|---|---|---|---|
| 30 | BAS 00674022* | 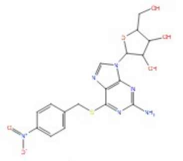 | -10.866 | >50 | SER 25, TRP 27, ALA 59, GLU 61, VAL 81, GLN 86, THR 87, ARG 128, LYS 138 |
| 31 | BAS 03571839* | 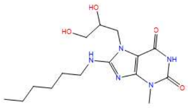 | -10.813 | >50 | TRP 27, HIS 28, ALA 59, VAL 81, ILE 83 |
| 32 | BAS 07805261* | 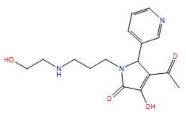 | -10.738 | >50 | GLU 61, VAL 81, ILE 83, HIE 89, ASN 114, LYS 138 |
| 33 | BAS 02327074* | 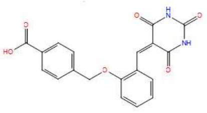 | -10.728 | >50 | TRP 27, THR 87, HIE 89, ASN 114, ARG 128, LYS 138 |
| 34 | ASN 11105540* | 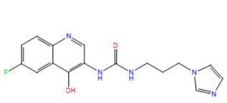 | -10.621 | >50 | TRP 27, VAL 81, ILE 83, GLU 124 |
| 35 | ASN02254483* | 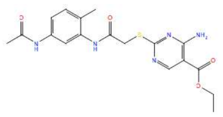 | -9.821 | >50 | TRP 27, ALA 59, VAL 81, ILE 83, ARG 128, GLU 136, LYS 138 |
| 36 | SF Srihari[#] | 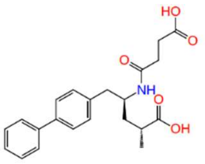 | -9.577 | >50 | TRP 27, ILE 83, THR 87, PHE 90 |
| 37 | NR 346[#] | 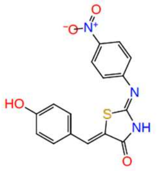 | -9.459 | >50 | ALA 59, VAL 81, THR 87, ASN 114 |
| 38 | ASN14403850* | 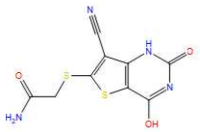 | -9.428 | >50 | VAL 81, ILE 83, GLN 86, THR 87, LYS 138 |
| 39 | SY-Srihari[*,#] | 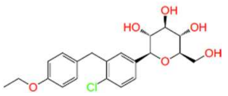 | -9.391* (-9.203[#]) | >50 | TRP 27, ALA 59, ILE 83, GLU 136, LYS 138 |
| 40 | ASN 03798609* | 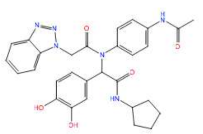 | -8.988 | >50 | TRP 27, HIS 28, ALA 59, VAL 81, HIE 89, ARG 128, GLU 136, LYS 138 |
| 41 | NR 322[#] | 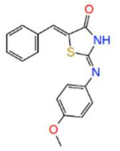 | -8.875 | >50 | TRP 27, ALA 59, VAL 81, ILE 83 |

| | | | | | |
|---|---|---|---|---|---|
| 42 | NIPER_TA_VAN* | 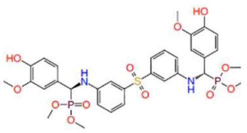 | -7.904*<br>-8.766# | >50 | TRP 27, ALA 59, ILE 60, VAL 81, ARG 128, GLU 136 |
| 43 | Srihari 26C# | 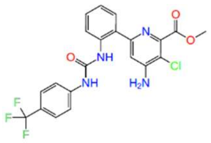 | -8.698 | >50 | TRP 27, ILE 83, THR 87, ARG 128, GLU 136 |
| 44 | NR 325# | 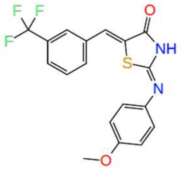 | -8.675 | >50 | ALA 59, VAL 81, ILE 83 |
| 45 | NR 347# | 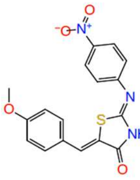 | -8.668 | >50 | ALA 59, VAL 81, ILE 83, GLY 85, ARG 128 |
| 46 | P10# | 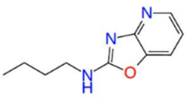 | -8.602 | >50 | ALA 59, ILE 60,GLU 61 |
| 47 | Srihari 24C# | 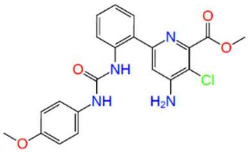 | -8.501 | >50 | TRP 27, ALA 59, ILE 83, GIY 85, THR 87, ARG 128 |
| 48 | NR 324# | 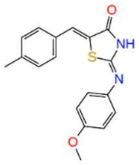 | -8.417 | >50 | ALA 59, VAL 81 |
| 49 | NR 304# | 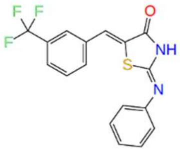 | -8.325 | >50 | ALA 59, VAL 81, ILE 83 |
| 50 | NR 314# | 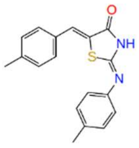 | -8.253 | >50 | TRP 27, ALA 59, VAL 81, ILE 83 |
| 51 | NR 303# | 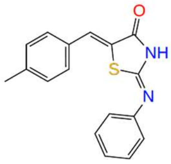 | -8.237 | >50 | ALA 59, VAL 81, ILE 83 |
| 52 | GP-226*,# | 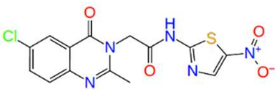 | -7.939*<br>(-8.179#) | >50 | TRP 27, ALA 59, GLY 85, THR 87, HIE 89, PHE 89, ARG 128 |

| | | | | | |
|---|---|---|---|---|---|
| 53 | ST Srihari# | 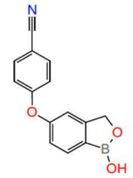 | -8.128 | >50 | TRP 27, ILE 60, GLY 85, THR 87, |
| 54 | Srihari-2M*,# | 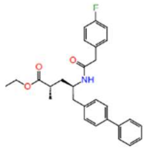 | -7.756* (-7.37#) | >50 | TRP 27, THR 87, HIE 89, ARG 128, LYS 138 |
| 55 | T8*,# | 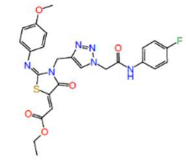 | -8.821* (-6.143#) | >50 | TRP 27, ALA 59, ILE 83, ARG 128 |
| 56 | B2-39-3*,# | 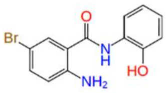 | -0.200* (-4.899#) | >50 | TRP 27, ALA 59, VAL 81, HIE 89, ASN 114 |
| 57 | RN-LR-17*,# | 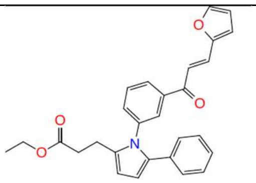 | -7.986* (-3.851#) | >50 | HIE 89, ASN 114, ARG 128, LYS 138 |

* Compounds screened by pharmacophore based virtual screening (PBVS)
### Compounds screened by Structure-based flexible docking
*,# Compounds screened by both pharmacophore based virtual screening (PBVS) and Structure-based flexible docking
-BITS in-house compound library was screened by 2 molecular docking methods and docking score outside bracket is obtained by e-pharmacophore based virtual screening and docking score inside the brackets denote those obtained by flexible docking.
-Compounds with code name ASN or BAS belong to ASINEX library (15) and rest of the compounds belong to BITS-in-house library.

**Supplementary Table 2: Input Protein Sequences used for various bioinformatic analysis**

| Sl. No. | Mycobacterial species | Lumazine synthase (RibH) protein sequence |
|---|---|---|
| 1 | *Mycobacterium tuberculosis (strain ATCC 25618 / H37Rv)* | >sp\|P9WHE9\|RISB_MYCTU OS=Mycobacterium tuberculosis (strain ATCC 25618 / H37Rv) OX=83332 GN=ribH PE=1 SV=1<br>MKGGAGVPDL PSLDASGVRL AIVASSWHGK ICDALLDGAR KVAAGCGLDD PTVVRVLGAI EIPVVAQELA RNHDAVVALG VVIRGQTPHF DYVCDAVTQG LTRVSLDSST PIANGVLTTN TEEQALDRAG LPTSAEDKGA QATVAALATA LTLRELRAHS |
| 2 | *Mycobacterium* abscessus | >sp\|B1MCA3\|RISB_MYCA9 OS=Mycobacteroides abscessus (strain ATCC 19977 / DSM 44196 / CCUG 20993 / CIP 104536 / JCM 13569 / NCTC 13031 / TMC 1543 / L948) OX=561007 GN=ribH PE=3 SV=1<br>MSGDGIPSLS VGDASGISLA IVASTWHDQI CTALLEGAQR VATEAGIDRP TVVRVLGAIE IPVVAQALAR THDAVVALGV VIQGETPHFG YVCDAVTTGL TRVSLDTSTP VANGVLTVNN EQQAIDRAGL PGSAEDKGAQ AAAAALDTAL TLRRLRQPWA |
| 3 | *Mycobacterium tuberculosis variant africanum* | >tr\|A0A120J0T0\|A0A120J0T0_MYCTX OS=Mycobacterium tuberculosis variant africanum OX=33894 GN=ribH PE=3 SV=1<br>MKGGAGVPDL PSLDASGVRL AIVASSWHGK ICDALLDGAR KVAAGCGLDD PTVVRVLGAI EIPVVAQELA RNHDAVVALG VVIRGQTPHF DYVCDAVTQG LTRVSLDSST PIANGVLTTN TEEQALDRAG LPTSAEDKGA QATVAALATA LTLRELRAHS |
| 4 | *Mycobacterium bovis* | >sp\|P66035\|RISB_MYCBO OS=Mycobacterium bovis (strain ATCC BAA-935 / AF2122/97) OX=233413 GN=ribH PE=3 SV=1<br>MKGGAGVPDL PSLDASGVRL AIVASSWHGK ICDALLDGAR KVAAGCGLDD PTVVRVLGAI EIPVVAQELA RNHDAVVALG VVIRGQTPHF DYVCDAVTQG LTRVSLDSST PIANGVLTTN TEEQALDRAG LPTSAEDKGA QATVAALATA LTLRELRAHS |
| 5 | *Mycobacterium tuberculosis variant bovis BCG* | >tr\|A0A0K2HV99\|A0A0K2HV99_MYCBI OS=Mycobacterium tuberculosis variant bovis BCG OX=33892 GN=ribH PE=3 SV=1<br>MKGGAGVPDL PSLDASGVRL AIVASSWHGK ICDALLDGAR KVAAGCGLDD PTVVRVLGAI EIPVVAQELA RNHDAVVALG VVIRGQTPHF DYVCDAVTQG LTRVSLDSST PIANGVLTTN TEEQALDRAG LPTSAEDKGA QATVAALATA LTLRELRAHS |
| 6 | *Mycobacterium canettii* | >tr\|A0A8I0EMM5\|A0A8I0EMM5_9MYCO OS=Mycobacterium canettii OX=78331 GN=ribH PE=3 SV=1<br>MTGGAGVPDL PSLDASGVRL AIVASSWHGK ICDALLDGAR NVAAGCGLDD PTVVRVLGAI EIPVVAQELA RNHDAVVALG VVIRGQTPHF DYVCDAVTQG LTRVSLDSST PIGNGVLTTN TEEQALDRAG LPTSAEDKGA QATVAALATA LTLRELRAQP |
| 7 | *Mycobacterium orygis* | >tr\|A0A829C689\|A0A829C689_9MYCO OS=Mycobacterium orygis 112400015 OX=1305739 GN=ribH PE=3 SV=1<br>MPDLPSLDAS GVRLAIVASS WHGKICDALL DGARKVAAGC GLDDPTVVRV LGAIEIPVVA QELARNHDAV VALGVVIRGQ TPHFDYVCDA VTQGLTRVSL DSSTPIANGV LTTNTEEQAL DRAGLPTSAE DKGAQATVAA LATALTLREL RAHS |
| 8 | *Mycobacterium tuberculosis variant pinnipedii* | >tr\|A0A328GNP2\|A0A328GNP2_MYCTX OS=Mycobacterium tuberculosis variant pinnipedii OX=194542 GN=ribH PE=3 SV=1<br>MKGGAGVPDL PSLDASGVRL AIVASSWHGK ICDALLDGAR KVAAGCGLDD PTVVRVLGAI EIPVVAQELA RNHDAVVALG VVIRGQTPHF DYVCDAVTQG LTRVSLDSST PIANGVLTTN TEEQALDRAG LPTSAEDKGA QATVAALATA LTLRELRAHS |
| 9 | *Mycobacterium avium* | >sp\|A0QI08\|RISB_MYCA1 OS=Mycobacterium avium (strain 104) OX=243243 GN=ribH PE=3 SV=1<br>MSPAAGVPEM PALDASGVRL GIVASTWHSR ICDALLAGAR KVAADSGVEN PTVVRVLGAI EIPVVAQELA RNHDAVVALG VVIRGQTPHF EYVCDAVTQG ITRVSLDAST PVANGVLTTD NEQQALDRAG LPDSAEDKGA QAAGAALSAA LTLRELRARS |
| 10 | *Mycobacterium kansasii* | >tr\|X7Y0E1\|X7Y0E1_MYCKA OS=Mycobacterium kansasii 824 OX=1299328 GN=ribH PE=3 SV=1<br>MSGGAGVPDI PALDASGVRL AIVASTWHSQ ICDALLAGAR KVAADSGIDN PTVVRVLGAI EIPVVAQELA RQHDAVVALG VVIRGETPHF DYVCDAVTQG LTRVSLDEST PVANGVLTTN TEEQALDRAG LPTSAEDKGA QATAAALTTA LTLRELRVQS |
| 11 | *Mycobacteroides chelonae* | >tr\|A0A1S1LMU2\|A0A1S1LMU2_MYCCH OS=Mycobacteroides chelonae OX=1774 GN=ribH PE=3 SV=1<br>MSGEGIPSLT VGDASSLSLA IVASTWHDQI CTALLEGAKR VAADAGIDRP TVVRVLGAIE IPVVAQALAR THDAVVALGV VIQGETPHFN YVCDAVTTGL TRVSLDTSTP VANGVLTVNN EQQALDRAGL PESSEDKGAQ AAAAALDTAL TLRRLRQPWT EA |
| 12 | *Mycolicibacterium fortuitum* | >tr\|A0A1A0TGF6\|A0A1A0TGF6_MYCFO OS=Mycolicibacterium fortuitum OX=1766 GN=ribH PE=3 SV=1<br>MSAHGVPDLP QVDASNVKLA IVASTWHTQI CDALLDGARK VAADAGIAEP TVVRVLGAIE |

| | | |
|---|---|---|
| | | IPVVAQALAA THDAVVALGV VIRGQTPHFD YVCDAVTQGL TRVSLDASTP VANGVLTTDN EAQALDRAGL PDSTEDKGAQ AAAAALSTAL TLRELRSKA |
| 13 | *Mycobacterium intracellulare* | >tr\|X8AJW1\|X8AJW1_MYCIT OS=Mycobacterium intracellulare OX=1767 GN=ribH PE=3 SV=1<br>MSPAEGVPEV PPLDASGLRL ALVASTWHSE ICDALLAGAS KVASESGVDD PTVVRVIGAI EIPVVAQELA RNHDAVVALG VVIRGQTPHF EYVCDAVTQG LTRVSLDTST PVANGVLTTD TEQQALDRAG LPESAEDKGA QATLAALTTA LTLRELRARS |
| 14 | *Mycobacterium malmoense* | >tr\|A0A1S2WFZ2\|A0A1S2WFZ2_MYCMA OS=Mycobacterium malmoense OX=1780 GN=ribH PE=3 SV=1<br>MSGGAGVPEI PALDASGLRL GIVASTWHGQ ICDALLAGAR KMAAESGVEN PTVVRVLGAI EIPVVAQQLA RDHDAVVALG VVIRGETPHF DYVCDSVTQG LTRVSLDACT PIGNGVLTTN TEEQALARAG LPASAEDKGA QAAGAALTAA LTLRDLRARS |
| 15 | *Mycobacterium simiae* | >tr\|A0A5B1BKW0\|A0A5B1BKW0_MYCSI OS=Mycobacterium simiae OX=1784 GN=ribH PE=3 SV=1<br>MSGGAGVPDI PALDASGVRL AIVASTWHAE ICDALLAGAR KVATESGIDN PTVARVLGAI EIPVLAQELA RSHDAVVALG VVIRGETPHF DYVCDAVTQG LTRVSLDAST PVANGVLTTN TEEQARDRAG LPTSAEDKGA QATAAALTTA LALRELRAQS |
| 16 | *Mycobacterium xenopi* | >tr\|A0A2X1TE33\|A0A2X1TE33_MYCXE OS=Mycobacterium xenopi OX=1789 GN=ribH PE=3 SV=1<br>MSPAAGMPEL PALDASGLKL AIVASTWHTE ICDALLTGAR KTAAEWGIDD PTVVRVLGAI EIPVVAQELA RSHDAVVALG VVIRGETPHF DYVCDAVTQG LTRVSLDAST PVANGVLTTN TEEQARDRAG LPTSTEDKGA QATAAALNTA LTLRELRTRS |
| 17 | *Mycobacterium gordonae* | >tr\|A0A0Q2RQN3\|A0A0Q2RQN3_MYCGO OS=Mycobacterium gordonae OX=1778 GN=ribH PE=3 SV=1<br>MSGGAGVPEM PVLDASGLRL AIVASTWHDK ICDALLAGAR KVATDSGIDT PTVVRVLGAI EIPVVAQELA RNHDAVVALG VVIRGETPHF DYVCDAVTQG LTRVSLDAST PVANGVLTTN TEAQALDRAG LPSSAEDKGA QAAAAALTTA LTLRDLRAQS |
| 18 | *Mycobacterium celatum* | >tr\|A0A1X1RSX5\|A0A1X1RSX5_MYCCE OS=Mycobacterium celatum OX=28045 GN=ribH PE=3 SV=1<br>MSPAAGVPDM PALDASSVKL AIVASTWHTT ICDALLGGAR KTAAEWGVDD PTVVRVLGAI EIPVVAQALA REHDAVVALG VVIRGQTPHF DYVCDAVTQG LTRVSLDAST PVANGVLTTN TEEQALDRAG LPDSAEDKGA QATAAALSTA LTLRDLRAKP |
| 19 | *Mycobacterium haemophilum* | >tr\|A0A0I9TYM3\|A0A0I9TYM3_9MYCO OS=Mycobacterium haemophilum OX=29311 GN=ribH PE=3 SV=1<br>MSGGAGVPEV PAIDASGLRL GIVASTWHGR ICDALLAGAR KVAANSGVDN PTVVRVLGAI EIPVVAQELA RTHDAVVALG VVIRGATPHF DHVCNSVTQG LTRVALDTST PVGNGVLTTN TEEQALDRAG LPTSAEDKGA QAAAAALTTA LTLLNLRART |
| 20 | *Mycobacterium ulcerans* | >sp\|A0PPL7\|RISB_MYCUA OS=Mycobacterium ulcerans (strain Agy99) OX=362242 GN=ribH PE=3 SV=1<br>MSGGAGIPDV PAFDASGVRL AIVASTWHTK ICDALLAGAR NTAADSGIDN PTVVRVLGAI EIPVVAQELT RNHDAVVALG VVIRGETPHF DYVCDVVTQG LTRVSLDSST PVANGVLTTN SEEQALNRAG LPTSDEDKGA QATAAALTTA LTLRELRAES |
| 21 | *Mycobacterium marinum* | >sp\|B2HP68\|RISB_MYCMM OS=Mycobacterium marinum (strain ATCC BAA-535 / M) OX=216594 GN=ribH PE=3 SV=1<br>MSGGAGIPDV PAFDASDVRL AIVASTWHTK ICDALLAGAR NTAADSGIDN PTVVRVLGAI EIPVVAQELT RNHDAVVALG VVIRGETPHF DYVCDAVTQG LTRVSLDSST PVANGVLTTN SEEQALNRAG LPTSDEDKGA QATAAALTTA LTLRELRAEA |
| 22 | *Mycobacterium nonchromogen icum* | >tr\|A0A1X1ZI26\|A0A1X1ZI26_MYCNO OS=Mycobacterium nonchromogenicum OX=1782 GN=ribH PE=3 SV=1<br>MSPAAGVPDM PALDASDVRL AIVASTWHTR ICDALLAGAR RVAAEAGIPE PTVVRVLGAI EIPVVTQEVA RNHDAVVALG VVIRGATPHF DYVCDAVTQG LTRVSLDAAT PVANGVLTVN DEEQALDRAG LPGSTEDKGA QAAAAALSTA LTLRELRAAP |
| 23 | *Mycobacterium parascrofulace um* | >tr\|D5PHU0\|D5PHU0_9MYCO OS=Mycobacterium parascrofulaceum ATCC BAA-614 OX=525368 GN=ribH PE=3 SV=1<br>MEPCDRRRQD DAVRVSPAWG TAQVSGSGEP EIPALDATGL RLGIVASTWH GKICEALLQG<br>ARRVAAESGI DNPTVVRVLG AIEIPVVAQE LTRNHDAVIA LGVVIRGETP HFSYVCDAVT QGLTRVALDA STPVANGVLT TNTEQQALDR AGLPESVEDK GAQAAAAALT AALTLRDLRA RP |
| 24 | *Mycobacterium szulgai* | >tr\|A0A1X2E6B9\|A0A1X2E6B9_MYCSZ OS=Mycobacterium szulgai OX=1787 GN=ribH PE=3 SV=1<br>MSGGAGVPEM PVLDAADVQL AIVASTWHTE VCDALLAGAL KVAADSGIPS PTVVRVLGAI |

| | | |
|---|---|---|
| | | EIPVVVQELA RSHDAVVALG VVIRGETPHF DYVCDAVTQG LTRVSLDEST PVGNGVLTTN TEEQALDRAG LPTSAEDKGA QATAAALATA LTLRDLRAQS |
| 25 | *Mycobacterium scrofulaceum* | >tr\|A0A1A2TYK3\|A0A1A2TYK3_MYCSC OS=Mycobacterium scrofulaceum OX=1783 GN=ribH PE=3 SV=1<br>MSGSGEPDIP ALDASGLRVG IVASTWHSKI CEALLDGARR VAAESGIDNP TVVRVLGAIE LPVVAQELAR NHDAVIALGV VIRGGTPHFE YVCDAVTQGL TRVALDASTP VANGVLTTNT EQQALDRAGL PESVEDKGAQ AAGAALTAAL TLRELRARS |
| 26 | *Mycobacterium tuberculosis variant microti* | >PLV44826.1 6,7-dimethyl-8-ribityllumazine synthase [Mycobacterium tuberculosis variant microti OV254]<br>MSGAGIPDLGELDASGLRLGIVASTWHSTICDALLAGAERVVARSGVDNPTVVRVHGAIEIPV VAQELAR<br>NHDAVIALGVVIRGGTPHFEYVCDAVTQGLTRVSLDASTPVANGVLTTNTEEQALDRAGLPT SAEDKGAQAAGAALTAALALRDLRARS |
| 27 | *Mycobacterium terrae* | >BBX22440.1 6,7-dimethyl-8-ribityllumazine synthase [Mycolicibacter terrae]<br>MSGTGVPDMPALDASGVRLAIVASTWHTRICDALLAGARKVAADAGVADPTVVRVLGAIEIP VVAQEVAR<br>NHDAVVALGVVIRGGTPHFDYVCDAVTQGLTRVSLDAATPVANGVLTVNDEQQALDRAGLP GSAEDKGAQAAAAALSTALTLRDLRAAL |
| 28 | *Mycobacterium tuberculosis variant caprae* | >PRH94793.1 6,7-dimethyl-8-ribityllumazine synthase [Mycobacterium tuberculosis variant caprae]<br>MKGGAGVPDLPSLDASGVRLAIVASSWHGKICDALLDGARKVAAGCGLDDPTVVRVLGAIEI PVVAQELA<br>RNHDAVVALGVVIRGQTPHFDYVCDAVTQGLTRVSLDSSTPIANGVLTTNTEEQALDRAGLP TSAEDKGA<br>QATVAALATALTLRELRAHS |
| 29 | *M. leprae* | >[M. leprae]<br>MSGGAGIPEVPGIDASGLRLGIVASTWHSRICDALLAGARKVAADSGIDGPTVVRVLGAI<br>EIPVVVQELARHHDAVVALGVVIRGDTPHFDYVCNSVTQGLTRIALDTSTPVGNGVLTTN<br>TEKQALDRAGLPTSAEDKGAQAAAAALTTALTLLNLRSRI |

**Sequence alignment of M. tb *H37Rv* RibH protein with that of *M. leprae***

| Score | Expect | Method | Identities | Positives | Gaps |
|---|---|---|---|---|---|
| 239 bits(610) | 5e-84 | Compositional matrix adjust. | 125/158(79%) | 142/158(89%) | 0/158(0%) |

```
Query  1    MKGGAGVPDLPSLDASGVRLAIVASSWHGKICDALLDGARKVAAGCGLDDPTVVRVLGAI
60
            M GGAG+P++P +DASG+RL IVAS+WH +ICDALL GARKVAA  G+D PTVVRVLGAI
Sbjct  1    MSGGAGIPEVPGIDASGLRLGIVASTWHSRICDALLAGARKVAADSGIDGPTVVRVLGAI
60

Query  61   EIPVVAQELARNHDAVVALGVVIRGQTPHFDYVCDAVTQGLTRVSLDSSTPIANGVLTTN
120
            EIPVV QELAR+HDAVVALGVVIRG-TPHFDYVC++VTQGLTR++LD+STP+-NGVLTTN
Sbjct  61   EIPVVVQELARHHDAVVALGVVIRGDTPHFDYVCNSVTQGLTRIALDTSTPVGNGVLTTN
120

Query  121  TEEQALDRAGLPTSAEDKGAQATVAALATALTLRELRA   158
            TE+QALDRAGLPTSAEDKGAQA——AAL-TALTL--LR+
Sbjct  121  TEKQALDRAGLPTSAEDKGAQAAAAALTTALTLLNLRS   158
```

**Supplementary Table 3. Conservation in drug binding pocket in Rib H across mycobacterial species.** Summary of results obtained from sequence alignment of nucleotide and protein sequences of lumazine synthase (RibH) from different mycobacterial species. Chances of whether the drug would bind to the protein was assumed to be yes if the % of similarity in drug binding pocket was found to be above 90%.

| Serial No. | Organism | Nature of Mycobacterium (MTC/) NTM) | Nucleotide sequence accession ID | % Identity in Nucleotide sequence | Protein Accesion ID (Uniprot/ UniParc) | % Identity in Protein sequence | % Similarity in Protein sequence | No of residues identical in drug binding pocket (out of 22 residues) | % Identity in the amino acid in the drug binding pocket | No of residues identical in co-crystal ligand binding pocket (out of 29 residues) | % Identity in the amino acid in the co-crystal ligand binding pocket | Chance that it will bind to drug (Y/N) |
|---|---|---|---|---|---|---|---|---|---|---|---|---|
| 1 | *M. abscessus* | NTM | CP012044.1 | 75 | B1MCA3 | 72.7 | 81.4 | 22 | 100 | 26 | 89.6 | Y |
| 2 | *M. africanum* | MTC | CP014617.1 | 100 | A0A120J0T0 | 100 | 100 | 22 | 100 | 29 | 100 | Y |
| 3 | *M. avium* | NTM | AP020326.1 | 81.36 | A0QI08 | 81.9 | 91.2 | 22 | 100 | 28 | 96.5 | Y |
| 4 | *M. bovis* | MTC | LR699570.1 | 100 | P66035 | 100 | 100 | 22 | 100 | 29 | 100 | Y |
| 5 | *M. bovis BCG* | MTC | CP033311.1 | 100 | A0A0K2HV99 | 100 | 100 | 22 | 100 | 29 | 100 | Y |
| 6 | *M. canettii* | MTC | FO203507.1 | 99.59 | A0A8I0EMM5 | 96.9 | 96.9 | 21 | 95.45 | 28 | 96.5 | Y |
| 7 | *M. caprae* | MTC | CP016401.1 | 100 | PRH94793.1 | 100 | 100 | 22 | 100 | 29 | 100 | Y |
| 8 | *M. celatum* | NTM | NA | NA | A0A1X1RSX5 | 84.4 | 90 | 22 | 100 | 28 | 96.5 | Y |
| 9 | *M. chelonae* | NTM | CP058976.1 | 75.97 | A0A1S1LMU2 | 71.8 | 81 | 22 | 100 | 26 | 89.6 | Y |
| 10 | *M. fortuitum* | NTM | CP011269.1 | 76.97 | A0A1A0TGF6 | 79.4 | 86.9 | 22 | 100 | 28 | 96.5 | Y |
| 11 | *M. gordonae* | NTM | CP070973.1 | 79.04 | A0A0Q2RQN3 | 85.6 | 91.9 | 22 | 100 | 27 | 93.1 | Y |
| 12 | *M. haemophilum* | NTM | CP011883.2 | 80 | A0A0I9TYM3 | 81.2 | 91.2 | 21 | 95.45 | 26 | 89.6 | Y |
| 13 | *M. intracellulare* | NTM | CP023151.1 | 81.62 | X8AJW1 | 82.5 | 91.9 | 21 | 95.45 | 27 | 93.1 | Y |
| 14 | *M. kansasii* | NTM | CP089231.1 | 82.4 | X7Y0E1 | 88.1 | 93.1 | 22 | 100 | 27 | 93.1 | Y |
| 15 | *M. malmoense* | NTM | CP060015.1 | 82.64 | A0A1S2WFZ2 | 81.2 | 91.2 | 21 | 95.45 | 28 | 96.5 | Y |
| 16 | *M. marinum* | NTM | CP118910.1 | 78.69 | B2HP68 | 84.4 | 91.2 | 22 | 100 | 27 | 93.1 | Y |
| 17 | *M. microti* | MTC | CP010333.1 | 100 | PLV44826.1 | 80 | 86.9 | 21 | 95.45 | 26 | 89.6 | Y |
| 18 | *M. nonchromogenicum* | NTM | NA | NA | A0A1X1ZI26 | 80.6 | 88.1 | 22 | 100 | 27 | 93.1 | Y |
| 19 | *M. orygis* | MTC | CP063804.2 | 100 | A0A829C689 | 95.6 | 96.2 | 22 | 100 | 29 | 100 | Y |
| 20 | *M. parascrofulaceum* | NTM | JF271799.1 | 78.05 | D5PHU0 | 68.1 | 78 | 22 | 100 | 27 | 93.1 | Y |
| 21 | *M. pinnipedii* | MTC | NZ_MWXB01000092 | 100 | A0A328GNP2 | 100 | 100 | 22 | 100 | 29 | 100 | Y |

| | | | | | | | | | | | | |
|---|---|---|---|---|---|---|---|---|---|---|---|---|
| 22 | *M. scrofulaceum* | NTM | NA | NA | A0A1A2TYK3 | 79.4 | 90 | 22 | 100 | 27 | 93.1 | Y |
| 27 | *M. simiae* | NTM | AP022568.1 | 78.08 | A0A5B1BKW0 | 85.6 | 92.5 | 22 | 100 | 29 | 100 | Y |
| 24 | *M. szulgai* | NTM | NA | NA | A0A1X2E6B9 | 83.1 | 90.6 | 21 | 95.45 | 26 | 89.6 | Y |
| 25 | *M. terrae* | NTM | AP022564.1 | 80 | BBX22440.1 | 81.9 | 89.4 | 22 | 100 | 27 | 93.1 | Y |
| 26 | *M. ulcerans* | NTM | AP017624.1 | 78.62 | A0PPL7 | 85 | 91.2 | 22 | 100 | 27 | 93.1 | Y |
| 27 | *M. xenopi* | NTM | AP022314.1 | 79.45 | A0A2X1TE33 | 83.8 | 91.2 | 22 | 100 | 27 | 93.1 | Y |
| 28 | *M. leprae* | MLC | ML0560* | ns | Q9CCP3 | 79 | 89 | 21 | 95.45 | 26 | 89.6 | Y |

***Mycobacterium tuberculosis complex*** **(MTC); Non-tuberculous Mycobacterium (NTM);** ***Mycobacterium leprae complex*** **(MLC); NA: Not available in web resources, ns: non-significant**
* For *M. leprae* Nucleotide and protein sequence source is kegg, Uniprot, respectively.

**Supplementary Table 4. List of bacterial strains used in this study**

| Bacterial Strain | Description | Reference |
|---|---|---|
| *M. tb* strain *H37Rv* | Obtained from ATCC (ATCC27294 strain), TMC 102 [H37Rv] | https://www.atcc.org/products/27294 |
| *M. tb H37Rv* $mc^2$ 7902 | Auxotroph strain (*H37Rv* Δ*pan*CD ΔleuCD ΔargB) | Vilchèze C, Copeland J, Keiser TL, Weisbrod T, Washington J, Jain P, Malek A, Weinrick B, Jacobs Jr WR. Rational design of biosafety level 2-approved, multidrug-resistant strains of Mycobacterium tuberculosis through nutrient auxotrophy. MBio. 2018 Jul 5;9(3):10-128. |
| *M. tb ribH-KD* | Auxotroph strain (H37Rv Δ*pan*CD Δ*leu*CD Δ*arg*B), *ribH* conditional gene silencing achieved by ATc induction. | This study |
| *E.coli* DH5-α | Obtained from NEB (T1 phage resistant and *endA* deficient) | https://international.neb.com/products/c2987-neb-5-alpha-competent-e-coli-high-efficiency#Product%20Information |
| *E.coli* BL21 (DE3) | *E. coli* strain, suitable for transformation and protein expression from plasmids with T7 promoter. | https://international.neb.com/products/c2527-bl21de3-competent-e-coli#Product%20Information |

**Supplementary Table 5. List of primers used in the study.**

| Primer name | Gene number and description | Sequence (5'-3') | Size of primer (bp) | Experiments |
|---|---|---|---|---|
| *ribH*_RT_F | *Rv1416, ribH 6,7-dimethyl-8-ribityllumazine synthase (lumazine synthase)* | CAATCATGATGCCGTCGT | 18 | For RT-PCR analysis |
| *ribH*_RT_R | | GGGTCAGTCCCTGGGTTAC | 19 | |
| *SigA*_RT_F | Rv2703, *sigA* RNA polymerase sigma factor SigA (sigma-A) | ATCGCTGAACCCACCGAAAA | 20 | For RT-PCR analysis |
| *SigA*_RT_R | | GACCTCTTCCTCGGCGTTG | 19 | |
| *ribH*_F_P | Rv1416, *ribH 6,7-dimethyl-8-ribityllumazine synthase (lumazine synthase)* | GGGGCATATGAAGGGTGGCG CCGGGGT | 27 | PCR amplification for cloning of *ribH* in pet28a |
| *ribH*_R_P | | GGGGCTCGAGTAGTCACGAGT GAGCGCGCAGCT | 33 | |
| Cr_UP | *Rv1416*, *ribH 6,7-dimethyl-8-ribityllumazine synthase (lumazine synthase)* | GATCTTTCCGTGCCAGCTG | 19 | Guide sequence targeting *ribH* |
| Cr_DN | | CGCAGCTGGCACGGAAAGATC CATG | 25 | |
| *Kan*$^r$ F | *Kan*$^r$ Kanamycin resistance gene cassette | GAGAAAACTCACCGAGGCAG | 20 | For screening clones carrying *kan*$^r$ plasmid |
| *Kan*$^r$ F | | GTATTTCGTCTCGCTCAGGC | 20 | |
| *Hyg*$^r$ F | *Hyg*$^r$ Hygromycin resistance gene cassette | CCGGGCTCGCAGCAGCGGGC | 20 | For screening clones carrying *hyg*$^r$ plasmid |
| *Hyg*$^r$ R | | CCTCGAACACCTCGAAGTCG | 20 | |

*Rv- notation is used for gene numbers in M. tuberculosis H37Rv*

#### Supplementary Methods

#### Synthesis of compounds

##### Supplementary Table 6. Molecular weight, SMILES ID and structure of A10, A39 and A52

| Sl. No. | Data base Code | Tube Code | Mol. weight | Simplified molecular-input line-entry system (SMILES) ID | Structure |
|---|---|---|---|---|---|
| 1 | NR-310 | A10 | 315.30 | O=C(/C(S/1)=C\C2=CC=C([N+]([O-])=O)O2)NC1=N\C3=CC=CC=C3 | 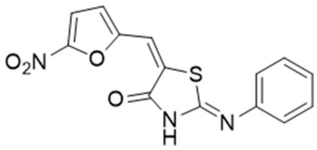 |
| 2 | NR-340 | A39 | 349.75 | ClC1=CC=C(/N=C2NC(/C(S/2)=C\C3=CC=C([N+]([O-])=O)O3)=O)C=C1 | 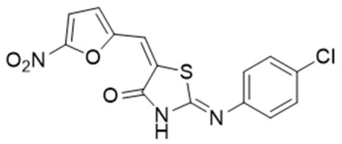 |
| 3 | NR 353 | A52 | 329.33 | CC1=CC=C(/N=C2NC(/C(S/2)=C\C3=CC=C([N+]([O-])=O)O3)=O)C=C1 | 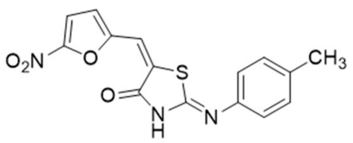 |

##### Synthesis scheme of 2-(phenylimino) thiazolidin-4-one derivatives

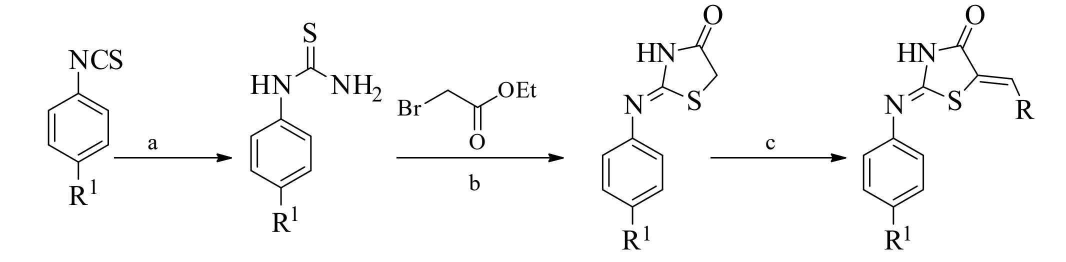

**TZ_01a-e** **TZ_02a-e** **TZ_03a-e** **TZ_04-TZ_56**

$R^1 = OCH_3, CH_3, H, Cl, NO_2$

Reagents & Conditions: a: $NH_3$, THF, rt; b: anhy NaOAc, EtOH, 60 °C; c: RCHO, piperidine, EtOH, 60 °C, 12 h.

The target molecules were synthesized by following the three-step synthetic protocol. Synthesis started with the commercially available various substituted aryl isothiocyanates on reaction with ammonia solution in THF and the resulting solid was filtered and washed with excess of water, cold ethanol and diethyl ether and dried to afford the corresponding substituted arylthioureas (**TZ_02a-e**). Further the cyclization reaction was carried out by the reaction of (**TZ_02a-e**) with ethyl 2-bromoacetate and anhydrous NaOAc in absolute ethanol at 60 °C to afford the key intermediate 2-(substituted aryimino) thiazolidin-4-one (**TZ_03a-e**) in good yields. These reactions were also successfully carried out using ethyl-2-chloroacetate, anhydrous NaOAc in absolute ethanol at 60 °C but the former reaction conditions resulted in good yields. In final step we used Knoevenagal condensation of the compound **TZ_03a-e** with various substituted aldehydes using piperidine in absolute ethanol at 60 °C, upon complete consumption of the starting material the reaction mixture was filtered to remove the bromide salts and the filtrate was concentrated and obtained residue was re-dissolved in ethyl acetate and washed with water and brine solution and purified to produce title compounds **TZ_04-TZ_56.**

**General procedure for the synthesis of (TZ_02a-e)**

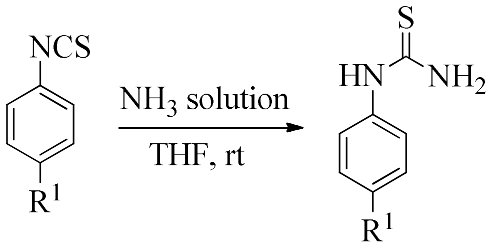


**TZ_01a-e** **TZ_02a-e**

$R_1$ = H, $CH_3$, $OCH_3$
Cl, $NO_2$

To the starting material **TZ_01a-e** (1.0 equiv) in THF, was added $NH_3$ solution (10 vol) under cooling conditions and allowed the reaction mixture to stir at room temperature for 3 h, then the solids formed in the reaction mixture was filtered and washed with diethyl ether and dried to afford the pure product **(TZ_02a-e)** as white solid (yields 90-95%).

**1-Phenylthiourea (TZ_02a)**

Following the general procedure, the product was synthesized from phenyl isothiocyanate **TZ_01a** (3.00 g, 22.22 mmol), $NH_3$ solution (30 mL) produced 1-phenylthiourea **(TZ_02a)** (3.2 g, 98%) as white solid. ESI-MS showed 153 $[M+H]^+$ and carried to next step.

**1-(*p*-Tolyl) thiourea (TZ_02b)**

Following the general procedure the product was synthesized from p-tolyl isothiocyanate **TZ_01b** (5.00 g, 33.55 mmol), $NH_3$ solution (50 mL) produced 1-(p-tolyl) thiourea **(TZ_02b**) (5.4 g, 98%) as white solid. ESI-MS showed 167 $[M+H]^+$ and carried to next step.

**1-(4-Chlorophenyl)thiourea (TZ_02d)**

Following the general procedure the product was synthesized from 4-chlorophenylisothio cyanate **TZ_01d** (5.00 g, 29.58 mmol), $NH_3$ solution (50 mL) produced 1-(4-chlorophenyl) thiourea **(TZ_02d**) (5.3 g, 96%) as white solid. ESI-MS showed 187 $[M+H]^+$ and carried to next step.

**General procedure for the synthesis of (TZ_03a-e)**

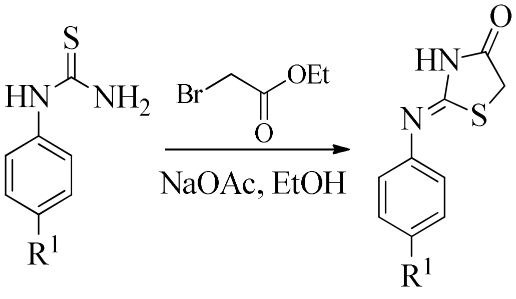


**TZ_02a-e** **TZ_03a-e**

To the stirred solution of **TZ_02a-e** (1.0 equiv) in EtOH (10 vol) was added anhydrous NaOAc (5.0 equivalents) followed by ethyl bromoacetate (2.0 equivalents) and the reaction mixture was heated at 60 °C for 7 h. The solids formed in the reaction mixture were filtered and washed with EtOH, the filtrate was concentrated and the solid obtained was partitioned between ethyl acetate and water then washed with brine solution, the Ethyl acetate layer was dried over anhydrous $Na_2SO_4$ and concentrated under vacuo then the solid obtained was triturated with $CH_2Cl_2$/hexanes the resulting solid was filtered, washed with diethyl ether and dried to get **TZ_03a-e** in pure form (yields >80%) and used for the next step.

**2-(Phenylimino) thiazolidin-4-one (TZ_03a)**

Following the general procedure, the product was synthesized from 1-phenylthiourea **TZ_02a** (3.20 g, 21.05 mmol), anhydrous NaOAc (8.63 g, 105.05 mmol) and Ethyl bromoacetate (4.65 mL, 42.10 mmol) produced 2-(phenylimino) thiazolidin-4-one (**TZ_03a**) (3.6 g, 89%) as yellow solid. ESI-MS showed 193 [M+H]$^+$. $^1$H NMR (400 MHz, DMSO-$d_6$) δ 11.73 (s,0.5×1H), 11.15 (s,0.5×1H), 7.70 (m, 1H), 7.40–7.37 (m, 2H), 7.18–7.13 (m, 1H), 7.00 (m, 1H), 4.01 (s, 1H), 3.97 (s, 1H).

**2-(p-Tolylimino) thiazolidin-4-one (TZ_03b)**

Following the general procedure, the product was synthesized from 1-(p-tolyl) thiourea **(TZ_02b)** (5.40 g, 32.53 mmol), anhydrous NaOAc (13.33 g, 162.65 mmol) and ethylbromoacetate (7.19 mL, 65.06 mmol) produced 2-(p-tolylimino) thiazolidin-4-one **(TZ_03b)** (5.8 g, 86%) as yellow solid. ESI-MS showed 207 [M+H]$^+$. $^1$H NMR (400 MHz, DMSO-$d_6$) δ 11.33 (s, 1H), 7.56 (d, *J* = 7.4 Hz, 1H), 6.91 (d, *J* = 7.6 Hz, 1H), 3.98 (s, 1H), 3.91 (s, 1H), 2.28 (s, 3H).

**2-((4-Chlorophenyl) imino) thiazolidin-4-one (TZ_03d)**

Following the general procedure, the product was synthesized from 1-(4-chlorophenyl) thiourea

**TZ_02d** (5.30 g, 28.49 mmol), anhydrous NaOAc (11.68 g, 142.47 mmol) and ethyl bromoacetate (6.30 mL, 56.98 mmol) produced 2-((4-chlorophenyl) imino) thiazolidin-4-one **(TZ_03d)** (5.4 g, 84%) as a brown solid. ESI-MS showed 227 [M+H]$^+$. $^1$H NMR (300 MHz, $CDCl_3$) δ 9.01 (s, 1H), 7.54–7.46 (bm, 4H), 3.54 (s, 2H).

**General procedure for the synthesis of compounds TZ_04 – TZ_56**

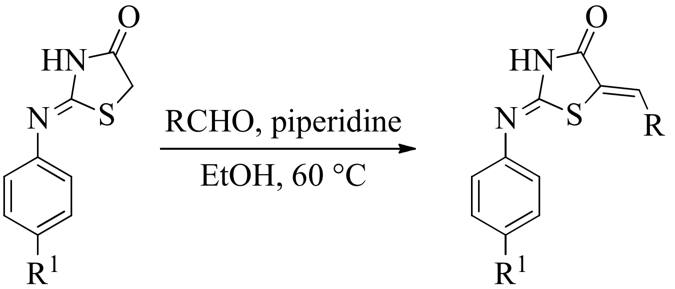


**TZ_03a-e** **TZ_04-TZ_56**

To the stirred solution of **TZ_03a-e** (1.0 equivalents) in EtOH, was added piperidine (1.0 equivalents) and RCHO (1.2 equivalents) and heated at 60°C for 12 h, then the solids formed in the reaction mixture were filtered and washed with EtOH, hexanes to afford the pure product in good yields.

**5-((5-Nitrofuran-2-yl) methylene)-2-(phenylimino) thiazolidin-4-one (TZ_13):** Yield: 65%; m.p. 241–242 °C; MS(ESI) *m/z* 316 [M+H]$^+$. $^1$H NMR (400 MHz, $CDCl_3$) δ 9.93 (s, 1H), 8.31 (d, *J* = 8.0

Hz, 1H), 7.72–7.63 (m, 2H), 7.58 (s, 1H), 7.54 (d, *J* = 8.0Hz, 1H), 7.49–7.32 (m, 3H); $^{13}$C NMR (100 MHz, $CDCl_3$) 176.3, 164.2, 156.0, 146.2, 135.2, 133.1, 132.0, 130.7(2C), 127.2, 125.9(2C), 124.2, 119.3. Anal calcd for $C_{14}H_9N_3O_4S$: C, 53.33; H, 2.88; N, 13.33% Found C, 53.42; H, 2.92; N, 13.45%.

**5-((5-Nitrofuran-2-yl) methylene)-2-(*p*-tolylimino) thiazolidin-4-one (TZ_24):** Yield: 61%; m.p. 229–230 °C; MS(ESI) *m/z* 330 $[M+H]^+$. $^1$H NMR (300 MHz, DMSO-$d_6$) δ 9.20 (s, 1H), 8.17 (d, *J* = 8.6 Hz, 1H), 7.70–7.61 (m, 4H), 7.55 (d, *J* = 8.0 Hz, 2H), 2.47 (s, 3H); $^{13}$C NMR (75 MHz, DMSO-$d_6$) 176.2, 164.1, 155.9, 146.2, 135.2, 133.0, 132.1, 130.8(2C), 127.1, 125.7(2C), 124.1, 119.1, 25.2. Anal calcd for $C_{15}H_{11}N_3O_4S$: C, 54.71; H, 3.37; N, 12.76% Found C, 54.77; H, 3.42; N, 12.81%.

**2-((4-Chlorophenyl) imino)-5-((5-nitrofuran-2-yl) methylene) thiazolidin-4-one (TZ_44):** Yield: 61%; m.p. 211–212 °C; MS(ESI) *m/z* 350 $[M+H]^+$. $^1$H NMR (300 MHz, DMSO-$d_6$) δ 9.20 (s, 1H), 8.11 (d, *J* = 8.8 Hz, 1H), 7.73–7.62 (m, 4H), 7.58 (d, *J* = 8.0 Hz, 2H); $^{13}$C NMR (75 MHz, DMSO-$d_6$) 178.4, 166.3, 156.7, 146.9, 136.1, 134.0, 132.7, 131.6(2C), 129.1, 126.6(2C), 124.8, 119.8. Anal calcd for $C_{14}H_8ClN_3O_4S$: C, 48.08; H, 2.31; N, 12.01% Found C, 48.14; H, 2.38; N, 12.06%.

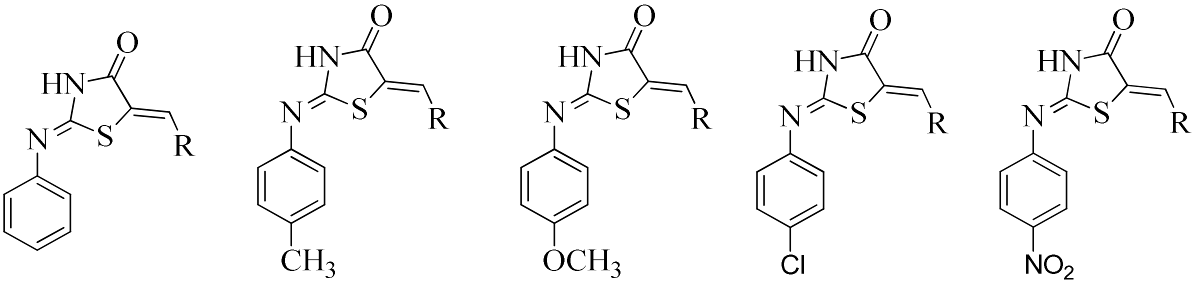


**TZ_04-TZ_14 TZ_15-TZ_25 TZ_26-TZ_34 TZ_35-TZ_45 TZ_46-TZ_56**

**Supplementary Table 7: R group description, yield, melting point (MP), molecular formula and weight of the shortlisted compounds**

| Compound | Compound Code | R | Yield (%) | M.P. (°C) | Molecular formula | Molecular weight |
|---|---|---|---|---|---|---|
| TZ_13 | A10 | 5-Nitro-2-furyl | 65 | 241-242 | $C_{14}H_9N_3O_4S$ | 315.30 |
| TZ_44 | A39 | 5-Nitro-2-furyl | 61 | 211-212 | $C_{14}H_8ClN_3O_4S$ | 349.75 |
| TZ_24 | A52 | 5-Nitro-2-furyl | 61 | 229-230 | $C_{15}H_{11}N_3O_4S$ | 329.33 |

#### Supplementary Table 8. ADME properties of A10, A39 and A52

| Compound Code | ILOGP | XLOGP3 | WLOGP | MLOGP | SILICOS-IT | Consensus LOG $P_{O/W}$ | Water solubility LOG S (ESOL) | Water solubility LOG S (Ali) | Water solubility LOG S (SILICOS-IT) | GI Absorption | BBB Permeant | Bioavailability Score | P-glycoprotein substrate | CYP inhibition [CYP1A2, CYP2C19, CYP2C9, CYP2D6, CYP3A4] |
|---|---|---|---|---|---|---|---|---|---|---|---|---|---|---|
| A10 | 1.89 | 2.47 | 2.59 | 1.43 | 1.16 | 1.91 | -3.52 (Soluble) | -4.75 (moderately soluble) | -4.43 (moderately soluble) | High | No | 0.55 | No | **CYP1A2 (Y), CYP2C19 (Y), CYP2C9 (Y), CYP2D6 (N), CYP3A4 (N)** |
| A39 | 2.16 | 3.10 | 3.24 | 1.95 | 1.80 | 2.45 | -4.12 (moderately soluble) | -5.41 (moderately soluble) | -5.03 (moderately soluble) | High | No | 0.55 | No | **CYP1A2 (Y), CYP2C19 (Y), CYP2C9 (Y), CYP2D6 (N), CYP3A4 (N)** |
| A52 | 2.30 | 2.83 | 2.90 | 1.69 | 1.66 | 2.28 | -3.82 (Soluble) | -5.13 (moderately soluble) | -4.81 (moderately soluble) | High | No | 0.55 | No | **CYP1A2 (Y), CYP2C19 (Y), CYP2C9 (Y), CYP2D6 (N), CYP3A4 (N)** |

- Interpretation note: As calculated by SwissADME[1], all the compounds are predicted to have moderate to good solubility with high gastrointestinal (GI) absorption. The Lipophilicity is >1 and < 5, indicating good oral absorption. Bioavailability of 0.55 indicates these drugs to be good candidates for further preclinical evaluation. None of the drug molecules are substrate for P-glycoprotein induction hence, are going to be bioavailable and have good bio distribution. As only 3 out of 5 CYP450 family of enzymes are inhibited suggesting that drugs may have high intracellular retention, but can be metabolized by the residual 2 CYP450 enzymes and can possibly be cleared by the renal system. They are anticipated to have low drug-drug interaction and low toxicity. None of the compounds are predicted to cross BBB- Blood brain barrier.

**Cloning, expression and purification of RibH**

**Cloning:** In order to obtain purified lumazine synthase enzyme of *M. tuberculosis*, the gene encoding lumazine synthase *ribH* (*Rv1416*) was PCR amplified using *H37Rv* genomic DNA as template and oligonucleotides (5' GGGGCATATGAAGGGTGGCGCCGGGGT 3') as forward primer and (5'GGGGCTCGAGTAGTCACGAGTGAGCGCGCAGCT 3') as reverse primer. The amplified PCR product was digested using restriction enzymes *Nde* I and *Xho* I (New England Biolabs) and ligated using The Quick Ligation™ Kit (New England Biolabs) with the *E. coli* expression vector pET28(a) (Novagen) digested with the same set of restriction enzymes. The ligated plasmid DNA was transformed in to DH5α *E. coli* strain (New England Biolabs) and colonies were selected on 25 μg/ml kanamycin (Himedia). The clones were confirmed by PCR amplification, restriction digestion and sequencing **(Supplementary Figure 8)**. Following this, the sequence verified plasmid DNA was transformed into BL21(DE3) (New England Biolabs) and colonies were selected on 25 μg/ml kanamycin (Himedia) and confirmed by PCR amplification.

**Induction condition optimization:** The expression vector offers an inducible expression system wherein, protein of interest with a 6x-his tag at N-terminus is expressed under the control of T7 promoter and lac operator. Addition of optimised concentration of Isopropyl β-d-1-thiogalactopyranoside (IPTG) leads to expression of protein of interest. In order to optimise the expression of lumazine synthase (16 kDa), BL21(DE3) clones were grown in LB media with kanamycin with various concentrations of IPTG (0.2 – 0.8 mM) at various temperature conditions (20-37$^{o}$C) for 4 hours-overnight. The induction conditions offering maximum production of properly folded protein in soluble fraction (0.2 mM IPTG at 16° C overnight incubation) was utilised for final purification. The protein was purified by affinity chromatography using Ni-NTA resin using a gradient concentration of imidazole in the range of 100-300 mM as per manufacturer's specification (Clontech Takara Ni-NTA Resin_ 635660). The fraction giving the maximum purity (250 mM) was pooled and dialyzed against a 50 mM Potassium phosphate buffer (pH 7.0) containing 10% glycerol. The purified protein quality was assessed by various methods such as, sodium dodecyl sulphate polyacrylamide gel electrophoresis (SDS PAGE) **(Supplementary Figure 3)**, circular dichroism (CD) and dynamic light scattering (DLS) **(Supplementary Figure 4)**. The protein was flash frozen using liquid nitrogen followed by long term storage at -80$^{o}$C.

**Binding affinity determination using Microscale Thermophoresis (MST) assay**

To determine the binding affinity, MST assay was carried out using Monolith NT.115 microscale thermophoresis instrument (Nano Temper Technology). In this method, ligand and protein interaction is studied by employing Monolith His-tag labelling kit RED-tris-NTA 2$^{nd}$ Generation (Nano Temper Technology) as per the manufacturer's specification and as described previously[2]. This method relies on the high-affinity but non-covalent interaction of RED-tris-NTA fluorescent dye (containing flour RED - NT647) with six histidine-tag of the protein of interest. This conjugate provides specific binding and a high fluorescence signal with best signal-to-noise ratio even in complex samples such as cell lysates which often auto fluorescence in the blue and green part of the spectrum. When such a labelled protein is subjected to temperature gradient, it leads to thermophoresis, which is the directed motion of molecules in response to Infrared light induced temperature gradient. Thermophoresis depends on molecule size, charge, hydration shell and the extent of ligand binding as described[2,3].

***Protein labelling****:* Purified his-tagged RibH protein was diluted in 1X PBST (Phosphate buffered saline with 0.05% tween 20) at a final concentration of 200 nM. RED-tris-NTA fluorescent dye (5 µM) was diluted in 1X PBST at a final concentration of 100 nM. For labelling, equal volume of dye and protein (90 µl each) were mixed and incubated at room temperature for 30 minutes followed by centrifugation at 15000 g, 4 $^{o}$C for 10 minutes. The supernatant containing the labelled protein was then transferred to a fresh tube.

***Ligand preparation**:* For binding assay, riboflavin (positive control), A10, A39 and A52 compounds were dissolved in autoclaved milli Q water to a final concentration of 1 μM. Next, 1μM of the ligands were then two-fold serially diluted 16 times. 10 μl of 100 nM labelled protein was then mixed with 10 μl of serially diluted compounds (1000 nM-0.03 nM).

***Protein and Ligand binding**:* The final concentration of 50 nM protein was titrated against the final ligand concentrations of 500 nM-0.015 nM in 16 different capillary tubes.

***MST analysis:*** MST experiment was run for 20 seconds using Monolith NT.115 microscale thermophoresis instrument at room temperature (23° to 25°C) using an inbuilt software called Monolith NT Control Software. Capillary scan images were collected and NT Analysis Software was then used for a precise analysis of Microscale Thermophoresis data and quantification of dissociation constant ($K_d$) as per the manufacturer's instructions (Nano Temper Technology, https://www.uni-hohenheim.de/fileadmin/einrichtungen/mst/Manual_NT115.pdf)

**Computational Pipeline to determine the conserved nature of drug binding pocket in RibH**

Nucleotide sequences for the lumazine synthase (RibH) gene were obtained from NCBI's nucleotide and genome databases for various species of the Mycobacterium tuberculosis complex as well as non-tuberculous mycobacteria. The protein sequences for RibH were obtained from UniProt. Nucleotide sequence alignment for various species were carried out against the *rib H* (*Rv 1416*) sequence of *Mycobacterium tuberculosis* (strain ATCC 25618 / *H37Rv*), using the BlastN suite of NCBI to obtain % similarity and % identity. We then carried out multiple sequence alignment of the RibH protein sequence using CLUSTAL O and T-Coffee.

In order to determine % similarity and % identity in protein sequences, EMBOSS NEEDLE was used to compare the sequence from various species against *M.*

*tuberculosis* (strain ATCC 25618 / H37Rv). We then constructed a phylogenetic tree based on RibH protein sequence using T-Coffee. For the organisms, for which protein sequences were unavailable on UniProt, Blast X was used to find the protein sequences from GenBank.

In order to find if the drugs discovered by us could potentially be used to treat other mycobacterial infections, we manually analysed the aligned protein sequences of RibH from various organisms to find the identical residues within the drug binding and co-crystal ligand binding pockets. The drug binding pocket and the residues present thereof within the pocket were obtained using the Maestro suite of Schrodinger, and the residues present in the co-crystal ligand binding pocket were identified from PDBSum. The % identity of the residues in both pockets was tabulated in Supplementary Table 4.

**Pipeline of computational studies**

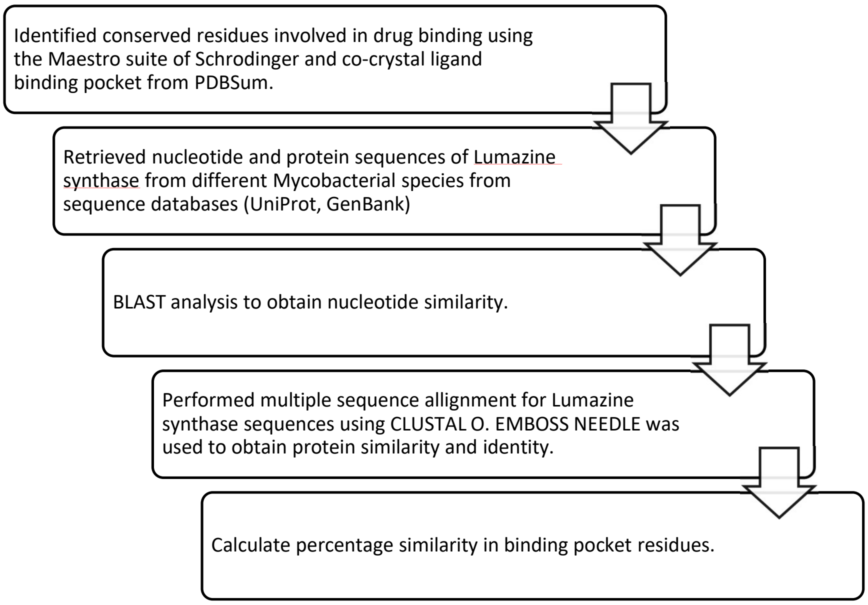


**Web resources used**
UniProt - https://www.uniprot.org/
NCBI - https://www.ncbi.nlm.nih.gov/
GenBank – https://www.ncbi.nlm.nih.gov/genbank/
PDBSUM – http://www.ebi.ac.uk/thornton-srv/databases/pdbsum/
MAESTRO (Schrodinger) - https://www.schrodinger.com/products/maestro
BLASTN – https://www.ncbi.nlm.nih.gov/geo/query/blast.html
BLASTX – https://blast.ncbi.nlm.nih.gov/Blast.cgi?PROGRAM=blastx&PAGE_TYPE=BlastSearch&LINK_LOC=blasthome
T-Coffee – https://www.ebi.ac.uk/Tools/msa/tcoffee/
EMBOSS NEEDLE – https://www.ebi.ac.uk/Tools/psa/emboss_needle/
